# Integrative spatial analysis reveals a multi-layered organization of glioblastoma

**DOI:** 10.1101/2023.07.06.547924

**Authors:** Alissa C. Greenwald, Noam Galili Darnell, Rouven Hoefflin, Dor Simkin, L. Nicolas Gonzalez-Castro, Christopher Mount, Dana Hirsch, Masashi Nomura, Tom Talpir, Merav Kedmi, Inna Goliand, Gioele Medici, Baoguo Li, Hadas Keren-Shaul, Michael Weller, Yoseph Addadi, Marian C. Neidert, Mario L. Suvá, Itay Tirosh

## Abstract

Glioma contains malignant cells in diverse states. Hypoxic regions are associated with a unique histology of pseudopalisading cells, while other regions appear to have limited histological organization, reflecting the diffuse nature of glioma cells. Here, we combine spatial transcriptomics with spatial proteomics and novel computational approaches to define glioma cellular states at high granularity and uncover their organization. We find three prominent modes of cellular organization. First, cells in any given state tend to be spatially clustered, such that tumors are composed of small local environments that are each typically enriched with one major cellular state. Second, specific pairs of states preferentially reside in proximity across multiple scales. Despite the unique composition of each tumor, this pairing of states remains largely consistent across tumors. Third, the pairwise interactions that we detect collectively define a global architecture composed of five layers. Hypoxia appears to drive this 5-layered organization, as it is both associated with unique states of surrounding cells and with a long-range organization that extends from the hypoxic core to the infiltrative edge of the tumor. Accordingly, tumor regions distant from any hypoxic foci and tumors that lack hypoxia such as IDH-mutant glioma are less organized. In summary, we provide a conceptual framework for the organization of gliomas at the resolution of cellular states and highlight the role of hypoxia as a long-range tissue organizer.

## Introduction

In his landmark 1932 paper, Percival Bailey elegantly characterized the spatial architecture of gliomas observing that "the microscopic structure of tumors of the brain is infinitely varied, yet among their kaleidoscopic appearances certain family resemblances may be traced…".^1^ Gliomas are typically highly heterogeneous and infiltrative, yet specific spatial patterns are consistently noted. In 1938, Hans Joachim Scherer detailed the recurring spatial structures of gliomas and, in particular, glioblastomas: organized secondary structures characteristic of invasion in which glioma cells rely on or mimic the pre-existing normal structures of the brain; amorphous arrangements of cancer cells; proper structures in which the cancer cells form patterns that do not depend on pre-existing normal structures; and a mesenchymal tissue response surrounding areas of necrosis.^2^ Accordingly, foci of necrosis (and hypoxia) as well as neuronal structures of the normal brain are associated with local organization, while glioma tissues are dominated by lack of histological organization.^3^

More recently, single-cell RNA sequencing (scRNA-seq) has reshaped our understanding of gliomas by capturing the full diversity of malignant cellular states and nonmalignant cell types from which they are composed. We previously defined four major cellular states for glioblastoma cancer cells: neural progenitor-like (NPC), oligodendrocyte progenitor-like (OPC), astrocyte-like (AC), and mesenchymal-like (MES).^4^ All states but the MES state mimic the gene expression programs of normal neurodevelopmental cell types; in contrast, the MES state is most influenced by the tumor microenvironment.^5^ IDH mutant gliomas (astrocytomas and oligodendrogliomas) contain a cycling NPC-like population as well as astrocyte-like (AC) and oligodendrocyte-like (OC) cancer cell states.^6, 7^

Additional studies identified specific spatial interactions among glioma cells. Examples include macrophages that induce the GBM MES state by secreting OSM, which binds to OSMR on cancer cells^5^, and synaptic crosstalk between OPC-like GBM cancer cells and neurons via NLGN3.^8^ However, many open questions remain concerning how the cell states and cell types relate to each other comprehensively and globally: Given their remarkable spatial diversity, are there consistent patterns or rules of organization across gliomas? To what degree does spatial location dictate the diversity of cellular states we and others described previously?

Spatial transcriptomics and proteomics methods allow us to bridge the gap between the features and patterns described by classical histology and the granular view of cell state and cell type granted by scRNA-seq. To this end, we profiled 19 glioma samples by 10X Visium and 12 by CODEX and integrated them with published GBM Visium data.^9^ We further developed a novel framework to describe the spatial organization of GBM quantitatively and systematically across platforms and multiple levels of resolution.

## Results

### Spatial transcriptomics of glioma

We used the 10X Visium spatial transcriptomics platform to spatially profile gliomas by RNA sequencing (**Fig. 1A**).^10, 11^ The diameter of a capture spot is 55µM, such that each spot contains a mixture of 1-35 cells, with a median of 8 cells in GBM based on image analysis (**Fig. S1A**). In four cases, we analyzed multiple frozen tissue blocks from the same tumor, each isolated from a different region and annotated as *necrotic*, *infiltrating*, or *T1 contrast-enhancing* by navigation-guided tumor resection. We profiled 13 IDH-WT glioblastoma sections and 6 IDH-mutant glioma sections (3 oligodendroglioma, 3 astrocytoma) (**Table S1**). These samples were integrated with 13 GBM sections from an external Visium dataset.^9^ We retained 70,618 spots for analysis after quality control (**Methods**).

**Figure 1.**
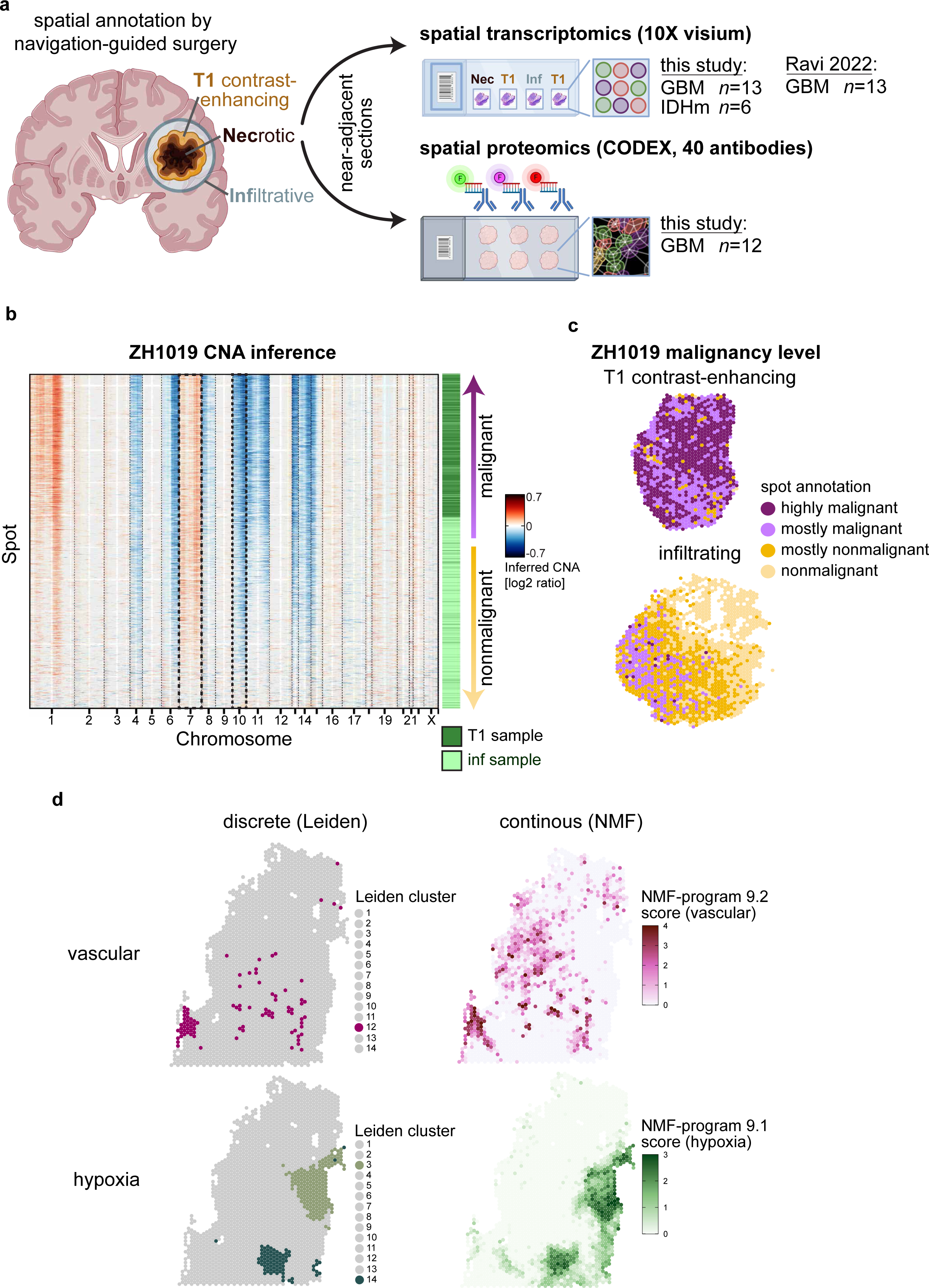
Experimental design and spot classification per sample. **(a)** Experimental design and patient cohort. Fresh frozen tissue sections from GBM (*n=*13) and IDH mutant gliomas (*n=*6) were profiled by 10X Visium spatial transcriptomics. CODEX spatial proteomics was performed on 12 near-adjacent GBM tissue sections with a panel of 40 antibodies. Four tumors were spatially annotated by the neurosurgeon during navigation-guided surgery. This GBM Visium cohort was combined with an external GBM Visium cohort (Ravi et al. 2022) for joint analysis. Created with BioRender.com. **(b)** Copy number aberrations (CNAs) were inferred by average relative expression in windows of 150 analyzed genes. ZH1019, from which we profiled both an infiltrating sample and T1-constrast enhancing sample, is shown here as an example. Rows correspond to spots arranged by malignancy level as inferred from CNA (Methods), columns correspond to genes arranged by chromosomal position. Annotation bars correspond to region from which the spot was derived (T1 or infiltrating) and malignancy level. **(c)** Spatial maps of ZH1019 T1 and ZH1019 infiltrating samples with spots annotated by malignancy level as described in (b). **(d)** Per sample Leiden (left) and NMF clustering (right) of ZH916bulk for vascular and hypoxia clusters (all other clusters are shown in gray).

We used two initial approaches to classify and annotate spots. First, we performed CNA inference by averaging the expression of genes in each chromosomal region to classify spots as mostly malignant, mostly nonmalignant, or mixed (**Methods**).^6, 12^ As expected, inferred CNAs included the hallmarks of GBM (chromosome 7 amplification and chromosome 10 deletion) and oligodendroglioma (chromosome 1p/19q co-deletion) and were significantly associated with cancer-rich regions (e.g., tumor core vs. infiltrating regions) (**Figs. 1B-C, Figs. S1B-C**). Second, we annotated spots by Leiden clustering both per sample and jointly per tumor in cases with multiple sections (**Fig. 1D, Fig. S1D**).

While the per-sample Leiden clustering discretely groups together spots of similar composition within samples, we next aimed to better capture the continuous nature of cellular states and the mixing of cells within spots. To this end, we performed non-negative matrix factorization (NMF) per sample and derived robust expression programs that were consistently detected across multiple parameters (**Fig. 1D, Methods**).^13, 14^

### Recurrent patterns of expression heterogeneity across gliomas

In order to define expression programs that reflect core patterns of expression heterogeneity, we compared the 492 gene expression programs identified in individual GBM samples either by Leiden clustering or by NMF. We found considerable similarities that allowed us to define 14 clusters of programs based on their gene overlap, each covering programs derived from multiple GBM samples (**Figs. 2A-B**, **Methods**). For each group of programs, we defined a consensus program of 50 genes, termed a metaprogram (MP), that reflects a recurrent pattern of heterogeneity in GBM (**Fig. 2C, Fig. S2A, malignant MPs in bold**). Similarly, we defined six MPs in IDH-mutant glioma (**Figs. S2B-C**, **Methods**).

**Figure 2.**
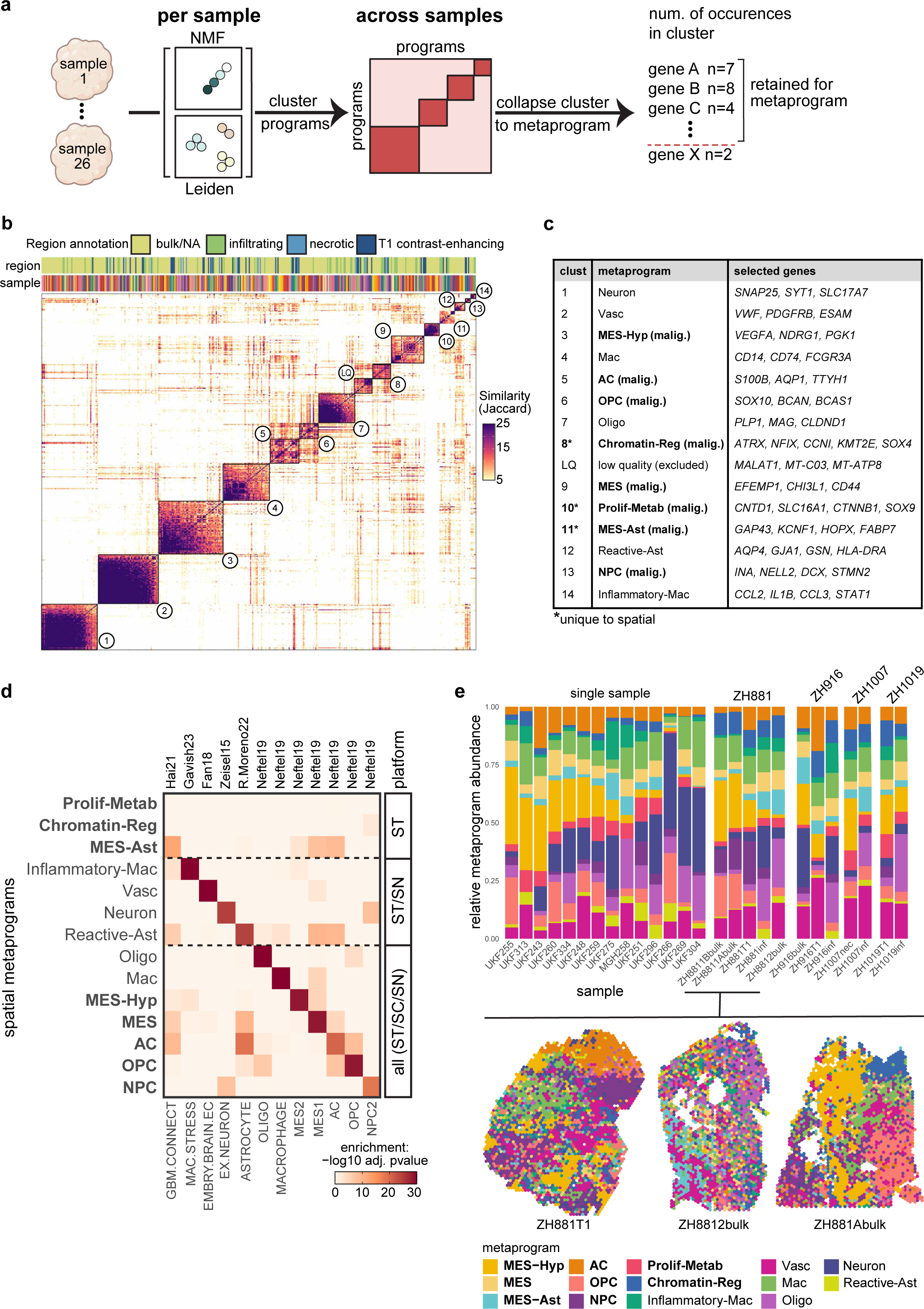
Deriving and annotating spatial metaprograms (MPs). **(a)** Scheme of metaprogram generation approach. Each sample is clustered individually by Leiden and NMF. All Leiden and all NMF programs across all samples are clustered by their gene overlap. Each cluster is collapsed to a consensus MP by selecting for the most recurrent genes across programs within the cluster. **(b)** Similarity matrix based on gene overlaps (quantified by Jaccard index), for all programs derived from NMF and Leiden clusters. Programs are annotated by sample and region identity. Cluster numbers correspond to the table in (c). **(c)** Table of metaprogram names and selected genes corresponding to clusters numbered in (b). Malignant metaprograms are depicted in bold. **(d)** Enrichments of spatial metaprograms (rows) with gene-sets (columns) previously defined from studies indicated at the top.^4, 9, 14, 33–35^ Enrichment calculated by hypergeometric test. **(e)** For each sample (represented by one bar), shown are fractions of spots assigned to each MP. Four tumors with multiple sections are at the left and are grouped by tumor. Below are spatial MP maps of several ZH881 tissue sections, demonstrating differences in composition and spatial organization between sections isolated from the same tumor.

While most Visium spots contain a mixture of multiple cell states and cell types, we speculated that MPs reflect the most robust signal that recurs across many spots, thereby highlighting the dominant cell state/type of the associated spots. CODEX analysis (described below) of Visium-sized spots, denoted as pseudospots, supported this speculation by revealing that 64% of pseudospots are dominated by one MP that accounts for most cells in that spot, and 99% of pseudospots are dominated by one or two MPs (**Fig. 3G**, **Fig. S2D**). Therefore, cells within a given spot tend to share the same state, such that spatial MPs may capture the signature of that cell state/type, allowing us to derive single-cell analogous programs directly from the spatial data (**Fig. 2D**). Indeed, our MPs are highly similar to those derived from single cell RNA-seq data and generally do not map to combinations of single-cell states, except for the combination of endothelial cells and pericytes that are tightly coupled (**Figs. S2E-F**). Our approach obviates the need for single-cell or single-nuclei profiling of adjacent samples or for external single-cell references, as done by other spatial transcriptomics studies with multiple limitations.^15, 16^ Our MP approach also differs from earlier work that focused on spatially-defined programs containing a mixture of cell types (**Fig S2H,** Ravi et al. 2022).

**Figure 3.**
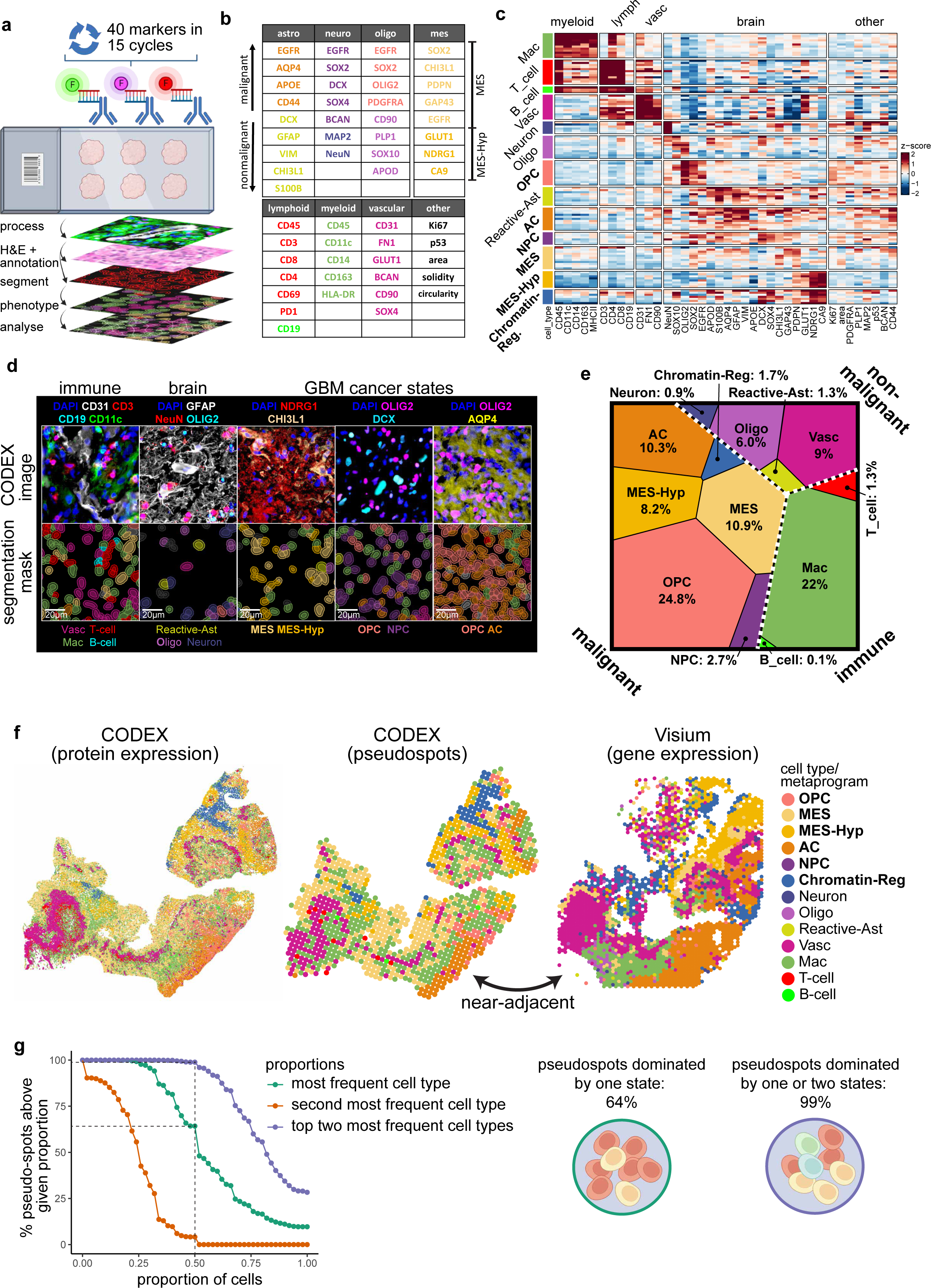
Spatial profiling of gliomas by CODEX. **(a)** Schematic workflow of CODEX experiment, image processing and computational analysis. Created with BioRender.com. **(b)** Protein markers profiled by CODEX. **(c)** Relative protein expression per cell type (or state). Columns represent proteins. Each row represents the indicated cell type/state in one sample. Number of rows differ between cell types/states because some samples lack specific cell types/states. **(d)** CODEX staining representative examples: Top row shows images with the indicated markers. Bottom row shows the corresponding nuclear segmentation masks with cytoplasmic expansion by 3 µm. Masks are color-coded by cell phenotype. **(e)** Voronoi diagram showing cell type/state abundances across all samples. Dashed lines separate malignant from non-malignant and immune cells. **(f)** Exemplary spatial maps of cell types/states annotations for (i) CODEX single cell data, (ii) CODEX pseudo-spots (annotated by most frequent cell type/state) and (iii) Near-adjacent section: Visium spots annotated by the highest expressed MP. **(g)** Left: Cumulative distribution plot showing the frequency of pseudospots in which, the highest cell type/state, the second highest, or both of them, cover a higher proportion of the cells than the value specified on the x-axis. Right: Schematic reflecting proportion of pseudospots dominated by one state and by one or two states.

We identified 14 GBM spatial MPs, including eight malignant and six nonmalignant programs, each reflecting a cancer cell state or non-malignant cell type (**Figs. 2C-D, Fig. S2G, Table S2**). Nonmalignant MPs included Mac (macrophage) and Inflammatory-Mac, Oligo (oligodendrocyte), Vasc (vascular -endothelial cells and pericytes), Neuron, and Reactive-Ast (reactive astrocyte). The latter included classical astrocytic markers (e.g., *AGT, GJA1*) and additional markers suggesting a reactive astrocytic state (e.g., MHC-II and metallothioneins). Of the eight malignant MPs, five directly map to the single-cell GBM states: MES-Hypoxia (MES2), MES (MES1), NPC, OPC, and AC (**Fig. S2I**). The three additional malignant spatial MPs include: (i) an astrocytic-like mesenchymal MP (MES-Ast) with enrichment of genes associated with glioma tumor microtubes^17, 18^ (e.g., *GAP43, KCNF1, PTN*) (**Figs. S2J-L)**, (ii) Proliferation and metabolism (Prolif-Metab), enriched with proliferation-related (e.g., *TP53, CTNNB1, CNTD1*) and metabolism (e.g., *SLC16A1* (MCT1)*, GGCX, PHGK1*) genes, and (iii) Chromatin regulation (Chromatin-Reg), enriched with chromatin and transcriptional regulators (e.g., *ATRX, KMT2E, BRD4, SOX4*). MPs identified in IDH mutant glioma include AC, an OC/NPC1-like program unique to IDH mutants, Mac/Microglia, Oligo, Neuron, and Vasc (**Figs. S2B-C, Table S2**).

Next, we annotated all spots by their highest-scoring spatial MP (**Fig. 2E, Fig. S2M**). Due to the relatively small number of IDH mutants (*n*=6), we annotated the spots of IDH mutant tumors with MPs defined from GBM samples that were highly represented in the IDH mutant spots in addition to the IDH mutant MPs (**Methods**). The overall composition of each GBM sample was highly variable, with MES-Hyp and Neuron frequencies varying the most between samples and MES frequency varying the least. Samples isolated from different regions of the same tumor were highly variable in their cell type composition (**Fig. 2E, Fig. S1D**), highlighting the degree of sampling bias when a single tissue section is considered representative of the composition of an entire tumor.

### Spatial proteomics as a complementary approach to map gliomas

While Visium provides comprehensive data by covering most genes, it suffers from low spatial resolution. Therefore, we used CODEX as a complementary approach to validate our findings on a true single-cell protein level in large tissue sections (6.5x6.5mm). Our 40-marker antibody panel was designed based on single-cell GBM MPs along with canonical markers to cover almost all relevant cell types and cell states (**Figs. 3A**-**B, Table S3**). After quality control, we retained 428,395 cells from 12 samples (**Fig. 3E**, **Methods**). All major differentiated nonmalignant cell types were identified, including astrocytes, oligodendrocytes, neurons, vascular cells (endothelial cells and pericytes), T cells, B cells, and macrophages/microglia (**Figs. 3C-D, Fig. S3B**). We also identified the major malignant cell states including MES, MES-Hyp, Chromatin-Reg, OPC, NPC, and AC. Assignment of MPs to CODEX data was highly consistent with the assignments of Visium data from near-adjacent sections, supporting the robustness of both platforms (**Fig. 3F, Fig. S3A**).

In Visium data, several cell types and states were not detected by unsupervised analysis since they tend to represent a minor component of the spots in which they reside. For example, we did not identify a cell cycle MP, raising the possibility that, in most cases, a cycling cell is surrounded by many non-cycling cells that dilute the cell cycle signal (**Fig. S3C**). Similarly, T cells and B cells rarely dominate a spot, and their low mRNA content further limits their signal such that we could not identify T cell or B cell MPs by unsupervised analysis. Accordingly, CODEX was better suited for identifying these cell types at the single-cell level (**Fig. 3D**). T cells were detected in all samples but were lowly abundant in most tissues (0.4% median per sample) whereas B cells were absent in all but three samples (**Fig. S3B**). In line with our previous findings^4^, OPC was the most cycling malignant state (8.5% cycling), while Vasc was the most cycling nonmalignant state (5.1% cycling), highlighting abundant microvascular proliferation, a hallmark of GBM (**Fig. S3D-E**). Supervised cell cycle analysis in the Visium data further supported these results and additionally showed that among the malignant states, Prolif-Metab contained the largest percentage of cycling spots (median 10.6%) followed by OPC (median 9.4%) (**Fig. S4A**, **Methods**).

The single-cell resolution of CODEX data enabled us to assess the cell density of each state. Nonmalignant brain cell types had lower density (median 5 cells/spot) than malignant cells, immune cells and endothelial cells (median of 9 cells/spot). Of the malignant states, Chromatin-Reg and MES were the densest, while NPC, OPC, and AC were the least dense (**Figs. S3E-F**). Moreover, we calculated average cell densities for classical GBM histology features. As expected, *infiltrating* regions had the lowest density (6 cells/spot), while *microvascular proliferation (MVP)* (16 cells/spot) and *pseudo-palisading* (18 cells/spot) had the highest cell density, though in the case of *pseudo-palisading* this increased density was not associated with increased proliferation (**Fig. S3G**).

### Structured vs. disorganized regions

After annotating each spot by its highest scoring spatial MP (**Fig. S2M, Fig. S3A**), we wondered to what degree each state is enriched in neighboring vs. in scattered spots in each sample, which we term *spatial coherence* (**Fig. 4A, Methods**). Spatial coherence varies between samples, such that in some samples (termed *structured*), most states tend to have high coherence, while in other samples (termed *disorganized*), most states tend to have low coherence (**Fig. 4B**). While most states varied together in spatial coherence based on sample and location, a few states were either consistently grouped (high spatial coherence) or scattered (low spatial coherence) (**Fig. 4C**). These exceptions included MES-Hyp and Neuron which had consistently high spatial coherence across samples and Prolif-Metab, which had consistently low spatial coherence. Apart from these exceptions, spatial coherence varied more across samples than across states, indicating that it is more region-specific than state-specific (two-way ANOVA: sample effect *p*=0.000003; state effect *p*=0.23).

**Figure 4.**
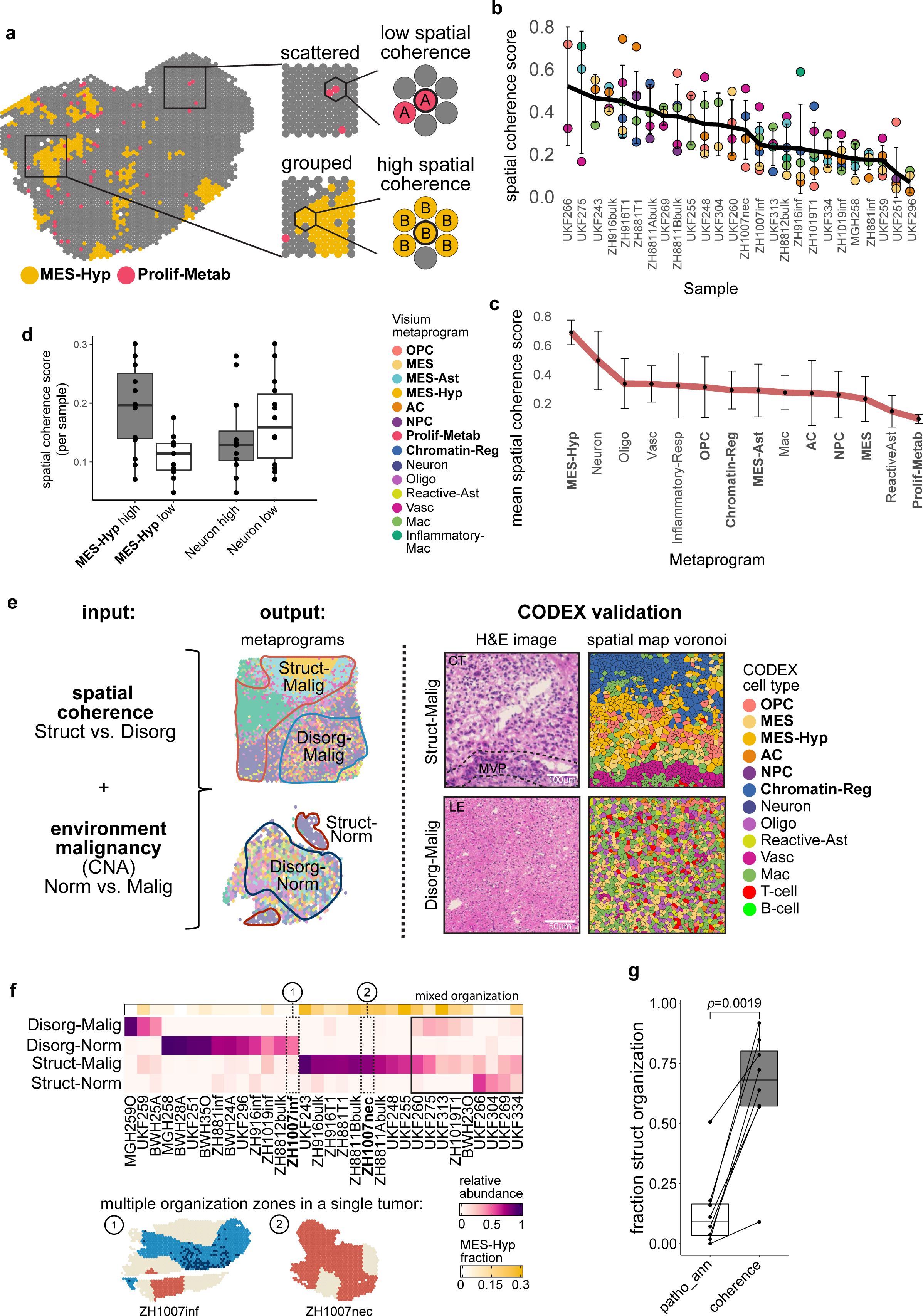
Spatial distribution of cell states and organization zones. **(a)** Scheme illustrating spatial coherence score calculation depicting high spatial coherence score for a grouped cell state and a low spatial coherence score for a scattered cell state. **(b)** Cell type/cell state spatial coherence score by sample. Standard deviation is shown in error bars. **(c)** Cell state/type mean spatial coherence score across all samples, standard deviation is shown in error bars. (**d)** Mean spatial coherence score of samples with high abundance (>10%) of MES-Hyp or Neuron vs. samples with low abundance (<10%) of MES-Hyp or Neuron. MES-Hyp or Neuron spots were removed from the calculation of mean spatial coherence scores. **(e)** Left: Scheme describing the generation of spatial organizational zones. Spatial coherence and spot malignancy level (inferred from CNA as described in Fig. 1B) were used to assign spots to Structured-Malignant (Struct-Malig), Structured-Normal (Struct-Norm), Disorganized-Malignant (Disorg-Malig) and Disorganized-Normal (Disorg-Norm) in Visium data. Right: Structured and disorganized regions are also found in CODEX data. H&E of Struct-Malig zone shows microvascular proliferation (MVP) but no pseudopalisades or necrosis in the region that is structured at the cell state level. **(f)** Top: Heatmap of the relative abundance of the different organizational patterns (rows) in each sample (columns). Annotation bar shows per sample MES-Hyp abundance. Bottom: Example of samples coming from the same tumor showing multiple organization zones. **(g)** Fraction of structured regions based on histology (MVP and PAN) vs. fraction of structured regions based on cell state spatial coherence in the same samples (CODEX) (*p*=0.0019).

To further explore this distinction, we devised a computational approach to classify each spot as belonging to a *structured*, *disorganized,* or *intermediate* region based on the coherence of its local environment (**Methods**). We also identified structured and disorganized regions in the CODEX data, with high consistency between near-adjacent Visium and CODEX samples (*R*=0.84, *p*=0.0013; **Fig. 4E, Figs. S4B-C, Fig. S3Av**). Structured regions (48% of total spots) were found both at the core of tumors (enriched with MES-Hyp) and in infiltrated areas of the normal brain (enriched with Neuron) (**Fig. 4F, Fig. S4D-E**). Therefore, we further subdivided structured and disorganized regions into malignant environments based on CNA signal: Struct-Malig (*n=*20 regions, 39% of spots) and Disorg-Malig (*n*=10 regions, 22% spots) and nonmalignant environments: Struct-Norm (*n=*6 regions, 8% spots) and Disorg-Norm (*n*=4 regions, 6% spots) (**Fig. 4E, Fig. S4F**). All GBM tumors with multiple sections were comprised of both structured and disorganized compartments, suggesting that the co-occurrence of these patterns is a recurring spatial feature of GBM (**Fig. 4F, Figs. S4F-G**). Struct-Norm regions reflected the organization of the nonmalignant brain parenchyma and had the highest frequency of Neuron, while Disorg-Norm was composed of normal brain regions with a high degree of cancer infiltration and had the highest frequency of Oligo (**Fig. S4E)**.

As noted above, MES-Hyp was enriched in Struct-Malig regions. Samples enriched with MES-Hyp had high mean spatial coherence even after removing the MES-Hyp spots from the calculation of spatial coherence (*p*=0.0001, **Fig. 4D**). Therefore, the increased organization associated with hypoxia extends beyond hypoxia itself, and the malignant cell states in proximity to MES-Hyp spots were more organized than those same states in regions lacking MES-Hyp spots. Accordingly, structured regions around hypoxia were significantly more abundant (*p*=0.0019) than the two histological annotations associated with hypoxia-related structure – *pseudopalisading and necrosis* (PAN) and *microvascular proliferation* (MVP) **Fig 4G**). While Neuron spots were also highly spatially coherent, this effect of extended structure on adjacent cell states was not observed for Neuron-containing regions (*p*=0.35, **Fig. 4D**). Thus, hypoxia may provide an extrinsic force that confers a structured continuum of expression states over large tissue regions^30^ resulting in a known local structure that is visible by standard histology and a previously unappreciated long-range structure that can only be detected by molecular profiling of cellular states. Hypoxia-poor regions in both GBM and IDH mutant gliomas were largely disorganized (Disorg-Malig) (**Fig. 4F, Fig. S4E, Fig. S4H**), consistent with the overall disorganization of gliomas seen on histology.

While MES-Hyp and Neuron reflected the greatest difference between structured and disorganized regions, we observed additional differences in composition. Vasc spots were more abundant in Struct-Malig regions (*p*=0.03), while Prolif-Metab was more abundant in disorganized regions (*p*=0.003) (**Fig. S4I**). Vasc could be further subclustered into two populations with differing spatial coherence – an angiogenic (Vasc-Ang) program with high coherence and enrichment in structured regions, and an immunomodulatory (Vasc-IMEC) program with lower coherence and enrichment in disorganized and infiltrative regions (**Table S2**, **Figs. S4J-M**).

### State-State spatial associations

To quantify spatial relationships between states, we devised three complementary measures highlighting state coupling at varying levels of resolution (**Methods**). First, *regional composition* of two states, defined as the correlation between their abundance across hexagonal windows of a predefined radius (denoted by *r*) (**Fig. 5A**). Second, *adjacency* between two states, defined as the enrichment of one state in the immediate neighboring spots of the other state. Third, the *colocalization* of two states within the same spot (or pseudo-spot in CODEX). We observed overall consistency of state relationships over increasingly sized areas, while also identifying specific scale-dependent shifts (*distance-dependent coupling*) when considering all measures together (67% consistency across measures in structured regions, 57% consistency across measures in disorganized regions) (**Fig. 5F**, **Supp. Note 1**).

**Figure 5.**
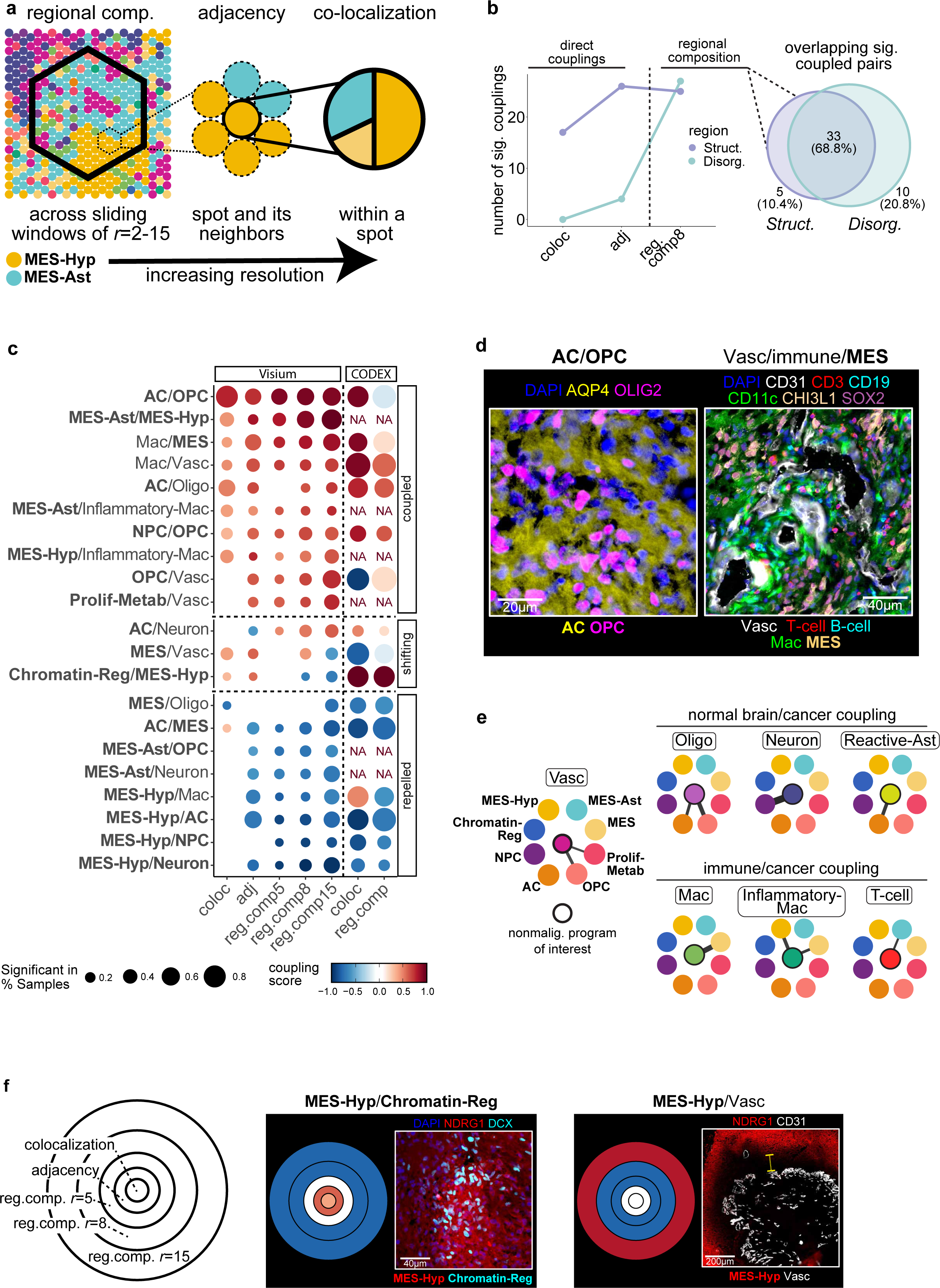
Spatial associations between states across scale. **(a)** Scheme depicting the three measures of spatial relationships between MPs across scales (from low to high resolution): i) Regional composition (across sliding windows of *r*=2-15), ii) Adjacency, and iii) Colocalization (Methods). The example shown highlights coupling between MES-Ast and MES-Hyp across scales of resolution. **(b)** Left: Line plot showing the number of significant interactions across different analyses in structured vs. disorganized regions. Right: Venn diagram of the number of significant interactions across the regional composition analysis. **(c)** Summary heatmap of state-state associations across scales in structured samples covering most consensus interactions and other pairs of interest (for full table see Supp. Note 1). Each column represents the summary of a different spatial relationship measure (Visium data) across all GBM samples, with the final two columns corresponding to CODEX colocalization and regional composition, respectively. Dots are colored by mean scaled relationship strength (Methods) and dot size corresponds to the fraction of samples in which the relationship is significant. **(d)** Images highlighting selected pairs of coupled cell types from (b). Left (CODEX): OPC/AC coupling. Right (CODEX): Spatial relationships in the vascular niche. While Vasc and immune cells are immediately adjacent, Vasc/MES coupling occurs over larger areas. **(e)** Top: Network graphs representing couplings between normal brain cell types or Vasc with the malignant states, Bottom: Couplings of immune cells with malignant states. In each graph, the central node is a nonmalignant cell type and the surrounding nodes represent cancer states. Edge strength represents the mean scaled relationship strength across all measures (Visium). Only consensus interactions are shown. **(f)** Nested circle plots depicting changes in spatial relationship strength across scales of resolution (distance-dependent associations). Left: Schematic of nested circle plot. Right: Examples of distance-dependent associations with nested circle plots (Visium) adjacent to immunonstaining.

Structured and disorganized regions had a similar number of significant spatial associations by regional composition, and these associations were largely consistent, while the colocalization and adjacency measures mostly identified interactions that were specific to structured regions (**Fig. 5B**). Therefore, regional composition tends to be maintained even in the absence of an overall structure, indicating that specific pairs of states are spatially correlated within glioma regions; yet, spatial patterning involving direct recurring spatial relationships were mostly lost in disorganized regions.

To identify the most robust interactions that drive the organization of structured GBM regions, we defined a consensus set of state-state interactions that are supported by multiple measures and across multiple samples **(Methods)**. We found ∼10-fold more consensus interactions in structured GBM regions than in disorganized GBM regions (21/91 vs. 2/91 state-pairs) (**Fig. 5B**). The only consensus interactions in disorganized GBM regions were MES/Mac and NPC/Neuron. Likewise, IDH mutants, which are mostly disorganized, had fewer consensus interactions (9/55 pairs). The consensus interactions were validated by CODEX (**Figs. 5C, Fig. S5A, Supp. Note 1**). Most consensus interactions are briefly summarized below **(Fig. 5C)** and all interactions are described in more detail in **Supp. Note 1**.

Malignant states were divided into two groups, each with many within-group interactions: (i) neurodevelopmental states (OPC, NPC and AC) and (ii) a set of mesenchymal and hypoxia-associated states (MES-Hyp, MES-Ast, MES and Chromatin-Reg) (**Figs. 5C-D, Fig. S5A**). The neurodevelopmental malignant states not only interacted among themselves but also with the non-malignant states corresponding to the same lineage – AC (malignant) interacted with Reactive-Ast, NPC (malignant) interacted with Neuron, and OPC (malignant) interacted with Oligo (**Fig. 5E**). OPC also interacted strongly with Vasc, reminiscent of OPC-endothelial migration^19^ and cross-talk^20^ in development. Likewise, Prolif-Metab was strongly associated with Vasc, in line with a previously proposed metabolic vessel-associated GBM state.^21^

The three mesenchymal states all had strong interactions with immune cells, consistent with earlier studies.^5^ However, the mesenchymal states differed in several immune-related ways. First, while MES and MES-Ast interacted strongly with Mac, MES-Hyp instead interacted more strongly with Inflammatory-Mac, which are thought to be immune-suppressive **(Fig. 5E)**. Second, MES-Ast was the malignant state most enriched with T cells, while MES-Hyp was the malignant state most depleted of T cells (**Fig. S5B**). Third, MHC-II expression varied across mesenchymal states (MES-Hyp: MHC-II^low^; MES-Ast: MHC-II^med^; MES: MHC-II^high^), consistent with hypoxia-mediated downregulation of MHC class II (**Figs. S5C-D**).^22, 23^ Additionally, Inflammatory-Mac were associated with reduced expression of MHC class II and a bias of T cells towards CD8+ (**Figs. S5B and E-F**). Collectively, these results support the possibility that distinct MES states may have opposing effects on immune activity.^24, 25^

Immune cells (Mac, T cells, and B cells) also strongly interacted with Vasc, highlighting their dependence on trafficking (especially for bone marrow-derived macrophages) **(Fig. 5C, Fig. S5A)**. While Vasc was coupled to immune cells, it was decoupled from MES-Hyp, as expected **(Fig. 5F)**. This decoupling was observed up to a distance of ∼160 μM, suggesting that this is the effective length scale of vascular perfusion and that beyond this distance from blood vessels, there is a hypoxia response (**Fig. S5G**).

### A layered model of GBM spatial organization for structured regions

To examine whether the individual interactions described above may combine to form a higher-order organization, we generated a network graph in which nodes represent states and edges represent consensus interactions (**Figs. 6A-B**). This graph reveals a five-layered organization in which edges exist only within the same layer or between adjacent layers. This organization appears to be dominated by a hypoxia gradient, with a first layer consisting of MES-Hyp (***L1****: **core hypoxia***), followed by a second layer consisting of MES-Ast, MES, and Inflammatory-Mac (***L2: hypoxia-associated***). The third layer includes immune and angiogenesis-related cell types and states that may help to resolve the hypoxia, including Vasc, Mac, and Prolif-Metab (***L3: angiogenic response***). Following *L3*, hypoxia is presumably resolved, enabling the presence of neurodevelopmental malignant states that may be more oxygen-dependent – AC, OPC, and NPC (***L4: malignant oxygen-dep.***). Finally, nonmalignant brain cell types (Reactive-Ast, Oligo, and Neuron) reflect the transition to the infiltrated brain parenchyma (***L5: brain parenchyma***).

**Figure 6.**
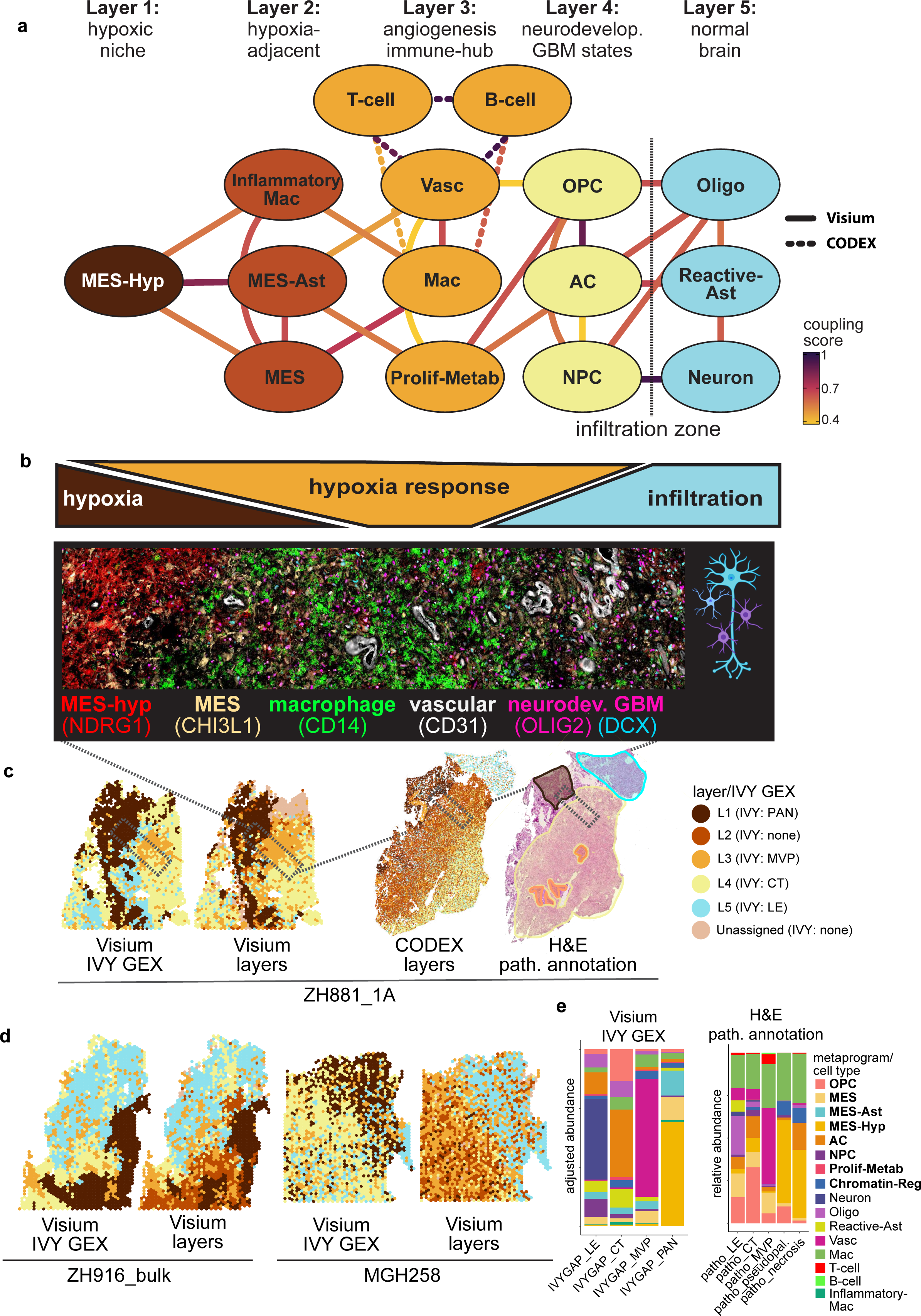
A layered model of GBM spatial organization. **(a)** Top: Network graph with nodes representing cell types/states and edges representing recurrent interactions (found in > 20% of samples). Edge color represents mean coupling score across levels of resolution, for structured regions of Visium data. Edges with dashed lines represent connections with cell types coming from CODEX (T cells, B cells). Bottom: Scheme showing gradients of hypoxia, hypoxia response and infiltration in alignment with the layers. **(b)** CODEX image showing the indicated cell types and markers, representing layers 1 – 4. Layer 5 was added as scheme depicting normal brain. Created with BioRender.com. **(c)** Spatial maps for sample ZH8811A: i) Visium sample annotated by IVY Gap transcriptional programs ii) Visium and iii) CODEX sample annotated by state layers iv) Neuropathologist annotations of H&E staining from the CODEX sample. **(d)** Spatial maps comparison for Visium samples annotated by IVY Gap transcriptional programs (left) vs. annotated by our layers (right) **(e)** Stacked bar plot showing metaprogram composition per Ivy GAP transcriptional program annotation (left) and metaprogram composition per histological annotations (right).

Spatially mapping the layers revealed that they preserve the structured and disorganized nature of sample regions (**Fig. 6C, Figs. S6A-B**). CODEX data generated a similar graph with the addition of B cells and T cells to *L3* (*angiogenic response*) (**Figs. S6B-C**).

Finally, we asked how our 5-layer model relates to classical histological annotations of glioma, including *pseudo-palisading and necrosis* (PAN), *microvascular proliferation* (MVP), *cellular tumor* (CT), and *leading edge* (LE). We used two approaches to histologically annotate our samples - H&E annotations by a neuro-pathologist and transcriptional signatures of histological features as defined by Ivy GAP (**Fig. S6B**).^26^ The histological annotations were overall in agreement with the layers with a few inconsistencies. For example, the Ivy GAP annotations fail to capture the hypoxia adjacent layer (*L2*) (**Fig. 6D left, Fig. S6A**). Therefore, by the Ivy GAP programs, *L2* spots (MES-Ast, MES, Inflammatory-Mac) would be classified as either PAN or MVP (**Fig. 6E, Fig. S6D**). Additionally, spots belonging to *L4* (OPC, AC, NPC) are often classified as LE by the Ivy GAP gene expression programs (**Fig. 6D right, Fig. S6D**) or histological annotation (Fig. S6F). Lastly, small areas of local hypoxia that are classified by Visium as *L1* and *L2* are found in regions outside of visible PAN and therefore histologically defined as CT and infiltrating/LE zones (**Fig. S6E**). Therefore, by unsupervised analysis of gene expression we largely converged to the major histological features of glioma, yet with increased accuracy and with the addition of a new layer (*L2*) of hypoxia-adjacent states. Moreover, we can now redefine the classical histological features at a detailed resolution of cellular states (**Fig. 6E, Fig. S6E**).

## Discussion

We combined spatial transcriptomics with spatial proteomics and novel computational approaches to define the organization of gliomas. Previous analysis of spot-based spatial transcriptomics has relied on deconvolution with matched single-cell or single-nuclei RNA-Seq.^15, 16, 27^ In addition to being costly and labor-intensive, this approach suffers from under-represented or missing cell types in dissociated tumors. Instead, we demonstrated that it is possible to derive single-cell analogous MPs directly from spatial transcriptomics data by unsupervised analysis, thereby avoiding the technical issues associated with integration of disparate data types. Moreover, we identified several MPs in the spatial transcriptomics data that were not previously defined from single-cell or single-nuclei data. Some of these programs could be retrospectively identified in single-cell data by supervised analysis (**Figs. S2E-F and I-L**), suggesting that spatial clustering of rare or subtle cellular states may have amplified their signals and facilitated their identification by Visium. Conversely, the signals for cellular states with limited spatial clustering are diluted by Visium, as is the case for cell cycle (**Fig. S3C**)

Overall, we found three modes of spatial organization. First, cells tend to be surrounded by other cells in the same state (*state-specific clustering*), forming local environments that are highly enriched with an individual state. This mode is evident by the clustering of similar cells within each spot, as shown directly by CODEX, as well as by the clustering of spots assigned to the same MP. This clustering suggests that spatial location plays a central role in regulating cell state.

Second, many pairs of states are consistently associated across multiple scales in structured regions – within spots, in adjacent spots and within small tissue regions (*state-state association*). Some of these interactions may mimic those of normal brain cell types in development. For example, malignant OPC are coupled to Vasc, consistent with migration of normal OPCs along the brain endothelium^19^ and NPC is coupled with neurons in infiltrative areas, mimicking the migration of normal NPC towards developing neurons.^28, 29^ The exact mechanisms driving state-state associations require further studies and may reflect not only physical connections, but also recruitment through secreted factors, dynamic transitions from one state to the other, synergistic growth or survival of the interacting cells, or a common dependence on microenvironmental components.

Third, the associations between states in structured regions aggregate to form a higher-order organization with five layers, such that states in each layer are associated only with the same layer or adjacent layers (*state layers*). The presence of higher-order organization is reminiscent of normal tissue structure and may facilitate large-scale coordination between cancer cell states that could present a therapeutic opportunity. Four layers are largely consistent with histological features, allowing us to redefine the classical GBM histology by specific cell states and to add the novel layer of hypoxia-associated states (*L2*).

Hypoxia appears to be a central driver of this organization, and each layer can be interpreted by its relation to hypoxia. Importantly, the effect of hypoxia on spatial organization extends beyond the features visible by histology. Therefore, while the known structure of gliomas is typically restricted to small areas of hypoxia or brain parenchyma, structure at the level of cell states extends further. Additionally, some of the hypoxic regions defined by spatial-omics were not visible by histology yet still capable of generating structured organization of cell states. This hypoxia-centric model is consistent with recent modeling studies, which suggested that an external force such as hypoxia acting over a large area can result in a spatial gradient that generates a continuum of expression states.^30, 31^ Likewise, hypoxia gradients were shown to generate a continuum of macrophage states generating predictable spatial patterns in mouse breast cancer models.^32^

Although tissue organization may be intuitively linked to normal physiology while chaos is reminiscent of aggressive tumor phenotypes, we note that hypoxia, and hence cancer cell state organization, are hallmarks of high-grade glioma. Such organization may imply that certain layers are less accessible to drugs or to immune cells and thereby more resistant to particular therapies. In contrast, lower grade IDH-mutant gliomas tend to lack hypoxia and hence are typically disorganized. In the atypical case of a lower-grade glioma that does have a high degree of hypoxia we also observe a high degree of organization, consistent with the model of hypoxia as driving spatial organization (**Fig. S2M, Fig. S4H**). It is tempting to speculate that vascular normalization may decrease hypoxia and aid in reverting a structured organization into that which is typical of low-grade glioma.

In addition to its association with global organization, hypoxia appears to induce specific states in its vicinity, including the core hypoxic state of MES-Hyp, but also Chromatin-Reg and three additional states that constitute the hypoxia-associated layer. Two of these reflect malignant states not identified in our previous scRNA-seq analysis (Chromatin-Reg and MES-Ast). We further speculate that most (or all) of these MPs represent hypoxia-associated versions of other MPs observed further away from hypoxia. Pairs of related malignant states that may differ due to hypoxia proximity include MES-Hyp vs. MES, MES-Ast vs. AC, and Chromatin-Reg vs. NPC. For example, Chromatin-Reg includes upregulation of chromatin regulators but also of NPC-related genes (i.e., *SOX4, TCF4, NFIB, NFIX*) that highlight this potential connection. The pairing of similar states also extends to myeloid cells, with Inflammatory-Mac potentially reflecting the hypoxia-associated version of the macrophage program, characterized by upregulation of inflammatory cytokines, downregulation of MHC class II, and an association with CD8+ T cells.

In summary, we provide an extensive spatial description of glioma that demonstrates stereotypical spatial organization at multiple scales, with a prominent role of hypoxia as an organizer and with relative disorganization of regions that lack hypoxia. This adds a spatial dimension to our growing understanding of the glioma ecosystem and may aid the development of novel treatments.

## Author Contributions

A.C.G., N.G.D., R.H., and I.T. conceived the project, designed the study, interpreted results, and wrote the manuscript. C.M., M.C.N., and M.L.S. provided guidance, feedback, and reviewed the manuscript. A.C.G., N.G.D., R.H., D.S., and T.T. performed computational analyses. N.G.D. developed computational methods for spatial analysis. L.N.G.C., G.M., M.C.N., M.W. and M.L.S. provided glioma samples for spatial transcriptomics and spatial proteomics. A.C.G. performed spatial transcriptomics. R.H. performed spatial proteomics. C.M. annotated histology sections. M.N. performed experiments. D.H. and B.L. sectioned tissues and provided histology expertise. M.K. and H.K.S. provided genomics expertise and Visium support. I.G. and Y.A. provided imaging and microscopy expertise and CODEX support. I.T. supervised the study.

## Acknowledgements

This work was supported by the European Research Council Consolidator Grant (to I.T.) and a Broad Institute-Israel Science Foundation Collaborative Project Award (to I.T. and M.L.S.). I.T. is the incumbent of the Dr. Celia Zwillenberg-Fridman and Dr. Lutz Zwillenberg Career Development Chair, and is supported by the Zuckerman STEM Leadership Program, the Mexican Friends New Generation, the Benoziyo Endowment Fund, and the Israel Cancer Research Fund. R.H. is funded by the Walter Benjamin Programme from the German Research Foundation. N.G.D. is supported by the Israeli Council for Higher Education (CHE) via the Weizmann Data Science Research Center and by a research grant from the Estate of Tully and Michele Plesser. The authors would like to thank Dr. Ofra Golani for helpful discussions regarding image analysis and cell segmentation and Rony Chanoch-Myers and Rotem Tal for helpful feedback on the manuscript. The authors would like to thank Abcam for the generous donation of carrier-free antibodies for CODEX experiments.

## Experimental Methods

### Patient samples

Tumor samples used for Visium spatial transcriptomics and CODEX were obtained from patients undergoing tumor resection at University Hospital Zurich, Zurich, Switzerland (ZH samples), Massachusetts General Hospital, Boston, MA (MGH samples), and Brigham and Women’s Hospital, Boston, MA (BWH samples) carried out in accordance with approved guidelines and with patient written consent under ethics approval KEK-ZH-Nr. 2015-0163, University Hospital Zurich and IRB #1360-1, Weizmann Institute of Science. The clinical characteristics of the patient cohort are detailed in Table S1. Tumors ZH1007, ZH1019, ZH881, ZH916, and ZH1041 were spatially annotated by the surgeon during navigated-guided surgery. In these cases, multiple samples were collected from different regions of the same tumor annotated as *necrotic*, *T1 contrast-enhancing*, *infiltrating*, or *bulk*.

### Sample Preparation

Samples were either flash frozen by liquid nitrogen and embedded in cold OCT on dry ice (Scigen OCT Compound, #4586) when already frozen or embedded in OCT at the time of freezing. The RNA quality of each sample was evaluated by Tapestation (Tapestation RNA Screen Tape, Agilent) after isolating RNA (Zymo Quick RNA MicroPrep Kit, #ZR-R1051) from multiple tissue sections. Samples with RIN values >7 were profiled for spatial transcriptomics. Fresh frozen samples were sectioned at 10uM thickness with a cryostat onto 10X Visium Spatial Transcriptomics slides (Visium Spatial Gene Expression Slide and Reagent Kit, PN-1000184) with spatially barcoded capture areas according to the manufacturer’s instructions. Tissues sectioned onto Visium slides were profiled either on the same day as sectioning or within 1 week following storage at -80C.

### H&E staining and imaging

Tissue sections on Visium slides were first fixed in methanol (Millipore Sigma #34860) followed by an aqueous eosin-based H&E protocol according to manufacturer’s instructions (10X Visium Methanol Fixation, H&E Staining, and Imaging Protocol CG000160). Brightfield imaging was performed using a wide-field Leica DMIi8 inverted microscope (Leica-microsystems CMS GmbH Germany) equipped with a DFC310FX color camera. Images were acquired with a 10x/0.25 dry objective and stitched by Leica Application Suite X software. Image post-processing was performed using Fiji version 2.3.1.

### 10X Visium cDNA synthesis and library generation

Following imaging of H&E staining, permeabilization was carried out on the Visium slide to capture mRNA released from the tissue. The optimal permeabilization time (9 minutes) was determined by a permeabilization time course experiment (i.e., Visium tissue optimization experiment). cDNA synthesis and library generation were performed with the Visium Spatial Gene Expression Slide and Reagent Kit (10X Genomics). Briefly, reverse transcription was performed by a template switch oligo protocol to generate a second strand and cDNA synthesis was carried out according to qPCR results (KAPA FAST SYBR qPCR master mix, KAPA Biosystems). The amplified cDNA was fragmented, end repaired, ligated with index adaptors and size selected with cleanups between each step using the SPRIselect Reagent kit (Beckman Coulter). Quality control and quantification of the resulting dual-indexed barcoded libraries was performed with Agilent TapeStation and by qPCR (NEBNext Library Quant Kit for Illumina, New England Biolabs).

### Sequencing

QC of final libraries was perfomed by Tapestation and paired-end dual indexed final libraries were diluted to 1.8nM, pooled, and denatured prior to sequencing on Novaseq (Illumina) using the Novaseq SP 100 cycles sequencing kit (Illumina) with 1% PhiX and the following sequencing parameters: Read 1 – 28 cycles, Read 2 – 90 cycles, Index 1 – 10 cycles, Index 2 – 10 cycles.

### CODEX antibodies and imaging

A list of all antibodies can be found in Table S3. All antibodies not commercially available from Akoya and hence requiring custom-conjugation with DNA-barcodes were first validated with conventional immunofluorescence (**Supplementary Note 2**). Control tissues included healthy duodenum for immune markers, carcinosarcoma for p53, and adequate GBM samples containing the respective cell types and states as identified by Visium. Fresh frozen sections were fixed with 4% PFA for 20 min and blocked with 4% BSA and 0.25% TritonX-100 for 30 min. Primary antibodies were incubated overnight at 4 °C and secondary antibodies incubated for 2h at room temperature. All antibodies (including pre-conjugated antibodies from Akoya) were additionally validated on the respective GBM control samples using multiple CODEX runs to adjust antibody concentration and exposure times to optimize signal to noise ratio. CODEX runs were performed using PhenoCycler-Fusion and imaging with PhenoImager (both Akoya Biosciences) according to manufacturer instructions. After the CODEX run, hematoxylin and eosin staining was performed inside the flow cell using a syringe.

## Computational Methods

### Spatial Transcriptomics (Visium) Analysis

### Alignment, processing, and quality control of Visium data

Alignment of FASTQ files with human reference genome GRCh38, UMI counting, and spot barcode filtering were performed using SpaceRanger (versions 1.0 and 1.1, 10x Genomics). Alignment between positionally barcoded Visium spots and tissue images to obtain spatial coordinates necessary to generate spatial maps was performed using Loupe Browser (version 5.0.1, 10x Genomics). Expression levels were quantified as *E_i_*_,*j*_ = log_2_ (1 + CPM*_i_*_,*j*_/10), in which counts per million (CPM)*_i_*_,*j*_ refers to 10^6^ × UMI*_i_*_,*j*_/sum(UMI_1…*n*,*j*_), for gene *i* in sample *j*, with *n* being the total number of analyzed genes. The average number of UMIs detected per spot was less than 100,000; thus, CPM values were divided by 10 to avoid inflating the differences between detected (*E_i_*_,*j*_ > 0) and undetected (*E_i_*_,*j*_ = 0) genes as previously described.^36^

For each spot, the number of counts was used as a proxy for sample quality. Spots with fewer than 1000 counts and/or expressing more than 20% mitochondrial genes, another proxy for low quality, were filtered out. The top 7,000 most highly expressed genes were retained and centering was performed per sample in order to define relative expression values by subtracting the average expression of each gene *i* across all *k* spots: *Er_i_*_,*j*_ = *E_i_*_,*j*_ − average(*E_i_*_,1…*k*_), where *Er* represents relative expression values.

### Per sample clustering

We applied two clustering approaches to individual samples in order to capture discrete patterns of variation (Leiden clustering) and continuous patterns of variation (NMF). For each sample, following PCA, Leiden clustering was performed on the SNN graph (implemented with Seurat version 4.3.0). Gene programs were defined per Leiden cluster by differential expression analysis based on the top 50 most differentially expressed genes by the Wilcoxon Rank Sum Test with a *p* value of <0.005. For tumors in which we had profiled multiple tissue sections from different regions, we also performed per tumor Leiden clustering in which the expression matrices from the individual tissue sections were merged and jointly clustered.

We performed NMF on each sample separately to capture continuous patterns of gene expression variation. Negative values in each centered expression matrix were transformed to zero. To minimize the influence of selection of an individual *k* parameter, we ran NMF with multiple *k* values ranging from 2-11, generating 65 NMF programs per sample. Each NMF program was summarized by the top 50 genes based on NMF score.

### Generating consensus metaprograms (MPs)

To integrate across samples, we generated metaprograms – consensus gene signatures corresponding to a cell state or cell type. To identify robust gene programs across samples, we jointly clustered both the per sample NMF and Leiden gene programs collected from all GBM samples by their overlap (intersection/union, i.e., Jaccard index). Given the high number of NMF programs, programs included in the clustering were limited to those that were both robust within a sample (observed at multiple values of *k*), non-redundant (for a group of highly similar programs within a sample only one is retained), and similar to programs generated from other samples (i.e., robust across samples).

The clustering process used to define metaprograms was carried out as described in Gavish et al., 2023^14^ as follows: each robust program was compared to all other robust programs to assess the degree of gene overlap between programs. Considering overlap instances of at least 12 genes, the programs with the maximal number of overlaps was selected as a potential founder of a new cluster. If the number of overlapping programs (>12 genes) exceeded four instances, the program with the highest gene overlap to the founder program was added, and thus a cluster was formed. The MP for the cluster was initially defined by the genes that appeared in both programs such that each metaprogram reflects the most robust genes within each cluster of programs. By this approach, 13 clusters corresponding to 13 MPs were derived from all per sample GBM programs.

In two cases where we manually observed clear subclusters, we further split a single MP cluster to two clusters by hierarchical clustering: cluster 5 and cluster 7 (one subcluster of cluster 7 was excluded due to strong enrichment of mitochondrial and ribosomal genes suggesting low data quality). We additionally generated an extended version of each MP (100 genes); however, the extended MPs were not used for the main analysis. The MP generation process described above was performed separately for the IDH mutant samples resulting in 7 IDH mutant MPs. The Vasc, Mac, and Inflammatory-Mac MP clusters were also further subclustered by hierarchical clustering to identify cell subtype signatures. Consensus gene programs were derived from the gene program subclusters selecting for the top 50 most recurring genes per subcluster.

### Gene set enrichment analysis

MPs typically represented a single cell state or cell type. We assessed the enrichment of MP signatures with Gene Ontology terms (MSigDB modules H, C2, C5, C8) as well as published signatures for glioma cell states,^4, 17, 35^ nonmalignant brain cell types,^33, 34^ and pan-cancer cell states and types^14^ by hypergeometric test (FDR adjusted *p*<0.01 was considered significant).

### MP correlations between platforms

In order to directly compare the core GBM malignant cell states (MES1, MES2/MES-Hyp, OPC, NPC, and AC) across platforms (ST=spatial transcriptomics, SN=single-nuclei RNA-Seq, SC=single-cell RNA-Seq), correlations were calculated between each spatial MP gene and the MP score of the analogous SC^4^ or SN (Spitzer et al., unpublished) MP in the respective SC or SN dataset. In order to derive truly single-cell equivalent MPs, a filtering step was added in order to remove genes that were more correlated with another single-cell MP than the MP it currently belonged to according to the following criteria: i)The gene has a correlation below 0.3 for the spatial MP it currently belongs to ii) The gene has a higher correlation with a single-cell program different from the MP it belongs to iii) The difference between the correlation of the best-matching single-cell program and the MP it belongs to is >0.2.

### Spot scoring and assignment of spots to MPs

Spots were scored for MPs as previously described using the scalop R package (https://github.com/jlaffy/scalop).^4,5^ Given a set of genes (*Gj*) reflecting an MP corresponding to a cell state or cell type, we calculate for each spot *i*, a score, *SCj(i),* quantifying the relative expression of *Gj* in spot *i*, as the average relative expression (*Er*) of the genes in *Gj*, compared to the average relative expression of a control gene set (*Gj cont*): *SCj (i*) = average[*Er(Gj,i)]* – average[*Er(Gj cont,i*)]. The control gene-set is defined by first binning all analyzed genes into 30 bins of aggregate expression levels (*Ea*) and then, for each gene in the gene set *Gj*, randomly selecting 100 genes from the same expression bin. In this way, the control gene set has a comparable distribution of expression levels to that of *Gj*, and the control gene set is 100-fold larger, such that its average expression is analogous to averaging over 100 randomly selected gene sets of the same size as the considered gene set. Spots were assigned to the MP for which it scored most highly. For the analysis in which single cells from Neftel et al. 2019 were scored for the spatial MPs, a stricter approach was used for assignment of MPs to cells. In this case, cells needed to have a score ≥ 1 for the highest-scoring MP and a difference of at least 0.1 between the highest and second-highest MP scores in order to be confidently annotated with that MP.

Due to the relatively low number of IDH mutant samples (*n*=6) and corresponding MPs generated from them when they are considered separately (*n*=7), when assigning IDH mutant spots to MPs we integrated the IDH mutant MPs with GBM MPs according to the following procedure. First, we scored IDH mutant spots for the GBM MPs with the following thresholds: if an IDH mutant spot scores highest for a GBM MP, in order for it to be assigned to a GBM MP, it must have a minimum score of at least 1.5 for the MP and there must be a difference of at least 0.2 between the highest and second-highest MP score. If there is no equivalent IDH mutant MP, and at least 5% of spots could be confidently assigned to the GBM MP, then that GBM MP was added to the list of IDH mutant MPs for scoring and assignment of IDH mutant spots. By this criteria, the following GBM MPs were added to the IDH mutant MP list: Prolif-Metab, Reactive-Ast, Inflammatory-Mac, MES-Hyp, and Chromatin-Reg.

### CNA inference

CNAs were estimated similarly to as described previously.^6, 37^ We used an increased size sliding window of 150 genes as ST data is noisier than single cell data, and an external Visium normal brain reference from Ravi et al. 2022 to define a baseline of normal karyotype. We then scored each spot for ‘‘CNA signal’’, defined as the mean of the absolute of CNA values across regions with high CNAs, and ‘‘CNA correlation’’ which refers to the correlation between the CNA profile of each spot and the average CNA profile of all spots of the same sample. Following this we reclassified spots using the two measurements, and spots within the query sample with low scores were integrated with the external reference. Finally, we performed another iteration after which we scored again each spot for CNA signal and correlation. To infer spots and MP malignancy level we used the CNA correlation score plus a scaled CNA signal score: 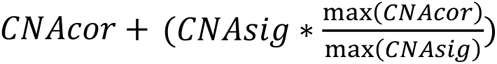. We then performed a min-max scaling for the final score.

### Spatial coherence score

The spatial coherence score of a specific MP in a sample is defined as the scaled average number of immediate neighbors of the same MP across all MP spots. To scale the observed spatial coherence score we shuffle the spots positions 100 times and perform a similar calculation over the shuffled samples, averaging across the results to generate an expected minimum value. We also define a maximum expected value by modeling the average number of immediate neighbors of the same MP where all the MP spots are organized in a regular hexagon pattern. The final score is derived as follows: 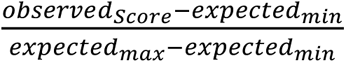

### Defining organizational zones

In order to assign each spot as *structured* or *disorganized* we first generate subsamples of three different sizes (radii 5, 8, 11) surrounding each spot. Next, we calculated the spot’s spatial coherence score in each subsample by averaging the spatial coherence of all MPs in the subsample. Each spot is defined as *structured* if it scores higher than the 40% quantile (across all spots from all samples) for each of the three different window sizes, and *disorganized* if it scores lower than the 40% quantile for each of the three different window sizes. Otherwise, the spot is defined as intermediate. To smooth out the resulting regions, we set a sliding window with *r*= 4 surrounding each spot and make a final region assignment based on the assignment of the majority of spots, to either *structured, disorganized* or intermediate, within the window. After the initial division to *structured* and *disorganized* regions, we performed a second division by malignancy level of the spots. We inferred the spots malignancy level based on their CNA as described above. Finally, we scaled the score to range from 0 to 1. Spots that were classified as *structured* and had a malignancy score ≥ 0.4 were classified as *Structured-Malignant* and those with a malignancy score <0.4 were classified as *Structured-Normal*. Spots that were classified as disorganized and had a malignancy score ≥0.5 were classified as *Disorganized-Malignant* and those with a malignancy score <0.5 were classify as *Disorganized-Normal*.

### Spot Colocalization

We first performed a scoring-based deconvolution (per sample) on all spots by scoring spots for MPs as described in ‘Spot scoring and assignment of spots to MPs’. For each spot, all MP scores above 0.1 were considered as presence of the MP in a spot and deconvolution matrices were binarized such that for a given spot, presence of MP=1 and absence of MP=0. The following analysis was performed separately on spots in structured and disorganized regions per sample. First, for MP *a* and MP *b*, we compute the proportion of spots containing *a* or *b* (denoted as *Ta* or *Tb*). Second, we compute *N*, the number of times *a,b* co-occur within the same spot. Finally, we compute a colocalization value defined as *N/ Ta+Tb*. After calculating the mean colocalization of each MP pair per sample (the observed colocalization value), an expected colocalization value was computed by performing the measure on 500 shuffled deconvolution matrices per sample. A *p* value was then computed by Fisher’s exact test. The per sample effect size was defined as *expected/observed* per MP. Colocalization of each MP pair per sample was defined as significant if the effect size was ≥1.3 and the *p* value was ≤0.01. Finally, the proportion of samples for which a MP pair was considered significant was calculated to consider robustness across samples. For downstream analysis, the mean colocalization value per pair and the proportion of samples for which it was significantly colocalized were used.

### Spot Adjacency

Spot adjacency was calculated per sample. The observed asymmetrical adjacency between MP *A* and MP *B* within a given sample is defined as MP *A* number of immediate MP *B* neighbors out of all MP A neighbors: 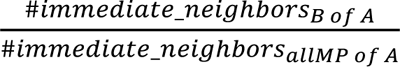

In order to regress out the spatial coherence effect of MP *B* on its adjacency, we normalize this score by the *relative adjacency capacity* of MP *B,* i.e., the number of MP *B* spots with non-MP *B* neighbors out of all MP adjacency capacities (except for MP *A*): 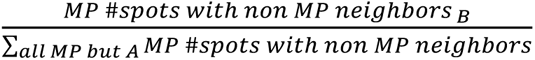

Next, we shuffle the spots positions 10,000 times and calculate the expected adjacency scores to create a null distribution. The *p* value is computed by the number of times the *observed* score was higher/lower than the shuffled scores (the smaller of the two).

### Regional Composition

Regional composition was computed per sample. For each radius (defined as the number of hexagons circling a spot) ranging from 1 to 15, we defined a window around each spot. We then calculated the abundance of all MPs in the window and finally, calculated the correlations of all pairs’ abundance across all the windows of the same size, omitting windows that included only one MP. We then repeated the process over 500 shuffled permutations of the spots positions to generate a random distribution, based on which we calculated significance for the correlation by Fisher’s exact test.

### Robust couplings across analyses and scales (consensus interactions)

To be able to compare scores of pairs between analyses, we scaled the scores of all analyses by a min-max scaling so that all scores range between 1 to -1. For each analysis after scaling, a MP pair that exhibits a mean scaled score above 0.35 across samples and was significantly connected in at least 20% of samples was considered strongly connected. A MP pair was defined as consistently strongly connected across analyses (i.e., consensus interaction) if it was strongly connected within at least 2 out of 5 analyses (colocalization, adjacency, regional composition with window sizes *r*= 5,8,15) and in at least one of the measures of direct coupling: *colocalization* or *adjacency*. By this definition, 21 structured pairs were consistently strongly connected compared to only 2 pairs in disorganized regions.

### Generation of spatial layers network graph

We generated a graph of robustly coupled pairs in structured regions across analyses with Cytoscape (version 3.9.1) by the following criteria: pairs that are defined as “strongly connected” in any 2/5 measures of spatial relationships. By this definition, we include an additional 4 pairs beyond the 21 pairs initially defined as robustly coupled. All MPs except for the Chromatin-Reg MP could be confidently assigned to layers. Chromatin-Reg is unassigned in the layers model due to its internal inconsistency in coupling between measures in the Visium analysis and inconsistency between the Visium and CODEX analysis.

### MP frequency in infiltrative vs. non-infiltrative environments

We defined a spot to be in an infiltrative environment if 30% of the spots surrounding it, within a window of a radius of 4 spots, are annotated as normal brain (i.e., Neuron, Oligo, or Reactive-Ast). Next, we measured the overall frequency of each MP in the two environments (infiltrative and non-infiltrative) across all the samples.

### T cell inference

To infer the presence of T cells within spots, we lowered the UMI count QC filter to 150 to include low complexity spots due to the association between T cells and low RNA content/low complexity. Spots that include at least two counts for at least two canonical T cell markers (CD2, CD3E, CD3G, CD3D, FOXP3, CTLA4, CD28) were identified as putative T cell containing spots. Putative T cell-containing spots with at least one CD8A or CD8B count and more CD8A/CD8B counts than CD4 counts were further classified as CD8 T cell-containing and spots containing at least two CD4 counts and more CD4 counts than CD8A or CD8B counts were further classified as CD4 T cell containing spots with the caveat that macrophages can also express CD4.

### Cell cycle inference

Using the cell cycle metaprogram gene signatures obtained from Neftel et al. 2019 we scored each spot per sample for “g1_s” and “g2_m” programs and maintain the higher of the two as the spot cell cycle signal. We set a threshold of 0.85 for both GBM and IDH-mut samples and in both cases this corresponds to 90% to 95% quantiles, above which a spot was classified as cycling. Next, we calculated the frequency of spots classified as cycling within each MP. The results remain overall consistent within each group at different thresholds.

## CODEX analysis

### Segmentation, data preprocessing, and phenotyping of CODEX data

Nuclear segmentation was performed with the StarDist^38^ plugin within QuPath^39^ applying the pre-trained model (dsb2018_heavy_augment.pb) to the DAPI channel. Areas with staining or tissue artifacts (i.e. tissue folds, freezing artifacts) were excluded manually from segmentation and further analysis. Nuclei were expanded by 3 µm to get conservative estimates for cell boundaries. Mean fluorescence intensity for every marker and cell compartment, together with morphology quantifications (area, solidity, circularity) were exported and further analyzed in R. Data pre-processing included log2-transformation and per sample z-scoring. Phenotyping was done per sample by over-clustering using PhenoGraph^40^ and calculating differentially expressed proteins per cluster. Clusters were named based on their differentially expressed protein expression and visual inspection in QuPath. Mixed or ambiguous clusters were further sub-clustered. In cases where the sub-clustering did not derive clean clusters, cells were annotated as “unknown” and excluded from further analysis. Sub-clustering for macrophages, vascular and T cells were performed across the entire cohort only using their differentially expressed proteins or known functional markers.

### Creation of pseudospots

We created and overlaid a virtual tissue microarray (TMA) grid corresponding to the Visium spot properties (diameter=55µm, centroid distance=100 µm) to every sample in QuPath. Cells were associated to pseudospots if their nuclear segmentation mask centroid was within the pseudospot boundaries. We then view the true MP composition of each pseudospot. The identity of a pseudospot was defined by the dominant MP within each pseudospot. To estimate the level of homogeneity of each pseudospot, we visualize the density of the abundance of the most abundant MP within each pseudospot vs. the second most highly abundant MP (Fig. S2D, Fig. 3H).

### Defining organizational zones

To define spatial patterns (*structured/disorganized*) in the CODEX samples we used CODEX pseudospots and performed a similar process to Visium with *r*=5,8 and a similar threshold of 40%.

### Minimum distance to vascular cell

We created for every sample a distance matrix using the x- and y-coordinates of the segmentation masks centroids and calculated the minimum distance from every cell type to the closest vascular cell.

### Colocalization

We created a neighborhood matrix for every sample using the imcRtools package in R. In the neighborhood matrix rows represent cells in a sample and columns represent the count of every cell type in that neighborhood. The neighborhood for every cell was defined by a radius of 27.5um (coloc1: corresponding to the dimensions of 1 visium spot) or 127.5um (coloc3: corresponding to the dimension of 3 visium spots). We then calculated the mean interaction count between every possible cell type pair. Example: For *cell type A* we calculated the sum of counts that *cell type B* is in the neighborhood of *cell type A* and divided this value by the count of *cell type A* in a given sample. To focus on co-localizations between different cell types (rather than emphasizing self-pairs), we excluded neighborhoods that were dominated by a cell type with a fraction > 0.8. This correction was important to detect neighboring cell types (i.e. MES) next to cell types with very high spatial coherence (i.e. MES-Hyp). Next, we shuffled cell type labels n=500 times and calculated z-scores from the observed and shuffled interaction counts. Significance was reached if 95% of shuffled z-scores were larger or smaller (depending on directionality: coupled vs. uncoupled) than the observed value.

### Regional composition

CODEX regional composition was calculated similarly to Visium regional composition on the single cell level, using the same radii as used for Visum. For each radius *i* (defined as the distance between *I* number of spots) ranging from 1 to 15, we defined a window around each cell. We then calculated the abundance of all MPs in the window and finally, calculated the correlations of all pairs’ abundance across all the windows of the same size, omitting windows that included only one MP. We then repeated the process over 500 shuffled permutations of the spots positions to generate a random distribution, based on which we calculated significance for the correlation by Fisher’s exact test.

