## Supplementary Information for "Integrative spatial analysis reveals a multi-layered organization of glioblastoma"

**Table S1: Overview of patient cohort**

| Sample-ID | Patient-ID | Region | Tumor | Histology | Grade | MGMT | Location | Age | Sex | Visium | CODEX |
|---|---|---|---|---|---|---|---|---|---|---|---|
| MGH258 | MGH258 | bulk | IDH WT | GBM | 4 | methylated | left frontal | F | 43 | 1 | 1 |
| ZH881_1A | ZH881 | bulk | IDH WT | GBM | 4 | partially methylated | right temporal | M | 79 | 1 | |
| ZH881_1B | ZH881 | bulk | IDH WT | GBM | 4 | partially methylated | right temporal | M | 79 | 1 | 1 |
| ZH881_2 | ZH881 | bulk | IDH WT | GBM | 4 | partially methylated | right temporal | M | 79 | 1 | |
| ZH881_T1 | ZH881 | T1 | IDH WT | GBM | 4 | partially methylated | right temporal | M | 79 | 1 | 1 |
| ZH881_inf | ZH881 | inf | IDH WT | GBM | 4 | partially methylated | right temporal | M | 79 | 1 | 1 |
| ZH916_bulk | ZH916 | bulk | IDH WT | GBM | 4 | methylated | left temporal | M | 62 | 1 | |
| ZH916_T1 | ZH916 | T1 | IDH WT | GBM | 4 | methylated | left temporal | M | 62 | 1 | 1 |
| ZH916_inf | ZH916 | inf | IDH WT | GBM | 4 | methylated | left temporal | M | 62 | 1 | 1 |
| ZH1007_nec | ZH1007 | nec | IDH WT | GBM | 4 | NA | NA | NA | NA | 1 | 1 |
| ZH1007_inf | ZH1007 | inf | IDH WT | GBM | 4 | NA | NA | NA | NA | 1 | 1 |
| ZH1007_bulk | ZH1007 | bulk | IDH WT | GBM | 4 | NA | NA | NA | NA | | 1 |
| ZH1019_T1 | ZH1007 | T1 | IDH WT | GBM | 4 | NA | NA | NA | NA | 1 | 1 |
| ZH1019_inf | ZH1007 | inf | IDH WT | GBM | 4 | NA | NA | NA | NA | 1 | 1 |
| ZH1041_T1 | ZH1041 | T1 | IDH WT | GBM | 4 | NA | NA | NA | NA | | 1 |
| BWH23 | BWH23 | bulk | IDH-mut | ODG | 4 | unmethylated | left frontal | M | 32 | 1 | |
| BWH24 | BWH24 | bulk | IDH-mut | AA | 4 | methylated | right parietal | F | 54 | 1 | |
| BWH25 | BWH25 | bulk | IDH-mut | AA | 4 | methylated | bifrontal | F | 51 | 1 | |
| BWH28 | BWH28 | bulk | IDH-mut | AA | 2/3 | partially methylated | right frontotemporal | M | 26 | 1 | |
| MGH259 | MGH259 | bulk | IDH-mut | ODG | 3 | partially methylated | left frontal | F | 40 | 1 | |
| BWH35 | BWH35 | bulk | IDH-mut | ODG | 2 | methylated | right parietal | F | 36 | 1 | |

bulk: tumor bulk, T1: T1-contrast enhancing, inf: infiltrating, NA: not annotated

### Table S2: Metaprograms

#### GBM metaprograms

| Neuron | Vasc | MES.Hyp (malig.) | Mac | Oligo | MES (malig.) | Metabolism ( | MES.Ast (malig.) | Reactive.Ast | NPC (malig.) | inflammatory.resp | Chromatin.reg (malig.) | OPC (malig.) | AC (malig.) |
|---|---|---|---|---|---|---|---|---|---|---|---|---|---|
| SNAP25 | VWF | VEGFA | C1QB | PLP1 | EFEMP1 | GGCX | METTL7B | MGST1 | INA | CCL2 | ATRX | SIRT2 | BCAN |
| SYT1 | COL4A1 | NDRG1 | C1QA | MAG | CHI3L1 | LRRC75A | GPC1 | S100A13 | NELL2 | CCL3 | BPTF | SOX10 | MYBPC1 |
| VSNL1 | COL4A2 | ADM | FCER1G | CLDND1 | CD44 | IQCK | CXCL14 | FXYD1 | NEUROD2 | CCL4 | ZBTB20 | BCAN | S100B |
| SLC17A7 | PDGFRB | IGFBP5 | C1QC | TF | C1R | TRA2A | LPL | MT1M | NSG2 | CCL4L2 | ANKRD12 | BCAS1 | ALDOC |
| RTN1 | COL1A2 | MT1X | LAPTM5 | CLDN11 | MGST1 | CTNNB1 | MOXD1 | MT-ND4L | SYT4 | DNAJB1 | CCDC88A | GPR17 | AQP1 |
| NRGN | ENG | SLC2A1 | CD14 | ENPP2 | LTF | HNRNPH1 | POSTN | MT1E | ZBTB18 | EGR1 | EIF5B | PLP1 | TTYH1 |
| TAGLN3 | FN1 | BNIP3 | CD74 | CNP | ANXA1 | SYNC | RCAN1 | S100A1 | CDK5R1 | IL1B | HBA2 | MBP | ATP1A2 |
| CCK | RGS5 | PGK1 | SRGN | ERMN | C1S | ATP1B1 | TNFRSF12A | SLC1A2 | DCX | NR4A1 | MALAT1 | OLIG1 | AQP4 |
| CHN1 | COL18A1 | ANGPTL4 | AIF1 | PPP1R14A | CHI3L2 | C1orf56 | FABP5 | ADIRF | NSG1 | CD83 | MAN1A2 | OLIG2 | TSC22D4 |
| ENC1 | COL3A1 | HILPDA | FCGR3A | TMEM144 | SOD2 | GOLIM4 | FABP7 | AGT | RUNX1T1 | FOS | NFIX | SCRG1 | GPR37L1 |
| STMN2 | BGN | ERO1A | TYROBP | MOG | AQP1 | PHKG1 | HOPX | CLU | STMN2 | BTG2 | DST | SOX8 | MLC1 |
| DNM1 | ITM2A | NRN1 | HLA-DRB1 | RNASE1 | TNC | AC018557.1 | MT2A | DTNA | BASP1 | CCL3L1 | HBA1 | CNP | C1orf61 |
| UCHL1 | CLDN5 | AKAP12 | CSF1R | SELENOP | ID3 | ARGLU1 | PDLIM4 | HLA-DRA | BHLHE22 | CH25H | HBB | TUBB4A | CST3 |
| NEFL | HSPG2 | LGALS3 | HLA-DRA | CNTN2 | MAOB | MT-ND6 | PTN | LINC00844 | CAMK2B | CXCL10 | MT-ATP8 | DLL3 | NDRG2 |
| OLFM1 | ESAM | SLC2A3 | TREM2 | MAL | CCL2 | ACOT13 | F3 | PTGDS | CXADR | CYR61 | NEAT1 | HES6 | LINC00844 |
| SYN1 | MGP | LDHA | APOC1 | MBP | CP | GATC | GAP43 | AHCYL1 | LY6H | DUSP1 | NFIC | MARCKSL1 | PLP1 |
| SYN2 | MYL9 | ENO2 | C3 | PTGDS | F3 | LIG3 | KCNF1 | AQP4 | NREP | GBP1 | TTC3 | SLC44A1 | FADS2 |
| CHGA | NDUFA4L2 | GAPDH | HAMP | MOBP | MAN1C1 | CCDC152 | KLHDC8A | CSRP1 | RAB3A | HSPA1A | ZC3H13 | VCAN | HEPACAM |
| STXBP1 | DCN | FAM162A | ITGB2 | CARNS1 | NAMPT | NME7 | LGALS3 | FAM107A | REEP1 | HSPA1B | AKAP9 | AC009041.2 | SLC1A2 |
| BASP1 | IGFBP4 | CA12 | VSIG4 | EVI2A | S100A10 | STARD7 | PDPN | GJA1 | SNAP25 | HSPA6 | CCNI | CLDN11 | FAM107A |
| GNG3 | EGFL7 | DDIT4 | ALOX5AP | CNDP1 | AL049839.2 | EIF2S3 | SERPINE1 | GSN | SYT1 | HSPH1 | GOLGA4 | DBI | GFAP |
| NAPB | ITGA1 | LOX | FCGBP | ANLN | ATP1B2 | EIF5A | FJX1 | HLA-DPB1 | TUBB4A | IER2 | HIST1H4C | ERBB3 | GJA1 |
| YWHAH | ANGPT2 | IGFBP3 | SPP1 | EDIL3 | C3 | KIF21A | SCG2 | HLA-DRB1 | UCHL1 | IER5 | KMT2E | FIBIN | HES6 |
| SERPINI1 | GNG11 | S100A10 | CD163 | LHPP | CLU | MAP3K12 | SPOCD1 | IFITM3 | ATP1A3 | IRF1 | MTRNR2L12 | GNB4 | ID3 |
| BEX1 | NOTCH3 | SERPINE1 | HLA-DPA1 | PIP4K2A | GAP43 | SLC16A1 | AGT | MALAT1 | PCP4 | JUN | MTRNR2L8 | S100B | METTL7A |
| SNCA | COL1A1 | IGFBP2 | CD68 | SEPT4 | SERPING1 | SOX9 | CHI3L1 | METTL7A | KIF21B | JUNB | PPIG | SNX22 | DDR1 |
| LY6H | CTGF | NUPR1 | S100A8 | UGT8 | RCAN1 | YWHAG | CHPF | MT-ND5 | EPHA5 | RGS1 | PRRC2C | TCF12 | ADCYAP1R1 |
| PHYHIP | NID1 | ENO1 | HLA-DRB5 | APLP1 | ZFP36 | AC009133.1 | DNAJB1 | MYBPC1 | CAMKV | ZFP36 | ELAVL3 | TSC22D4 | ADGRG1 |
| ATP1A3 | TAGLN | CAV1 | MS4A6A | HSPA2 | ACTN1 | CDC42SE1 | ELOVL2 | S100A10 | RPRM | AHRR | MAP4K4 | TTYH1 | AGT |
| ATP6V1G2 | ACTA2 | HSPA1B | RGS1 | MYRF | GJA1 | F3 | GADD45A | AL078639.1 | CDKN2D | EGR3 | NFIB | MYRF | FAM181B |
| NSF | IGFBP7 | MT2A | RNASET2 | GSN | IGFBP7 | HEPN1 | GDF15 | ATP1A2 | Mar-04 | CLPTM1L | PLCG2 | PLPPR1 | HOPX |
| THY1 | PECAM1 | PLOD2 | S100A9 | QDPR | LRIG1 | MMACHC | IGFBP2 | ATP1B2 | STX1A | TAP1 | PLEKHA4 | PTPRZ1 | MBP |
| ATP1B1 | PLVAP | BNIP3L | HLA-DPB1 | SLAIN1 | S100A6 | PPP1CB | IGFBP3 | CD44 | TUBB3 | STAT1 | RSF1 | SMOC1 | RGMA |
| SYP | AGRN | CA9 | CTSS | KLK6 | NTRK2 | SUMF2 | LMO2 | CD74 | MPPED1 | SERPINE1 | SPECC1 | CADM2 | CSPG5 |
| TSPAN7 | A2M | CXCL8 | SLCO2B1 | SCD | SOCS3 | TP53 | LZTS1 | CPE | SCN3B | EXOC3-AS1 | TCAF1 | CNTN1 | ENHO |
| TUBB2A | SERPINH1 | DNAJB1 | CD53 | ABCA2 | WLS | AC011603.2 | MT3 | GFAP | TMEM35A | GADD45B | APC | CSPG5 | CKB |
| PRKAR1B | CD93 | BHLHE40 | CTSC | PLA2G16 | FOS | ARPC4 | NAMPT | IFITM2 | BCL11A | SEZ6L | ARHGAP5 | ETV1 | CRISPLD1 |
| TMEM59L | ITGB1 | PLIN2 | HLA-DMB | APOD | EMP1 | BEST1 | PLA2G5 | LGALS3 | OLFM1 | IGFBP3 | BRD4 | FXYD6 | EDNRB |
| CALM3 | MYH9 | CHPF | CHI3L1 | PLLP | KCNN3 | CNTD1 | RAMP1 | MBP | EPHA4 | ATF3 | EEA1 | HMGB2 | GATM |
| NPTXR | MCAM | HMOX1 | HLA-DQB1 | AMER2 | MAPK4 | CTBP1 | S100A16 | MT1F | STMN4 | SLPI | GOLGB1 | MAG | ITM2C |
| TMEM130 | APOLD1 | HSPA1A | S100A11 | ELOVL1 | SCARA3 | FGD5-AS1 | SLC6A11 | MT1G | CD24 | ROPN1L | KTN1 | PTN | PTN |
| MEG3 | HIGD1B | ARRDC3 | SLC2A5 | NKX6-2 | VIM | MARCKS | SLN | NTRK2 | GNAO1 | GPX3 | MT-ND4L | SCD5 | SLC1A3 |
| NELL2 | MIR4435-2H( | HSPA5 | A2M | PLEKHH1 | PDPN | MSH6 | TMEM158 | PLPP3 | Sep-03 | AZGP1 | NCOR1 | TNR | BCAN |
| HPCA | SPARC | SPP1 | EFEMP1 | OPALIN | NNMT | PAQR3 | TTYH3 | S100A6 | TUBB2A | WARS | NFIA | TNS3 | MYBPC1 |
| IDS | EPAS1 | UPP1 | GPNMB | SLC44A1 | TRIM47 | PDGFA | EMP1 | SEC62 | MLLT11 | CCDC127 | PCDH9 | TSPAN7 | S100B |
| AK5 | LAMA4 | ZNF395 | HLA-DQA1 | SPP1 | EFEMP1 | RAB6A | JAG1 | SLC1A3 | JPT1 | G0S2 | SLPI | UGT8 | ALDOC |
| ENO2 | IFI27 | ARL4C | LILRB4 | BCAS1 | CHI3L1 | RPL23 | HMGA1 | SPARC | ENC1 | PLAUR | SOX4 | AC008080.4 | AQP1 |
| NCDN | FSTL1 | HK2 | STAB1 | CSRP1 | CD44 | SPTBN1 | CCL2 | ECEL1 | STMN1 | VGF | TCF4 | ANP32B | TTYH1 |
| SNCB | CYTOR | GLUL | VAMP8 | HHIP | C1R | ZBTB18 | COL9A3 | CHI3L1 | DPYSL3 | CD14 | VCAN | SIRT2 | ATP1A2 |
| CRYM | CD248 | | IBSP | LINC00844 | MGST1 | AC092069.1 | ZYX | AQP1 | INA | APLNR | ZNF91 | SOX10 | AQP4 |

IDH-mutant metaprograms

| macrophage | oligo | vascular | low.quality | AC | OC.NPC1 | neuron |
|---|---|---|---|---|---|---|
| LAPTM5 | CLDN11 | APOLD1 | PLCG2 | GJA1 | HES6 | DNM1 |
| SPP1 | CLDND1 | CLDN5 | MALAT1 | LRIG1 | OLIG2 | ENC1 |
| TYROBP | CNP | FN1 | MTRNR2L12 | AGT | DLL3 | NAPB |
| APOC1 | APOD | IFI27 | MTRNR2L8 | EDNRB | ETV1 | NEFL |
| C1QB | ENPP2 | CTGF | MT-ATP8 | MLC1 | H2AFV | NRGN |
| C1QC | ERMN | ITM2A | MT-ND4L | SFRP2 | PDGFRA | SLC17A7 |
| CD68 | MAG | EPAS1 | AL078639.1 | ATP1A2 | SMOC1 | SNCA |
| CD74 | MAL | GNG11 | MT-ND5 | SLC1A2 | SOX8 | STMN2 |
| FCER1G | MOG | SLC2A1 | MTRNR2L1 | SLC1A3 | AC009041.2 | SYT1 |
| S100A11 | RNASE1 | IFITM3 | AL355075.4 | ALDOC | AC084033.3 | VSNL1 |
| C1QA | SELENOP | MT1E | HBA2 | CPE | AC091138.1 | ATP1B1 |
| C3 | TF | BGN | HBB | ETNPPL | AMOTL2 | RAB3A |
| FTL | TMEM144 | COL1A2 | TAF1D | TTYH1 | B4GALNT1 | RTN1 |
| HLA-DRA | TUBB4A | DCN | MAP2 | ADCYAP1R1 | CCND2 | SNAP25 |
| HLA-DRB1 | ABCA2 | MFSD2A | MT-ND1 | ANOS1 | CD24 | SYN1 |
| NPC2 | APLP1 | VWF | C5orf63 | AQP4 | CDK4 | SYP |
| TREM2 | CARNS1 | A2M | DST | BAG3 | CDKN2A | TAGLN3 |
| A2M | CNDP1 | COL4A1 | EIF4A2 | CDC42EP4 | CDKN2C | UCHL1 |
| CTSB | CNTN2 | COL4A2 | MT-CO1 | DCLK2 | CKS2 | BASP1 |
| CYBA | PLP1 | MT2A | MT-CO3 | DDAH1 | DCTN2 | CHN1 |
| HLA-DPA1 | PPP1R14A | RGS5 | MT-ND2 | DKK 3.00 | DSEL | MAP1LC3A |
| CTSS | ANLN | SLC38A5 | MT-ND4 | GABBR1 | DTX3 | NNAT |
| GPNMB | EDIL3 | ENG | SNHG12 | GPRC5B | FERMT1 | OLFM1 |
| HLA-DPB1 | HSPA2 | HYAL2 | ZBTB20 | ID4 | HMGB2 | PLD3 |
| LGALS1 | RAPGEF5 | IGFBP7 | HBA1 | KCNN3 | JPT1 | PTPRN |
| APOE | SPOCK3 | MYH9 | MAP1B | NMB | MARCH9 | STXBP1 |
| CD14 | MOBP | NDUFA4L2 | MT-ATP6 | NTSR2 | NOVA1 | SYN2 |
| CD53 | MYRF | SLC7A5 | MT-ND3 | NUDT4 | OLIG1 | SYNGR1 |
| CSF1R | PIP4K2A | TIMP1 | NCL | SLC4A4 | OS9 | TAC1 |
| CTSD | SLAIN1 | TMSB10 | NEAT1 | SPARCL1 | PGRMC1 | TUBA4A |
| HLA-B | STMN4 | ABCG2 | RPL27A | TIMP3 | PHLDA1 | TUBB2A |
| ITGB2 | GPR37 | CAVIN2 | RPL39 | WLS | SHD | YWHAG |
| KCTD12 | QDPR | COL3A1 | WDR74 | AQP1 | SOX4 | VGF |
| LGMN | TTYH2 | HLA-B | AD000090.1 | GABBR2 | TCF12 | SCG2 |
| NUPR1 | UGT8 | PECAM1 | CLOCK | ADGRB1 | TOP2A | CCK |
| PSAP | CHADL | SLCO2B1 | MT-CYB | EEPD1 | TRIM28 | NPTX2 |
| SCIN | ELOVL1 | SRGN | CSKMT | EFEMP1 | TSFM | GNG3 |
| SRGN | EVI2A | TAGLN2 | ANKRD12 | NCAN | TSPAN31 | NPTXR |
| TMSB10 | KLK6 | ESAM | CCDC88A | EZR | MBD6 | PHYHIP |
| TMSB4X | MBP | VIM | SLC1A2 | MAOB | GRIA2 | PNOC |
| VIM | PMP22 | CD93 | SOX2 | TRIL | ELMO1 | HPCA |
| ZFP36L2 | SLC12A2 | COL18A1 | RBM25 | PLPP3 | FAM110B | BEX1 |
| GFAP | SORT1 | SERPINH1 | BPTF | ITM2C | SALL3 | NRN1 |
| LYZ | ATP1B1 | MT1M | RPL23A | LIFR | TNR | VAMP2 |
| GRN | PLEKHH1 | IGFBP3 | DYNC1H1 | MGST1 | ASCL1 | SNCB |
| CXCR4 | MTURN | NET1 | PEG10 | BBOX1 | SQLE | PRKAR1B |
| SLCO2B1 | NKX6-2 | MCAM | NRCAM | IGFBP7 | MMP16 | TMEM130 |
| RGS1 | CDKN1C | TIMP3 | SNHG19 | RGMA | SCN3A | SNCG |
| VSIG4 | SLC44A1 | SLC39A10 | COX8A | ATP1B2 | SOX10 | GABARAPL1 |
| GPR34 | Sep-04 | ITIH5 | TNR | FAM213A | APCDD1 | CHGB |

### GBM extended metaprograms

| AC_ext | OPC_ext | chromatin.re | metabolism_ | vasc_ext | neuron_ext | hypoxia_ext | macrophage_ | oligo_ext | mes_ext | ac.mes_ext | reactive.astro | npc_ext | inflammatory.resp_ext |
|---|---|---|---|---|---|---|---|---|---|---|---|---|---|
| BCAN | SIRT2 | ATRX | GGCX | VWF | SNAP25 | VEGFA | C1QB | PLP1 | EFEMP1 | METTL7B | MGST1 | INA | CCL2 |
| MYBPC1 | SOX10 | BPTF | LRRC75A | COL4A1 | SYT1 | NDRG1 | C1QA | MAG | CHI3L1 | GPC1 | S100A13 | NELL2 | CCL3 |
| S100B | BCAN | ZBTB20 | IQCK | COL4A2 | VSNL1 | ADM | FCER1G | CLDND1 | CD44 | CXCL14 | FXYD1 | NEUROD2 | CCL4 |
| ALDOC | BCAS1 | ANKRD12 | TRA2A | PDGFRB | SLC17A7 | IGFBP5 | C1QC | TF | C1R | LPL | MT1M | NSG2 | CCL4L2 |
| AQP1 | GPR17 | CCDC88A | CTNNB1 | COL1A2 | RTN1 | MT1X | LAPTM5 | CLDN11 | MGST1 | MOXD1 | MT-ND4L | SYT4 | DNAJB1 |
| OLIG1 | PLP1 | EIF5B | HNRNPH1 | ENG | NRGN | SLC2A1 | CD14 | ENPP2 | LTF | POSTN | MT1E | ZBTB18 | EGR1 |
| TTYH1 | MBP | HBA2 | SYNC | FN1 | TAGLN3 | BNIP3 | CD74 | CNP | ANXA1 | RCAN1 | S100A1 | CDK5R1 | IL1B |
| ATP1A2 | OLIG1 | MALAT1 | ATP1B1 | RGS5 | CCK | PGK 1.00 | SRGN | ERMN | C1S | TNFRSF12A | SLC1A2 | DCX | NR4A1 |
| AQP4 | OLIG2 | MAN1A2 | C1orf56 | COL18A1 | CHN1 | ANGPTL4 | AIF1 | PPP1R14A | CHI3L2 | FABP5 | ADIRF | NSG1 | CD83 |
| TSC22D4 | SCRG1 | NFIX | GOLIM4 | COL3A1 | ENC1 | HILPDA | FCGR3A | TMEM144 | SOD2 | FABP7 | AGT | RUNX1T1 | FOS |
| GPR37L1 | SOX8 | DST | PHKG1 | BGN | STMN2 | ERO1A | TYROBP | MOG | AQP1 | HOPX | CLU | STMN2 | BTG2 |
| MLC1 | CNP | HBA1 | AC018557.1 | ITM2A | DNM1 | NRN1 | HLA-DRB1 | RNASE1 | TNC | MT2A | DTNA | BASP1 | CCL3L1 |
| NCAN | TUBB4A | HBB | ARGLU1 | CLDN5 | UCHL1 | AKAP12 | CSF1R | SELENOP | ID3 | PDLIM4 | HLA-DRA | BHLHE22 | CH25H |
| C1orf61 | DLL3 | MT-ATP8 | MT-ND6 | HSPG2 | NEFL | LGALS3 | HLA-DRA | CNTN2 | MAOB | PTN | LINC00844 | CAMK2B | CXCL10 |
| CST3 | HES6 | NEAT1 | ACOT13 | ESAM | OLFM1 | SLC2A3 | TREM2 | MAL | CCL2 | F3 | PTGDS | CXADR | CYR61 |
| NDRG2 | MARCKSL1 | NFIC | GATC | MGP | SYN1 | LDHA | APOC1 | MBP | CP | GAP43 | AHCYL1 | LY6H | DUSP1 |
| LINC00844 | SLC44A1 | TTC3 | LIG3 | MYL9 | SYN2 | ENO2 | C3 | PTGDS | F3 | KCNF1 | AQP4 | NREP | GBP1 |
| PLP1 | VCAN | ZC3H13 | CCDC152 | NDUFA4L2 | CHGA | GAPDH | HAMP | MOBP | ITM2C | KLHDC8A | CSRP1 | RAB3A | HSPA1A |
| FADS2 | AC009041.2 | AKAP9 | NME7 | DCN | STXBP1 | FAM162A | ITGB2 | CARNS1 | MAN1C1 | LGALS3 | FAM107A | REEP1 | HSPA1B |
| HEPACAM | CLDN11 | CCNI | STARD7 | IGFBP4 | BASP1 | CA12 | VSIG4 | EVI2A | NAMPT | PDPN | GJA1 | SNAP25 | HSPA6 |
| SLC1A2 | DBI | GOLGA4 | EIF2S3 | EGFL7 | GNG3 | DDIT4 | ALOX5AP | CNDP1 | S100A10 | SERPINE1 | GSN | SYT1 | HSPH1 |
| FAM107A | ERBB3 | HIST1H4C | EIF5A | ITGA1 | NAPB | LOX | FCGBP | ANLN | AL049839.2 | FJX1 | HLA-DPB1 | TUBB4A | IER2 |
| GFAP | FIBIN | KMT2E | KIF21A | ANGPT2 | YWHAH | IGFBP3 | SPP1 | EDIL3 | ATP1B2 | SCG2 | HLA-DRB1 | UCHL1 | IER5 |
| GJA1 | GNB4 | MTRNR2L12 | MAP3K12 | GNG11 | SERPINI1 | S100A10 | CD163 | LHPP | C3 | SPOCD1 | IFITM3 | ATP1A3 | IRF1 |
| HES6 | S100B | MTRNR2L8 | SLC16A1 | NOTCH3 | BEX1 | SERPINE1 | HLA-DPA1 | PIP4K2A | CLU | AGT | MALAT1 | BCL11A | JUN |
| ID3 | SNX22 | PPIG | SOX9 | COL1A1 | SNCA | GPNMB | CD68 | SEPT4 | GAP43 | CHI3L1 | METTL7A | CAMKV | JUNB |
| METTL7A | TCF12 | PRRC2C | YWHAG | CTGF | LY6H | IGFBP2 | S100A8 | UGT8 | SERPING1 | CHPF | MT-ND5 | CD24 | RGS1 |
| DDR1 | TSC22D4 | ELAVL3 | AC009133.1 | NID1 | PHYHIP | NUPR1 | HLA-DRB5 | APLP1 | FCGBP | DNAJB1 | MYBPC1 | CDKN2D | ZFP36 |
| SCRG1 | TTYH1 | MAP4K4 | CDC42SE1 | TAGLN | ATP1A3 | ENO1 | MS4A6A | HSPA2 | RCAN1 | ELOVL2 | S100A10 | CRMP1 | AHRR |
| ADCYAP1R1 | MYRF | NFIB | F3 | ACTA2 | ATP6V1G2 | CAV1 | RGS1 | MYRF | ZFP36 | GADD45A | AL078639.1 | DPYSL3 | APLNR |
| ADGRG1 | PLPPR1 | PLCG2 | HEPN1 | IGFBP7 | NSF | HSPA1B | RNASET2 | GSN | ACTN1 | GDF15 | ATP1A2 | ENC1 | ATF3 |
| AGT | PTPRZ1 | PLEKHA4 | MMACHC | PECAM1 | THY1 | MT2A | S100A9 | QDPR | GJA1 | IGFBP2 | ATP1B2 | EPHA4 | AZGP1 |
| FAM181B | SMOC1 | RSF1 | PPP1CB | PLVAP | ATP1B1 | PLOD2 | HLA-DPB1 | SLAIN1 | IGFBP7 | IGFBP3 | CD44 | EPHA5 | B2M |
| HOPX | AQP4 | SPECC1 | SUMF2 | AGRN | SYP | BNIP3L | CTSS | KLK6 | LRIG1 | LMO2 | CD74 | GNAO1 | C1QA |
| MBP | CADM2 | TCAF1 | TP53 | A2M | TSPAN7 | CA9 | SLCO2B1 | SCD | MLC1 | LZTS1 | CPE | JPT1 | CCDC127 |
| NKAIN4 | CNTN1 | APC | AC011603.2 | SERPINH1 | TUBB2A | CXCL8 | CD53 | ABCA2 | AQP4 | MT3 | GFAP | KIF21B | CD14 |
| RGMA | CSPG5 | ARHGAP5 | ARPC4 | CD93 | PRKAR1B | DNAJB1 | CTSC | PLA2G16 | EMP1 | NAMPT | IFITM2 | MARCH4 | CLPTM1L |
| CSPG5 | ETV1 | BRD4 | BEST1 | ITGB1 | TMEM59L | BHLHE40 | HLA-DMB | APOD | FOS | PLA2G5 | LGALS3 | MEF2C | EGR2 |
| ENHO | FXYD6 | EEA1 | CNTD1 | MYH9 | CALM3 | PLIN2 | CHI3L1 | PLLP | KCNN3 | RAMP1 | MBP | MLLT11 | EGR3 |
| APOE | HMGB2 | GOLGB1 | CTBP1 | MCAM | NPTXR | CHPF | HLA-DQB1 | AMER2 | MAPK4 | S100A16 | MT1F | MPPED1 | EXOC3-AS1 |
| CKB | ID3 | KTN1 | FGD5-AS1 | APOLD1 | TMEM130 | HMOX1 | S100A11 | ELOVL1 | NCAN | SLC6A11 | MT1G | OLFM1 | FOSB |
| CRISPLD1 | MAG | MT-ND4L | MARCKS | HIGD1B | MEG3 | HSPA1A | SLC2A5 | NKX6-2 | NNMT | SLN | NTRK2 | PCP4 | G0S2 |
| EDNRB | PTN | NCOR1 | MSH6 | MIR4435-2HG | NELL2 | ARRDC3 | A2M | PLEKHH1 | NTRK2 | TMEM158 | PLPP3 | PLAT | GADD45B |
| GATM | SCD5 | NFIA | PAQR3 | SPARC | HPCA | HSPA5 | EFEMP1 | OPALIN | PDPN | TTYH3 | S100A6 | RPRM | GPX3 |
| ITM2C | TNR | PCDH9 | PDGFA | EPAS1 | IDS | SPP1 | GPNMB | SLC44A1 | S100A6 | CA12 | SEC62 | SCN3B | HLA-B |
| OLIG2 | TNS3 | SLPI | RAB6A | LAMA4 | AK5 | UPP1 | HLA-DQA1 | SPP1 | SCARA3 | CCL2 | SLC1A3 | Sep-03 | IGFBP3 |
| PTN | TSPAN7 | SOX4 | RPL23 | IFI27 | ENO2 | ZNF395 | LILRB4 | BCAS1 | SOCS3 | CDH2 | SPARC | STMN1 | IGKC |
| SCD5 | UGT8 | TCF4 | SPTBN1 | CD248 | NCDN | ARL4C | STAB1 | CSRP1 | TRIM47 | CITED1 | AL031056.1 | STMN4 | KLF6 |
| SLC1A3 | AC008080.4 | VCAN | ZBTB18 | CYTOR | SNCB | GLUL | VAMP8 | HHIP | VIM | COL9A3 | APLNR | STX1A | MAN1C1 |
| SOX8 | ANP32B | ZNF91 | AC092069.1 | FSTL1 | CRYM | HK2 | CCL2 | LINC00844 | WLS | CSRP2 | APOD | TMEM35A | MCL1 |
| ABAT | APCDD1 | ADD3 | AMFR | MMP9 | KLC1 | MIF | FTL | PLEKHB1 | AEBP1 | DBI | APOE | TUBA1A | MIDN |
| CERS1 | APOD | AL078639.1 | CPEB4 | PLXDC1 | CA11 | CEBPD | HLA-DMA | TUBB4A | APLNR | DPY19L1 | AQP1 | TUBB | OTUD1 |
| DBI | ATP6V0E2 | ANP32B | DALRD3 | THY1 | CREG2 | GBE1 | IBSP | CRYAB | BAG3 | EGR1 | C1orf61 | TUBB2A | PLAUR |
| EGFR | BEX2 | ARID4B | EIF4E | COL6A2 | GABRA1 | DDIT3 | NPC2 | LARP6 | CD99 | ELMOD1 | CAPS | TUBB2B | PSME2 |
| FABP7 | CA10 | ASH1L | GATM | MYO1B | NPTX2 | DNAJB9 | PLTP | TPPP | CXCL14 | EMP1 | CCK | TUBB3 | ROPN1L |
| FXYD6 | COL11A1 | BBX | GJA1 | SLC38A5 | BEX2 | LGALS1 | SCIN | CDKN1C | ID1 | FAM20C | CCL4 | INA | SERPINE1 |
| MARCKSL1 | COL20A1 | C5orf63 | IGDCC4 | SLC7A5 | DKK 3.00 | MT1G | CAPG | CMTM5 | PBXIP1 | FCGBP | CCNI | NELL2 | SEZ6L |
| NES | FCGBP | CAMK2N1 | MAT2A | TPM2 | FBXL16 | MT3 | CTSB | HAPLN2 | RAMP1 | FLNC | CHI3L1 | NEUROD2 | SLPI |
| NTRK2 | H3F3A | CXADR | RSRC1 | ARHGDIB | NSG2 | SLC16A1 | ZFP36 | MTURN | SPOCD1 | GATM | CNN3 | NSG2 | SOCS3 |
| OGFRL1 | ITM2A | DCX | SLITRK2 | COL6A3 | SLC1A2 | TGFBI | F13A1 | NDRG1 | ZFP36L2 | GRIA1 | COL1A2 | SYT4 | SRGN |
| PAQR8 | MDFI | ENAH | TJP1 | ESM1 | SNCG | VIM | FCGR2A | OLIG1 | A2M | HIF1A | CST3 | ZBTB18 | STAT1 |
| PHGDH | PDGFRA | HIPK2 | TOB1 | IGFBP3 | SNRPN | APLN | IFITM3 | PRR18 | ATP1A2 | HMGA1 | DKK 3.00 | CDK5R1 | TAP1 |
| RAMP1 | PFN2 | MTRNR2L1 | WEE1 | LAMC3 | TSPYL1 | CHI3L1 | OLR1 | SPOCK3 | CNN3 | ITPKC | ECEL1 | DCX | VGF |
| SPARCL1 | PRDX1 | NFE2L2 | ACTB | TIMP3 | CHGB | EIF4EBP1 | RNASE1 | CDK18 | CPE | JAG1 | EMP1 | NSG1 | WARS |
| THRA | RCC2 | NIPBL | CCND2 | CCDC3 | MLLT11 | PDPN | SOD2 | FGFR2 | CYR61 | KCNIP1 | EMP3 | RUNX1T1 | ZFP36L1 |
| AIF1L | RPLP0 | PGM2L1 | CDH2 | LUM | MOAP1 | SEC61G | APOE | KCNMB4 | ETNPPL | MATN2 | FABP5 | STMN2 | A2M |
| APC2 | SAPCD2 | PNRC1 | CNN3 | MFSD2A | NPTN | SLC39A14 | ARHGDIB | SLC48A1 | F13A1 | NCAN | FABP7 | BASP1 | ADAMTS1 |
| BAALC | SOX4 | RAB13 | CPEB2 | PTP4A3 | SCG5 | TNC | C1R | SUN2 | FAM20C | NMB | FAM181B | BHLHE22 | AIF1 |
| CDO1 | TF | RBM25 | DNAH11 | SLC2A1 | MT-ND2 | ZFAS1 | CCL3 | TTYH2 | HSPB8 | NT5E | FGF1 | CAMK2B | APOC1 |
| CLU | TMSB15A | RHOBTB3 | DNAJB6 | TIMP1 | NNAT | FN1 | CTSD | AIF1L | RDH10 | OCIAD2 | GPRC5B | CXADR | APOLD1 |
| CRYAB | TNK2 | SET | GPR37L1 | ACE | SH3GL2 | SLC6A6 | FOS | C1QB | S100A9 | PIPOX | HLA-DPA1 | LY6H | BEX3 |
| DLL3 | TOP2A | SOX11 | HNRNPK | ECSCR | STMN1 | SMIM3 | FPR1 | CA2 | SDC4 | PLAT | ID4 | NREP | C1QB |
| ETV1 | TUBA1A | THOC2 | NADK2 | FLT1 | TUBA4A | SNHG12 | GPR34 | CERCAM | TAGLN | PRDX6 | IFI6 | RAB3A | C3AR1 |
| FGFR3 | WASF1 | TNRC6B | NFIA | GGT5 | FAIM2 | SOD2 | HMOX1 | CHI3L1 | TPST1 | PTPRZ1 | METRN | REEP1 | CA3 |
| GPR17 | APOC1 | VGF | Sep-07 | CALD1 | NDRG4 | AK4 | LGALS9 | CHI3L2 | AGT | SDC3 | MT-ATP8 | SNAP25 | CD44 |
| GPRC5B | C1orf61 | ZEB2 | SEZ6L | MYL12A | PI4KA | EPAS1 | LY86 | FA2H | AHCYL1 | TIMP1 | MT-ND1 | SYT1 | CD47 |
| KCNJ10 | C1QL1 | ZMAT3 | SOX11 | PRSS23 | PTPRN | FTL | MAFB | FAM107A | ANXA2 | TNC | MT-ND2 | TUBB4A | CD74 |
| NAT8L | CMTM5 | ATRX | UGCG | APLN | RAB3A | OLFM1 | MGST1 | FTH1 | CD163 | ZYX | MT1H | UCHL1 | CNTN2 |
| PTPRZ1 | CTHRC1 | BPTF | YTHDF1 | COL5A2 | SYT4 | PI3 | MS4A7 | FTL | CTSH | AC007952.4 | MT1X | ATP1A3 | DNAJA1 |
| SDC3 | DAPL1 | ZBTB20 | ZFP36L1 | HBA2 | TAC1 | TIMP1 | PLA2G2A | GJB1 | DNER | ACSS3 | MT2A | BCL11A | ETF1 |
| TNR | HMGN2 | ANKRD12 | AC005944.1 | IFITM2 | CAMK2B | TMEM158 | RGS10 | GPR37 | FABP5 | ADAM9 | NUPR1 | CAMKV | F3 |
| TSPAN7 | IGFBP2 | CCDC88A | AC106707.1 | PODXL | FXYD6 | TPI1 | S100A4 | GPRC5B | FABP7 | ADM | PABPC1 | CD24 | FAM96A |
| TUBA1A | MARCKS | EIF5B | ACLY | SPON2 | MAP2K1 | GADD45B | SAA1 | MARCKSL1 | GFAP | AKAP12 | PLP1 | CDKN2D | FIS1 |
| ATP6V0E2 | MEST | HBA2 | ADAM10 | ABCG2 | MBP | NOL3 | SAA2 | RHOU | GPC1 | ANXA2 | PON2 | CRMP1 | FTL |
| CNTN1 | MOG | MALAT1 | AL358781.1 | ADAMTS1 | MT-CO3 | P4HA1 | SLPI | S100A10 | IFITM3 | APLP1 | RARRES3 | DPYSL3 | GABARAPL2 |
| CSRP1 | MT-CO3 | MAN1A2 | C5orf15 | APOD | RGS4 | PFKP | TMEM176B | S100B | ITGB4 | APOC1 | SELENOP | ENC1 | HAMP |
| DCLK2 | NCAN | NFIX | CDK2AP1 | CSPG4 | YWHAG | PLOD1 | AGT | SIRT2 | MATN2 | ARC | SLC39A12 | EPHA4 | HBA1 |
| FEZ1 | NELL2 | DST | DCK | FBLN1 | CELF4 | RDH10 | B2M | AATK | PLP2 | ATP1B2 | WLS | EPHA5 | HBEGF |
| FGF1 | NKAIN4 | HBA1 | ETV1 | IFITM3 | COX7A1 | SLPI | BST2 | AL078639.1 | PRSS23 | B4GALT5 | ZCRB1 | GNAO1 | HLA-A |
| HSPB8 | NT5DC2 | HBB | EZR | ITIH5 | KCNK1 | TREM1 | C1S | APOC1 | RASD1 | C1R | ZFP36 | JPT1 | HLA-C |
| MOXD1 | PHLDA1 | MT-ATP8 | FAM198B | LAMB1 | MDH1 | FTH1 | CCL4 | ATP1B1 | RGS1 | CALB1 | ZFP36L1 | KIF21B | HLA-DQA1 |
| MT-CO1 | PMP2 | NEAT1 | FAM199X | LAMC1 | MT-ATP8 | GPI | CYBA | B2M | SPARCL1 | CAPN5 | MGST1 | Mar-04 | HLA-DRA |
| MT-CYB | POLR2F | NFIC | GIGYF1 | PCOLCE | MT-ND1 | HSPA6 | ID3 | BIN1 | BAALC | CD63 | S100A13 | MEF2C | HMGN3 |
| MT-ND1 | PTTG1 | TTC3 | GPRC5B | RGCC | NEFM | MXRA7 | ISG15 | BOK | DDR1 | CD68 | FXYD1 | MLLT11 | HMOX1 |
| MT-ND2 | RPL15 | ZC3H13 | HERPUD2 | ROBO4 | PLP1 | NAMPT | LST1 | CERS2 | DKK 3.00 | CDKN1A | MT1M | MPPED1 | HSPA8 |
| MT-ND3 | RPS2 | AKAP9 | HIF1A | S1PR3 | PPP3CA | NDUFA4L2 | LYZ | EFHD1 | EGR1 | CDKN2A | MT-ND4L | OLFM1 | HSPB1 |
| MT3 | SLC25A5 | CCNI | HNRNPA0 | COL6A1 | PRKCB | SLC6A8 | MGP | LAMP2 | GBP1 | CHL1 | MT1E | PCP4 | HSPB8 |
| PHYHIPL | SNAP25 | GOLGA4 | LTBP1 | COX7A1 | SST | SLN | MS4A4A | LANCL1 | GLRX | CST3 | S100A1 | PLAT | HSPE1 |
| PLEKHB1 | SNHG25 | HIST1H4C | MMP19 | CYR61 | B4GAT1 | VKORC1 | MT-ATP6 | LTF | GPX3 | CYR61 | SLC1A2 | RPRM | IER3 |
| PMP2 | SPP1 | KMT2E | NUDT4 | JAG1 | CNTN1 | ALDOA | PABPC1 | MTRNR2L12 | ID4 | DIRAS3 | ADIRF | SCN3B | IFI27 |

### Subcluster metaprograms full

| Vasc1 (Vasc-Ang) | Vasc2 (Vasc-IMEC) | Mac1 | Mac2 | Mac3 | MES_1 | MES_2 | MES_3 | MES.Ast_1 | MES.Ast_2 |
|---|---|---|---|---|---|---|---|---|---|
| COL4A1 | FN1 | FCER1G | CD74 | S100A8 | AQP1 | C1R | ELOVL2 | GPC1 | FABP5 |
| COL4A2 | VWF | C1QC | APOC1 | S100A9 | MGST1 | CHI3L1 | F3 | METTL7B | SERPINE1 |
| VWF | COL4A1 | FCGR3A | C1QA | C1QA | EFEMP1 | EFEMP1 | IGFBP2 | LPL | CCL2 |
| PDGFRB | CLDN5 | LAPTM5 | HLA-DPA1 | C1QB | ITM2C | C1S | KLHDC8A | MOXD1 | CXCL14 |
| COL1A2 | DCN | CD14 | C1QB | HAMP | GJA1 | CD44 | LPL | PDLIM4 | DNAJB1 |
| ENG | EPAS1 | SRGN | C1QC | CHI3L1 | MAOB | LTF | METTL7B | RCAN1 | EMP1 |
| RGS5 | ITM2A | AIF1 | HLA-DRA | FCER1G | ATP1B2 | SOD2 | MOXD1 | AGT | FLNC |
| COL18A1 | PDGFRB | C1QB | TYROBP | SPP1 | CD44 | CCL2 | POSTN | CXCL14 | GADD45A |
| COL3A1 | A2M | CSF1R | HLA-DRB1 | AIF1 | CHI3L1 | RCAN1 | SLC6A11 | FABP7 | GAP43 |
| HSPG2 | COL4A2 | VSIG4 | LAPTM5 | ALOX5AP | CHI3L2 | TNC | TTYH3 | GDF15 | HMGA1 |
| BGN | MGP | TREM2 | CD14 | C3 | KCNN3 | ANXA1 | CHPF | HOPX | JAG1 |
| FN1 | MYL9 | ALOX5AP | FCER1G | CD14 | LRIG1 | MAN1C1 | FJX1 | LGALS3 | METTL7B |
| NDUFA4L2 | SLC2A1 | C1QA | HLA-DPB1 | CD163 | NCAN | NAMPT | GPC1 | LZTS1 | NAMPT |
| ESAM | TAGLN | ITGB2 | C3 | CD74 | NTRK2 | ZFP36 | GRIA1 | MT2A | S100A16 |
| ITGA1 | TIMP3 | HLA-DRB5 | SRGN | F13A1 | ANXA1 | CHI3L2 | HOPX | POSTN | SCG2 |
| ITM2A | APOLD1 | CD163 | A2M | FCGBP | BAG3 | GAP43 | KCNIP1 | PTN | TNFRSF12A |
| ANGPT2 | BGN | CD53 | CD68 | FOS | ID3 | MGST1 | NCAN | FABP5 | ADM |
| CLDN5 | COL18A1 | CTSS | CSF1R | IBSP | LTF | NNMT | NT5E | GAP43 | AKAP12 |
| IGFBP4 | COL1A2 | HAMP | FTL | S100A11 | MLC1 | SOCS3 | PIPOX | IGFBP3 | ANXA2 |
| MGP | EGFL7 | HLA-DRB1 | AIF1 | APOC1 | PBXIP1 | C3 | PTN | KCNF1 | CDKN1A |
| MYL9 | ENG | MS4A6A | APOE | B2M | AL049839.2 | CP | AC007952.4 | NMB | F3 |
| NOTCH3 | GGT5 | RNASET2 | FCGR3A | C1R | AQP4 | FCGBP | ACSS3 | PDPN | GPC1 |
| CTGF | GNG11 | CTSC | GPNMB | C1S | ATP1A2 | FOS | ATP1B2 | RAMP1 | HIF1A |
| DCN | IFI27 | FCGBP | ITGB2 | CCL2 | CLU | ID3 | C1R | SLN | ITPKC |
| EGFL7 | MCAM | TYROBP | RGS1 | CCL3 | ETNPPL | PDPN | CA12 | SPOCD1 | LGALS3 |
| NID1 | RGS5 | S100A8 | RNASE1 | CTSB | F3 | SERPING1 | CHI3L1 | TNFRSF12A | LMNA |
| COL1A1 | ACTA2 | C3 | S100A11 | EFEMP1 | MAPK4 | SPOCD1 | COL9A3 | ADAM9 | MAP2K3 |
| GNG11 | CD248 | CD68 | SPP1 | HLA-DRB1 | S100A10 | AEBP1 | CST3 | APLP1 | MIDN |
| PLVAP | COL1A1 | SLCO2B1 | TREM2 | ID3 | SDC4 | CYR61 | CXCL14 | APOC1 | MT2A |
| PECAM1 | COL3A1 | SPP1 | APOD | LAPTM5 | WLS | EMP1 | FABP7 | CALB1 | OLR1 |
| ACTA2 | COL6A2 | CD74 | FCGBP | LTF | AGT | F13A1 | FCGBP | CAPN5 | PDLIM4 |
| AGRN | ESAM | HLA-DMB | HLA-DMB | PLTP | CP | F3 | ITGA3 | CD68 | PDPN |
| IGFBP7 | HIGD1B | HLA-DQB1 | MBP | RCAN1 | CPE | FAM20C | KCNF1 | CDH2 | PFKFB3 |
| SERPINH1 | IGFBP4 | HLA-DRA | MT-ATP6 | RGS1 | CTSH | ID1 | KLHL4 | CDKN2A | RCAN1 |
| TAGLN | IGFBP7 | RGS1 | MT-ND2 | SAT1 | CXCL14 | S100A10 | LANCL2 | CHL1 | SOD2 |
| CD93 | MMP9 | SLC2A5 | MT-ND4L | SOD2 | DNER | S100A6 | LGR6 | CITED1 | SPOCD1 |
| ITGB1 | SLC7A5 | HLA-DMA | MTRNR2L12 | STAB1 | HSPB8 | S100A9 | LYPD1 | CSRP2 | SPRY2 |
| A2M | THY1 | HLA-DPB1 | PABPC1 | TXNIP | IGFBP7 | TPST1 | MATN2 | DBI | TMEM158 |
| MYH9 | ABCG2 | LILRB4 | RNASET2 | TYROBP | SOD2 | TRIM47 | METRN | DIRAS3 | TNC |
| SPARC | APOD | S100A9 | SLCO2B1 | ZFP36 | TNC | ZFP36L2 | MRC2 | DLK1 | ZYX |
| MIR4435-2HG | CTGF | VAMP8 | TAGLN | A2M | ACTN1 | A2M | PEG10 | DPY19L1 | FABP5 |
| LAMA4 | CYTOR | CHI3L1 | CAPG | AGT | APLNR | ACTN1 | PLA2G5 | ELMOD1 | SERPINE1 |
| HIGD1B | FBLN1 | FCGR2A | CLDN5 | ANXA2 | C1R | AL049839.2 | PLAT | FAM20C | CCL2 |
| MCAM | FXYD5 | HLA-DPA1 | CTSD | AQP1 | C3 | ANXA2 | PTGFRN | FJX1 | CXCL14 |
| APOLD1 | HBA2 | HLA-DQA1 | DCN | BST2 | DKK 3.00 | CD163 | PTPRZ1 | GATM | DNAJB1 |
| FSTL1 | HLA-E | STAB1 | FAM107A | BTG2 | ID4 | CD99 | RCAN1 | GRIK3 | EMP1 |
| PLXDC1 | IFITM2 | EFEMP1 | HBA2 | C1QC | MATN2 | CLU | SLC4A4 | ITGB8 | FLNC |
| CYTOR | IFITM3 | FPR1 | HLA-DQA1 | CCL4 | PLPP3 | PRSS23 | SPON1 | KLHDC8A | GADD45A |
| IFI27 | IGFBP3 | LGALS9 | HLA-DQB1 | CCL4L2 | SCARA3 | RGS1 | SPRY4 | LMO2 | GAP43 |
| CD248 | KLF2 | LY86 | HLA-E | CCT6A | SERPING1 | TAGLN | TNFRSF12A | LSM 7.00 | HMGA1 |
| EPAS1 | LYZ | OLR1 | HSPA1A | CD44 | SLC14A1 | AQP1 | WSCD1 | MAN1C1 | JAG1 |
| MYO1B | MFSD2A | SCIN | MALAT1 | CD68 | SLC1A2 | FABP5 | ELOVL2 | MT3 | METTL7B |
| TPM2 | MYH9 | APOC1 | MGP | CHI3L2 | SLC1A3 | GBP1 | F3 | NRCAM | NAMPT |
| COL6A3 | PECAM1 | CAPG | MT-ATP8 | COL3A1 | SPARCL1 | GFAP | IGFBP2 | PLA2G5 | S100A16 |
| MMP9 | PODXL | CCL2 | MT-ND3 | CP | VIM | GLRX | KLHDC8A | PLAU | SCG2 |
| THY1 | PRSS23 | NPC2 | MT-ND4 | CTSL | C1S | GPC1 | LPL | PMEPA1 | TNFRSF12A |
| ARHGDIB | PTP4A3 | PLTP | MTRNR2L8 | CXCL10 | C2orf40 | IGFBP7 | METTL7B | PRDX6 | ADM |
| COL6A2 | S1PR3 | RGS10 | PLP1 | DUSP1 | CNN3 | MAOB | MOXD1 | RASD1 | AKAP12 |
| ESM1 | SERPINE1 | SERPINA1 | RPL39 | FABP5 | DCLK1 | PLA2G2A | POSTN | RBP1 | ANXA2 |
| LAMC3 | SLC38A5 | SLC11A1 | TPT1 | FCGR3A | EZR | PLP2 | SLC6A11 | SCG2 | CDKN1A |
| SLC38A5 | SLC39A10 | SLPI | ACTA2 | HLA-C | FBXO2 | PTX3 | TTYH3 | SDC3 | F3 |
| ACE | SRGN | ZFP36 | ACTG2 | HLA-DRA | FCGBP | RAMP1 | CHPF | SERPINE1 | GPC1 |
| CCDC3 | TGM2 | AL049839.2 | AL078639.1 | HMOX1 | GAP43 | RASD1 | FJX1 | SPOCK2 | HIF1A |
| ECSCR | TPM1 | CORO1A | AL355075.4 | IFITM2 | GPX3 | RGS2 | GPC1 | SULF2 | ITPKC |
| FLT1 | ADAMTS1 | CREG1 | AP2S1 | IFITM3 | IFITM3 | S100A8 | GRIA1 | TMEM158 | LGALS3 |
| LUM | AGRN | FCGR1A | ARHGDIB | IGFBP2 | ITGB4 | SAA1 | HOPX | TRIM47 | LMNA |
| SLC7A5 | ANGPT2 | FOLR2 | ARPC1B | ISG15 | MAN1C1 | SCARA3 | KCNIP1 | TRIR | MAP2K3 |
| TIMP1 | APCDD1 | GPNMB | BCAN | MGP | NAMPT | TCIM | NCAN | TSC22D1 | MIDN |
| IGFBP3 | APLN | GPR34 | BCAS1 | MGST1 | PFKFB3 | TGFBI | NT5E | TSC22D4 | MT2A |
| MYL12A | APOC1 | HP | C5orf63 | MS4A6A | PLEC | TNFRSF12A | PIPOX | UBE2S | OLR1 |
| COL5A2 | ARHGEF17 | IBSP | CCK | MS4A7 | PTPRZ1 | VIM | PTN | GPC1 | PDLIM4 |
| MFSD2A | ATP1B3 | LAIR1 | CCL3 | NAMPT | RAMP1 | ABCC3 | AC007952.4 | METTL7B | PDPN |
| PTP4A3 | C1QA | MAFB | CCL4 | NPC2 | RDH10 | AHCYL1 | ACSS3 | LPL | PFKFB3 |
| SPON2 | C1QC | MS4A4A | CD163 | PTN | SCD5 | APLNR | ATP1B2 | MOXD1 | RCAN1 |
| CALD1 | CALD1 | MSR1 | COTL1 | RBP1 | SERPINE2 | ATP1B2 | C1R | PDLIM4 | SOD2 |
| CSPG4 | CD93 | PLA2G2A | COX7A2 | S100A4 | SOX2 | BAALC | CA12 | RCAN1 | SPOCD1 |
| ITIH5 | CYP1B1 | SAA1 | CST3 | SERPINE1 | SOX9 | BCL6 | CHI3L1 | AGT | SPRY2 |
| LAMC1 | DUSP1 | SAA2 | CTSB | SOCS3 | TMEM132A | C1QL1 | COL9A3 | CXCL14 | TMEM158 |
| PCOLCE | EFNB1 | SIGLEC10 | CTSH | SRGN | TRIL | CNN3 | CST3 | FABP7 | TNC |
| RGCC | FSTL1 | TMEM176B | CYBA | TMEM176B | AC007952.4 | COL8A1 | CXCL14 | GDF15 | ZYX |
| ROBO4 | GPCPD1 | TMIGD3 | DCX | TSC22D1 | ADIRF | EGR1 | FABP7 | HOPX | FABP5 |
| APLN | HBA1 | ADORA3 | ENC1 | VIM | AHCYL1 | FABP7 | FCGBP | LGALS3 | SERPINE1 |
| LAMB1 | HBB | ARHGDIB | FTH1 | VSIG4 | ALDH1L1 | FLNA | ITGA3 | LZTS1 | CCL2 |
| PRSS23 | HLA-DPA1 | C1R | FXYD5 | S100A8 | ANOS1 | GLIPR1 | KCNF1 | MT2A | CXCL14 |
| SPINK8 | HLA-DQB1 | CTSB | GFAP | S100A9 | APOC1 | HAMP | KLHL4 | POSTN | DNAJB1 |
| TIMP3 | HLA-DRB1 | CYBA | GPM6A | C1QA | APOE | IL32 | LANCL2 | PTN | EMP1 |
| ADAMTS1 | HSPA1A | F13A1 | GPR34 | C1QB | B3GAT2 | JUNB | LGR6 | FABP5 | FLNC |
| COL6A1 | HYAL2 | GPX3 | GSN | HAMP | C1orf61 | MMP7 | LYPD1 | GAP43 | GADD45A |
| COX7A1 | ID1 | HCLS1 | HAMP | CHI3L1 | CA3 | OCIAD2 | MATN2 | IGFBP3 | GAP43 |
| CYR61 | IFNGR1 | HMOX1 | HBA1 | FCER1G | CCL2 | PLAU | METRN | KCNF1 | HMGA1 |
| GGT5 | ITGA1 | ID1 | HBB | SPP1 | CD99 | PLTP | MRC2 | NMB | JAG1 |
| HBA2 | ITGB1 | IFITM3 | HLA-B | AIF1 | CNTN1 | RARRES3 | PEG10 | PDPN | METTL7B |
| IFITM2 | JAG1 | LST1 | HLA-DRB5 | ALOX5AP | CRISPLD1 | RDH10 | PLA2G5 | RAMP1 | NAMPT |
| PLXND1 | LAMA4 | LYZ | HSPA1B | C3 | CST3 | SPARC | PLAT | SLN | S100A16 |
| PODXL | MIR4435-2HG | MGST1 | IFITM2 | CD14 | DDR1 | TAGLN2 | PTGFRN | SPOCD1 | SCG2 |
| SLC2A1 | MYO1B | MS4A7 | IFITM3 | CD163 | DEPP1 | TIPARP | PTPRZ1 | TNFRSF12A | TNFRSF12A |
| UACA | NAMPT | PLAU | IGHA1 | CD74 | EDNRB | ACTB | RCAN1 | ADAM9 | ADM |
| ABCG2 | NDUFA4L2 | PYCARD | IGHG1 | F13A1 | EMP1 | AHR | SLC4A4 | APLP1 | AKAP12 |
| ADGRF5 | NID1 | SOD2 | IGHG3 | FCGBP | EPHX1 | AQP4 | SPON1 | APOC1 | ANXA2 |
| APOD | NOTCH3 | TOP2A | IGKC | FOS | FABP7 | BTG2 | SPRY4 | CALB1 | CDKN1A |

#### Subcluster metaprograms unique

| Vasc-Ang_unique | Vasc-IMEC_MP2_unique | mac_MP1_uni | mac_MP2_uni | mac_MP3_uni | mes_MP1_uni | mes_MP2_uni | MES.Ast_MP1 | MES.Ast_MP2 | MES.Ast_MP3_unique |
|---|---|---|---|---|---|---|---|---|---|
| HSPG2 | FBLN1 | CD53 | FTL | FOS | ITM2C | RCAN1 | ELOVL2 | AGT | CCL2 |
| PLVAP | FXYD5 | CTSS | APOE | B2M | GJA1 | ZFP36 | IGFBP2 | GDF15 | DNAJB1 |
| SERPINH1 | HLA-E | CTSC | RNASE1 | C1S | KCNN3 | NNMT | SLC6A11 | LZTS1 | EMP1 |
| SPARC | IFITM3 | SLC2A5 | APOD | ID3 | LRIG1 | SOCS3 | TTYH3 | IGFBP3 | FLNC |
| PLXDC1 | KLF2 | HLA-DMA | MBP | LTF | NCAN | FOS | CHPF | NMB | GADD45A |
| TPM2 | LYZ | LILRB4 | MT-ATP6 | RCAN1 | NTRK2 | PDPN | GRIA1 | RAMP1 | HMGA1 |
| COL6A3 | S1PR3 | VAMP8 | MT-ND2 | SAT1 | BAG3 | SPOCD1 | KCNIP1 | SLN | JAG1 |
| ARHGDIB | SERPINE1 | FCGR2A | MT-ND4L | TXNIP | MLC1 | AEBP1 | NCAN | ADAM9 | NAMPT |
| ESM1 | SLC39A10 | FPR1 | MTRNR2L12 | AGT | PBXIP1 | CYR61 | NT5E | APLP1 | S100A16 |
| LAMC3 | SRGN | LGALS9 | PABPC1 | ANXA2 | ATP1A2 | F13A1 | PIPOX | APOC1 | ADM |
| ACE | TGM2 | LY86 | TAGLN | AQP1 | ETNPPL | FAM20C | AC007952.4 | CALB1 | AKAP12 |
| CCDC3 | TPM1 | OLR1 | CLDN5 | BST2 | MAPK4 | ID1 | ACSS3 | CAPN5 | ANXA2 |
| ECSCR | APCDD1 | SCIN | CTSD | BTG2 | SDC4 | S100A6 | ATP1B2 | CD68 | CDKN1A |
| FLT1 | APOC1 | RGS10 | DCN | CCL4L2 | WLS | S100A9 | C1R | CDH2 | HIF1A |
| LUM | ARHGEF17 | SERPINA1 | FAM107A | CCT6A | AGT | TPST1 | CA12 | CDKN2A | ITPKC |
| TIMP1 | ATP1B3 | SLC11A1 | HBA2 | CD44 | CPE | TRIM47 | CHI3L1 | CHL1 | LMNA |
| MYL12A | C1QA | SLPI | HLA-E | CHI3L2 | CTSH | ZFP36L2 | COL9A3 | CITED1 | MAP2K3 |
| COL5A2 | C1QC | AL049839.2 | HSPA1A | COL3A1 | CXCL14 | A2M | CST3 | CSRP2 | MIDN |
| SPON2 | CYP1B1 | CORO1A | MALAT1 | CP | DNER | ANXA2 | FCGBP | DBI | OLR1 |
| CSPG4 | DUSP1 | CREG1 | MT-ATP8 | CTSL | HSPB8 | CD163 | ITGA3 | DIRAS3 | PFKFB3 |
| ITIH5 | EFNB1 | FCGR1A | MT-ND3 | CXCL10 | DKK 3.00 | PRSS23 | KLHL4 | DLK1 | SOD2 |
| LAMC1 | GPCPD1 | FOLR2 | MT-ND4 | DUSP1 | ID4 | RGS1 | LANCL2 | DPY19L1 | SPRY2 |
| PCOLCE | HBA1 | HP | MTRNR2L8 | FABP5 | MATN2 | TAGLN | LGR6 | ELMOD1 | TNC |
| RGCC | HBB | LAIR1 | PLP1 | HLA-C | PLPP3 | FABP5 | LYPD1 | FAM20C | ZYX |
| ROBO4 | HLA-DPA1 | MAFB | RPL39 | IGFBP2 | SLC14A1 | GBP1 | MATN2 | GATM | CCL2 |
| LAMB1 | HLA-DQB1 | MS4A4A | TPT1 | ISG15 | SLC1A2 | GFAP | METRN | GRIK3 | DNAJB1 |
| SPINK8 | HLA-DRB1 | MSR1 | ACTA2 | NAMPT | SLC1A3 | GLRX | MRC2 | ITGB8 | EMP1 |
| COL6A1 | HSPA1A | PLA2G2A | ACTG2 | PTN | SPARCL1 | GPC1 | PEG10 | LMO2 | FLNC |
| COX7A1 | HYAL2 | SAA1 | AL078639.1 | RBP1 | C2orf40 | PLA2G2A | PLAT | LSM 7.00 | GADD45A |
| CYR61 | ID1 | SAA2 | AL355075.4 | S100A4 | DCLK1 | PLP2 | PTGFRN | MAN1C1 | HMGA1 |
| PLXND1 | IFNGR1 | SIGLEC10 | AP2S1 | SERPINE1 | EZR | PTX3 | PTPRZ1 | MT3 | JAG1 |
| UACA | JAG1 | TMIGD3 | ARPC1B | SOCS3 | FBXO2 | RASD1 | SLC4A4 | NRCAM | NAMPT |
| ADGRF5 | NAMPT | ADORA3 | BCAN | TSC22D1 | GPX3 | RGS2 | SPON1 | PLAU | S100A16 |
| HSPG2 | FBLN1 | GPX3 | BCAS1 | VIM | IFITM3 | S100A8 | SPRY4 | PMEPA1 | ADM |
| PLVAP | FXYD5 | HCLS1 | C5orf63 | FOS | ITGB4 | SAA1 | WSCD1 | PRDX6 | AKAP12 |
| SERPINH1 | HLA-E | ID1 | CCK | B2M | PFKFB3 | TCIM | ELOVL2 | RASD1 | ANXA2 |
| SPARC | IFITM3 | LST1 | COTL1 | C1S | PLEC | TGFBI | IGFBP2 | RBP1 | CDKN1A |
| PLXDC1 | KLF2 | LYZ | COX7A2 | ID3 | PTPRZ1 | TNFRSF12A | SLC6A11 | SDC3 | HIF1A |
| TPM2 | LYZ | PLAU | CST3 | LTF | SCD5 | ABCC3 | TTYH3 | SPOCK2 | ITPKC |
| COL6A3 | S1PR3 | PYCARD | CTSH | RCAN1 | SERPINE2 | BAALC | CHPF | SULF2 | LMNA |
| ARHGDIB | SERPINE1 | TOP2A | DCX | SAT1 | SOX2 | BCL6 | GRIA1 | TRIM47 | MAP2K3 |
| ESM1 | SLC39A10 | CD53 | ENC1 | TXNIP | SOX9 | C1QL1 | KCNIP1 | TRIR | MIDN |
| LAMC3 | SRGN | CTSS | FTH1 | AGT | TMEM132A | COL8A1 | NCAN | TSC22D1 | OLR1 |
| ACE | TGM2 | CTSC | FXYD5 | ANXA2 | TRIL | EGR1 | NT5E | TSC22D4 | PFKFB3 |
| CCDC3 | TPM1 | SLC2A5 | GFAP | AQP1 | AC007952.4 | FLNA | PIPOX | UBE2S | SOD2 |
| ECSCR | APCDD1 | HLA-DMA | GPM6A | BST2 | ADIRF | GLIPR1 | AC007952.4 | AGT | SPRY2 |
| FLT1 | APOC1 | LILRB4 | GSN | BTG2 | ALDH1L1 | HAMP | ACSS3 | GDF15 | TNC |
| LUM | ARHGEF17 | VAMP8 | HBA1 | CCL4L2 | ANOS1 | IL32 | ATP1B2 | LZTS1 | ZYX |
| TIMP1 | ATP1B3 | FCGR2A | HBB | CCT6A | APOC1 | JUNB | C1R | IGFBP3 | CCL2 |
| MYL12A | C1QA | FPR1 | HLA-B | CD44 | APOE | MMP7 | CA12 | NMB | DNAJB1 |
| COL5A2 | C1QC | LGALS9 | HSPA1B | CHI3L2 | B3GAT2 | OCIAD2 | CHI3L1 | RAMP1 | EMP1 |
| SPON2 | CYP1B1 | LY86 | IGHA1 | COL3A1 | C1orf61 | PLAU | COL9A3 | SLN | FLNC |
| CSPG4 | DUSP1 | OLR1 | IGHG1 | CP | CA3 | PLTP | CST3 | ADAM9 | GADD45A |
| ITIH5 | EFNB1 | SCIN | IGHG3 | CTSL | CNTN1 | RARRES3 | FCGBP | APLP1 | HMGA1 |
| LAMC1 | GPCPD1 | RGS10 | IGKC | CXCL10 | CRISPLD1 | SPARC | ITGA3 | APOC1 | JAG1 |
| PCOLCE | HBA1 | SERPINA1 | FTL | DUSP1 | CST3 | TAGLN2 | KLHL4 | CALB1 | NAMPT |
| RGCC | HBB | SLC11A1 | APOE | FABP5 | DDR1 | TIPARP | LANCL2 | CAPN5 | S100A16 |
| ROBO4 | HLA-DPA1 | SLPI | RNASE1 | HLA-C | DEPP1 | ACTB | LGR6 | CD68 | ADM |
| LAMB1 | HLA-DQB1 | AL049839.2 | APOD | IGFBP2 | EDNRB | AHR | LYPD1 | CDH2 | AKAP12 |
| SPINK8 | HLA-DRB1 | CORO1A | MBP | ISG15 | EPHX1 | BTG2 | MATN2 | CDKN2A | ANXA2 |

#### Table S3: Overview antibodies and cycles

| Cycle # | Target | Reporter | Dilution (1:x) | Exposure time in ms | Company | Cat number | Clone | Fluorochrome |
|---|---|---|---|---|---|---|---|---|
| Cycle 1 | CD45 | **RX001** | 200 | 150 | Akoya | 4150003 | NA | AF488 |
| | CD3 | **RX015** | 400 | 150 | Akoya | 4550103 | NA | AF647 |
| | CD69 | **RX041** | 200 | 150 | Akoya | 4250022 | NA | Atto550 |
| Cycle 2 | CD90 | **RX022** | 400 | 150 | Akoya | 4150021 | NA | AF488 |
| | CD11c | **RX027** | 200 | 150 | Akoya | 4550107 | NA | AF647 |
| | Ki67 | **RX047** | 200 | 150 | Akoya | 4250019 | NA | Atto550 |
| Cycle 3 | CD8 | **RX004** | 200 | 200 | Akoya | 4150004 | NA | AF488 |
| | CD4 | **RX021** | 200 | 200 | Akoya | 4550105 | NA | AF647 |
| | HLA-DR | **RX026** | 265 | 150 | Akoya | 4250006 | NA | Atto550 |
| Cycle 4 | FN1 | **RX034** | 1300 | 150 | Abcam | ab268022 | EPR23110 | AF488 |
| | SOX10 | **RX050** | 200 | 170 | Abcam | ab220078 | EPR4007-104 | AF647 |
| | CD31 | **RX032** | 200 | 200 | Akoya | 4250009 | NA | Atto550 |
| Cycle 5 | VIM | **RX025** | 1000 | 150 | BioLegend | 677802 | O91D3 | AF488 |
| | PLP1 | **RX006** | 600 | 150 | Abcam | ab275751-100 | EPR23504-106 | AF647 |
| | PDPN | **RX023** | 100 | 150 | Akoya | 4250004 | NA | Atto550 |
| Cycle 6 | GFAP | **RX031** | 600 | 150 | Akoya | 837404 | SMI24 | AF488 |
| | EGFR | **RX030** | 225 | 150 | Abcam | ab272293-100 | EP38Y | AF647 |
| | CD279 | **RX014** | 200 | 200 | Akoya | 4250010 | NA | Atto550 |
| Cycle 7 | SOX4 | **RX007** | 100 | 150 | Abcam | ab237903 | SOX4/2540 | AF488 |
| | CD14 | **RX024** | 100 | 175 | BioLegend | 325602 | HCD14 | AF647 |
| | DCX | **RX020** | 400 | 150 | Abcam | ab222921 | EPR19997 | Atto550 |
| Cycle 8 | AQP4 | **RX010** | 200 | 150 | Abcam | ab282586 | EPR24281-65 | AF488 |
| | OLIG2 | **RX045** | 200 | 150 | Abcam | ab220796 | EPR2673 | AF647 |
| | NeuN | **RX054** | 150 | 175 | Abcam | ab209898 | EPR12763 | Atto550 |
| Cycle 9 | PDGFRA | **RX016** | 100 | 150 | Abcam | ab234965 | EPR22059-720 | AF488 |
| | CA9 | **RX033** | 300 | 150 | Abcam | ab270401 | EPR23055-5 | AF647 |
| | GLUT1 | **RX052** | 1400 | 150 | Abcam | ab252403 | EPR3915 | Atto550 |
| Cycle 10 | BCAN | **RX040** | 800 | 150 | Abcam | ab285169 | EPR25743-59 | AF488 |
| | CD44 | **RX042** | 400 | 150 | Novus | NBP1-41266 | IM7 | AF647 |
| | p53 | **RX037** | 400 | 150 | Abcam | ab225531 | E26 | Atto550 |
| Cycle 11 | GAP43 | **RX019** | 400 | 150 | Abcam | ab219582 | EP890Y | AF488 |
| | CD163 | **RX043** | 400 | 150 | BioLegend | 333602 | GHI/61 | AF647 |
| | NDRG1 | **RX029** | 100 | 175 | Abcam | ab226082 | EPR5593 | Atto550 |
| Cycle 12 | APOD | **RX028** | 600 | 150 | Abcam | ab261911 | EPR22967-72 | AF488 |
| | SOX2 | **RX036** | 65 | 200 | Abcam | ab215970 | EPR3131 | AF647 |
| Cycle 13 | | | | | | | | |
| | S100B | **RX046** | 300 | 150 | CellSignaling | #42397 | E7C3A | AF647 |
| | APOE | **RX005** | 65 | 200 | BioLegend | 803301 | D6E10 | Atto550 |
| Cycle 14 | | | | | | | | |
| | CHI3L1 | **RX017** | 65 | 200 | Abcam | ab255864 | EPR19078-157 | Atto550 |
| Cycle 15 | | | | | | | | |
| | CD19 | **RX003** | 200 | 150 | Akoya | 4550099 | NA | AF647 |
| | MAP2 | **RX035** | 100 | 150 | Abcam | ab288577 | RM1037 | Atto550 |

Fig. S1

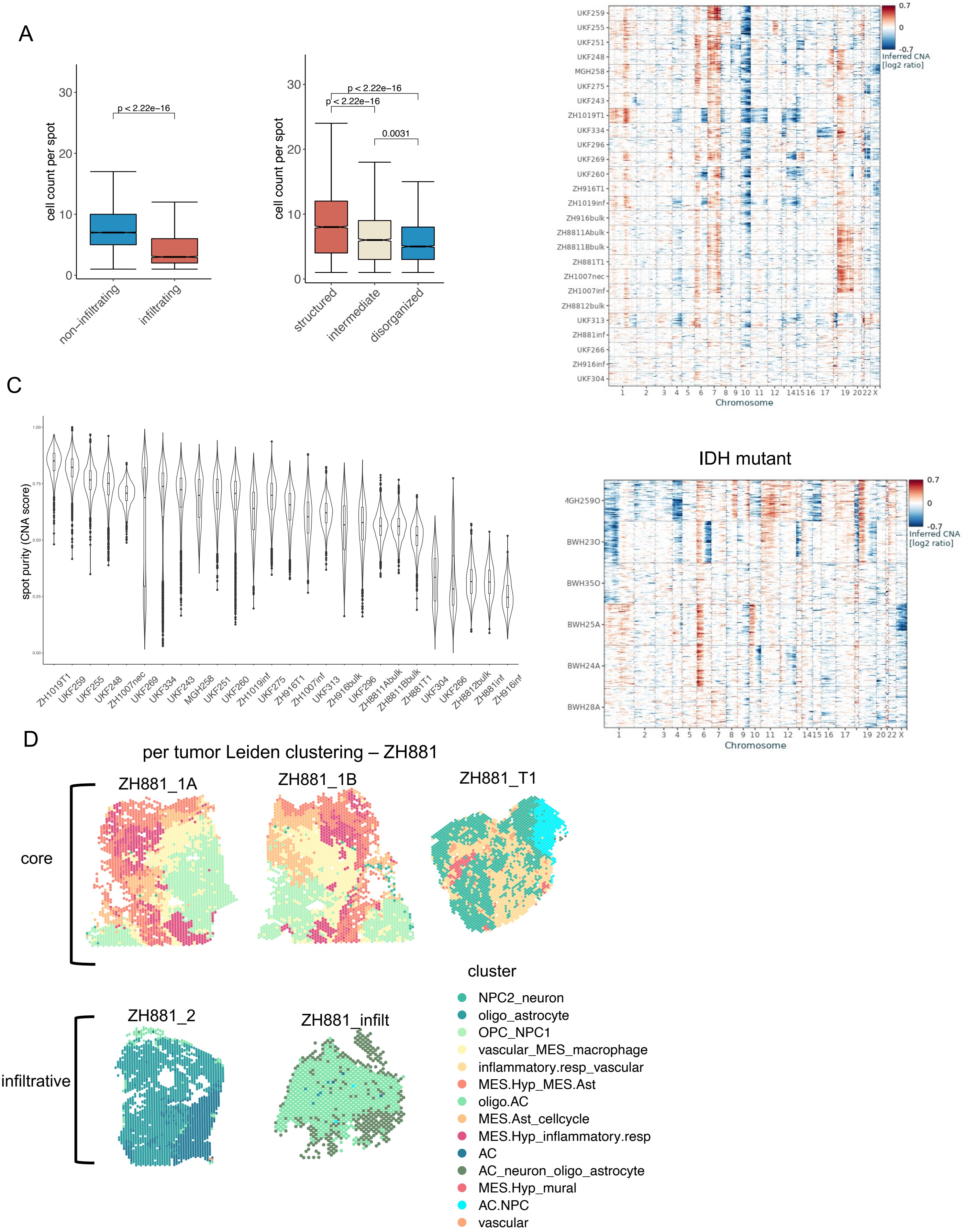
A
cell count per spot
p < 2.22e−16
non−infiltrating
infiltrating
p < 2.22e−16
p < 2.22e−16
0.0031
structured
intermediate
disorganized
B
IDH WT
Inferred CNA [log2 ratio]
UKF259
UKF255
UKF251
UKF248
MGH258
UKF275
UKF243
ZH1019T1
UKF334
UKF296
UKF269
UKF260
ZH916T1
ZH1019inf
ZH916bulk
ZH8811Abulk
ZH8811Bbulk
ZH881T1
ZH1007nec
ZH1007inf
ZH8812bulk
UKF313
ZH881inf
UKF266
ZH916inf
UKF304
Chromosome
IDH mutant
MGH259O
BWH23O
BWH35O
BWH25A
BWH24A
BWH28A
C
spot purity (CNA score)
ZH1019T1
UKF259
UKF255
UKF248
ZH1007nec
UKF269
UKF334
UKF243
MGH258
UKF251
UKF260
ZH1019inf
UKF275
ZH916T1
ZH1007inf
UKF313
ZH916bulk
UKF296
ZH8811Abulk
ZH8811Bbulk
ZH881T1
UKF304
UKF266
ZH8812bulk
ZH881inf
ZH916inf
D
per tumor Leiden clustering – ZH881
ZH881_1A
ZH881_1B
ZH881_T1
core
ZH881_2
ZH881_infilt
infiltrative
cluster
NPC2_neuron
oligo_astrocyte
OPC_NPC1
vascular_MES_macrophage
inflammatory.resp_vascular
MES.Hyp_MES.Ast
oligo.AC
MES.Ast_cellcycle
MES.Hyp_inflammatory.resp
AC
AC_neuron_oligo_astrocyte
MES.Hyp_mural
AC.NPC
vascular

**Figure S1. Segmentation and CNA inference**

**(a)** Segmentation analysis: Left panel: Cell density (cell count per 55 µm spot) for non-infiltrative and infiltrative samples. Right panel: Cell density (cell count per 55 µm spot) for structured, intermediate and disorganized regions as classified by spatial coherence analysis on CODEX pseudo-spots. **(b)** Copy number aberrations (CNAs) across all samples for GBM WT and IDH mutant cohorts. CNAs were inferred by average relative expression in windows of 150 analyzed genes. Rows correspond to spots arranged by malignancy level inference (see Methods), columns correspond to genes arranged by chromosomal position. **(c)** Spot CNA score by sample for GBM samples (Methods). **(d)** In tumors for which there were multiple samples, Leiden clustering was performed per tumor jointly across the tissue sections (in addition to per sample).

Fig. S2

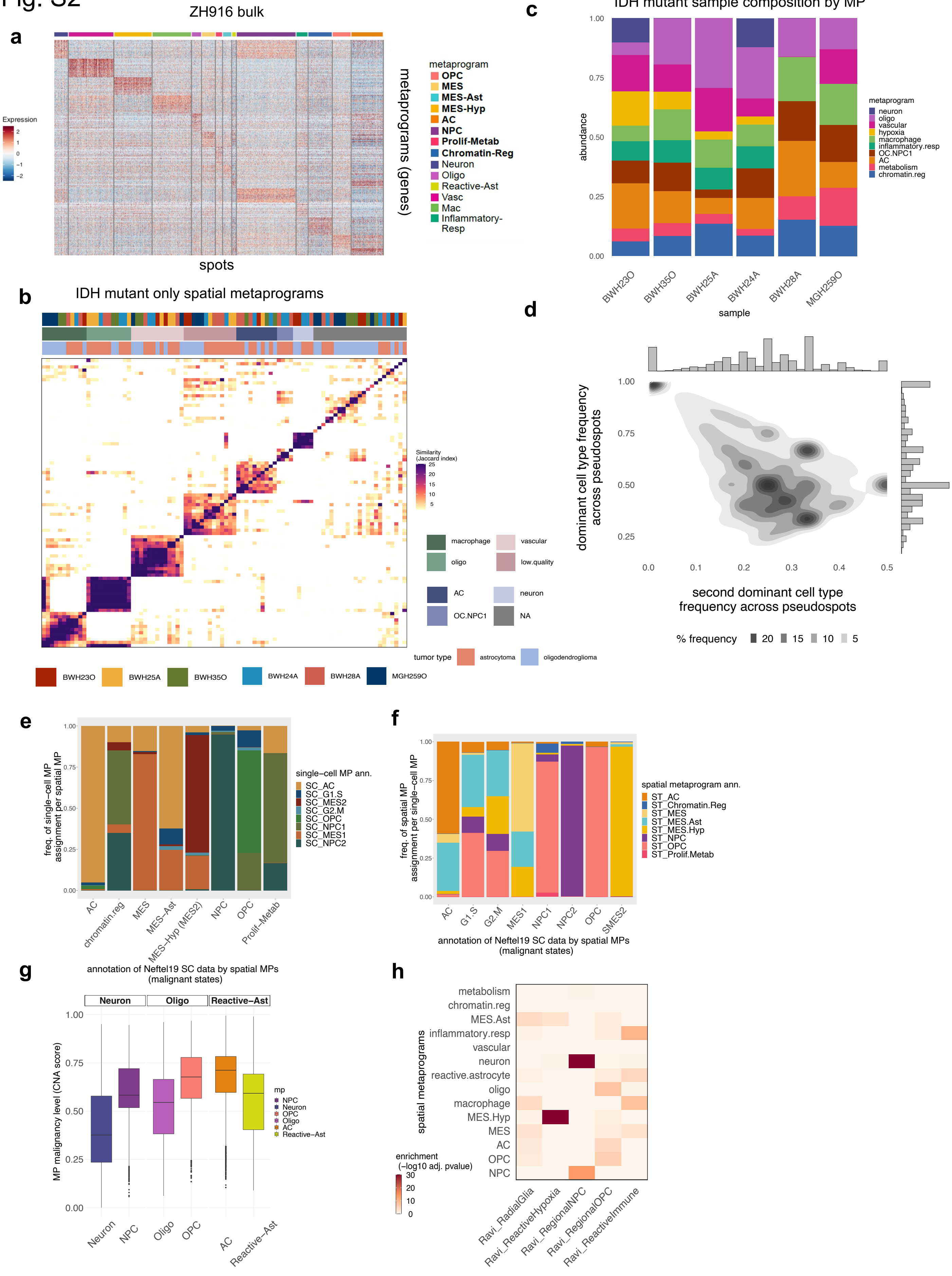
a
ZH916 bulk
Expression
metaprograms (genes)
spots
metaprogram
OPC
MES
MES-Ast
MES-Hyp
AC
NPC
Prolif-Metab
Chromatin-Reg
Neuron
Oligo
Reactive-Ast
Vasc
Mac
Inflammatory-Resp
b
IDH mutant only spatial metaprograms
Similarity (Jaccard index)
macrophage
vascular
oligo
low.quality
AC
neuron
OC.NPC1
NA
tumor type
astrocytoma
oligodendroglioma
BWH23O
BWH25A
BWH35O
BWH24A
BWH28A
MGH259O
c
IDH mutant sample composition by MP
abundance
sample
metaprogram
neuron
oligo
vascular
hypoxia
macrophage
inflammatory.resp
OC.NPC1
AC
metabolism
chromatin.reg
d
dominant cell type frequency across pseudospots
second dominant cell type frequency across pseudospots
% frequency
e
freq. of single-cell MP assignment per spatial MP
single-cell MP ann.
SC_AC
SC_G1.S
SC_MES2
SC_G2.M
SC_OPC
SC_NPC1
SC_MES1
SC_NPC2
AC
chromatin.reg
MES
MES-Ast
MES-Hyp (MES2)
NPC
OPC
Prolif-Metab
f
freq. of spatial MP assignment per single-cell MP
spatial metaprogram ann.
ST_AC
ST_Chromatin.Reg
ST_MES
ST_MES.Ast
ST_MES.Hyp
ST_NPC
ST_OPC
ST_Prolif.Metab
G1.S
G2.M
MES1
NPC1
NPC2
SMES2
annotation of Neftel19 SC data by spatial MPs (malignant states)
g
annotation of Neftel19 SC data by spatial MPs (malignant states)
Neuron
Oligo
Reactive-Ast
MP malignancy level (CNA score)
mp
NPC
Neuron
OPC
Oligo
AC
Reactive-Ast
h
spatial metaprograms
metabolism
chromatin.reg
MES.Ast
inflammatory.resp
vascular
neuron
reactive.astrocyte
oligo
macrophage
MES.Hyp
MES
AC
OPC
NPC
enrichment (-log10 adj. pvalue)
Ravi_RadialGlia
Ravi_ReactiveHypoxia
Ravi_RegionalNPC
Ravi_RegionalOPC
Ravi_ReactiveImmune

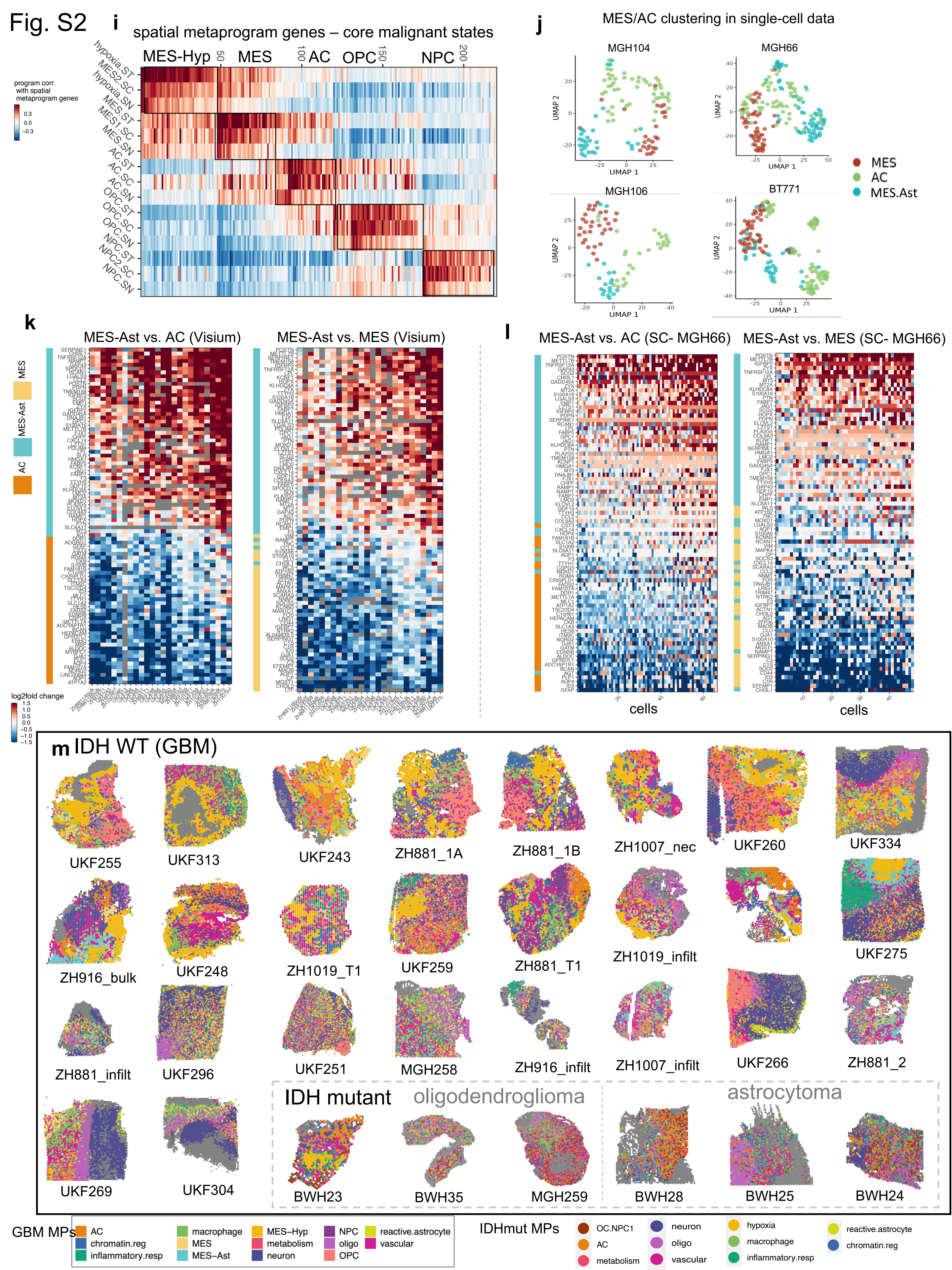
Fig. S2
i
spatial metaprogram genes – core malignant states
MES-Hyp
MES
AC
OPC
NPC
program corr. with spatial metaprogram genes
j
MES/AC clustering in single-cell data
MGH104
MGH66
MGH106
BT771
MES
AC
MES.Ast
k
MES-Ast vs. AC (Visium)
MES-Ast vs. MES (Visium)
log2fold change
l
MES-Ast vs. AC (SC- MGH66)
MES-Ast vs. MES (SC- MGH66)
cells
m IDH WT (GBM)
UKF255
UKF313
UKF243
ZH881_1A
ZH881_1B
ZH1007_nec
UKF260
UKF334
ZH916_bulk
UKF248
ZH1019_T1
UKF259
ZH881_T1
ZH1019_infilt
UKF275
ZH881_infilt
UKF296
UKF251
MGH258
ZH916_infilt
ZH1007_infilt
UKF266
ZH881_2
UKF269
UKF304
IDH mutant
oligodendroglioma
astrocytoma
BWH23
BWH35
MGH259
BWH28
BWH25
BWH24
GBM MPs
AC
chromatin.reg
inflammatory.resp
macrophage
MES
MES–Ast
MES–Hyp
metabolism
neuron
NPC
oligo
OPC
reactive.astrocyte
vascular
IDHmut MPs
OC.NPC1
AC
metabolism
neuron
oligo
vascular
hypoxia
macrophage
inflammatory.resp
reactive.astrocyte
chromatin.reg

**Fig S2. Spatial GBM metaprograms**

**(a)** Spots vs. MP genes in one sample (ZH916), values are log-transformed normalized expression. **(b)** Clustering of all Leiden and NMF gene programs across all IDH mutant samples clustered by gene overlap (Jaccard index) in order to derive IDH mutant metaprograms. **(c)** Composition of IDH mutant samples by MP frequency. Final IDH mutant metaprograms were generated by integrating metaprograms derived by unsupervised analysis as shown in b) with GBM metaprograms that were robust in IDH mutant samples (Methods). **(d)** CODEX pseudospot frequencies of dominant cell types. The distribution map shows the joint frequency of the top two dominant cell types within a pseudospot. Y-axis corresponds to the top dominant cell type frequency within a pseudospot. X-axis corresponds to the second most dominant cell type frequency within a pseudospot. **(e)** GBM single cells (Neftel et al. 2019) were assigned to spatial MPs. Graph shows composition of each spatial MP by their original single-cell MP annotation. **(f)** Similar to but showing composition of Neftel annotated single cells by their spatial MP annotation. **(g)** MP malignancy level (calculated from CNA score for all spots of a given MP across samples, see Methods) in neurodevelopmental malignant states vs. their normal brain counterparts. **(h)** Overlap of GBM spatial MPs with the five GBM spatial regional states defined by Ravi et al. 2022 (enrichment calculated by hypergeometric test). **(i)** Correlations between core malignant spatial metaprograms (per gene) with the equivalent metaprogram across each platform (ST= spatial transcriptomics, SC= single-cell RNA-seq, SN=single-nuclei RNA-seq). For each platform, GBM data of that type was used to calculate the correlations). **(j)** Leiden clustering of all MES and AC annotated single cells in four tumors from Neftel et al. 2019 forming three discrete clusters. **(k)** Relative expression of AC and MES-Ast genes (left) and MES and MES-Ast genes (right) in pseudo-bulk generated from mean expression profiles of each Visium sample. **(l)** Relative expression of AC and MES-Ast genes (left) and MES and MES-Ast genes (right) in scRNA-Seq data from two tumors in Neftel et al. 2019. **(m)** Spatial maps showing spot annotation by MP. Spots were scored for MPs and annotated by their maximum score. Samples are ordered by hypoxia abundance.

Fig. S3

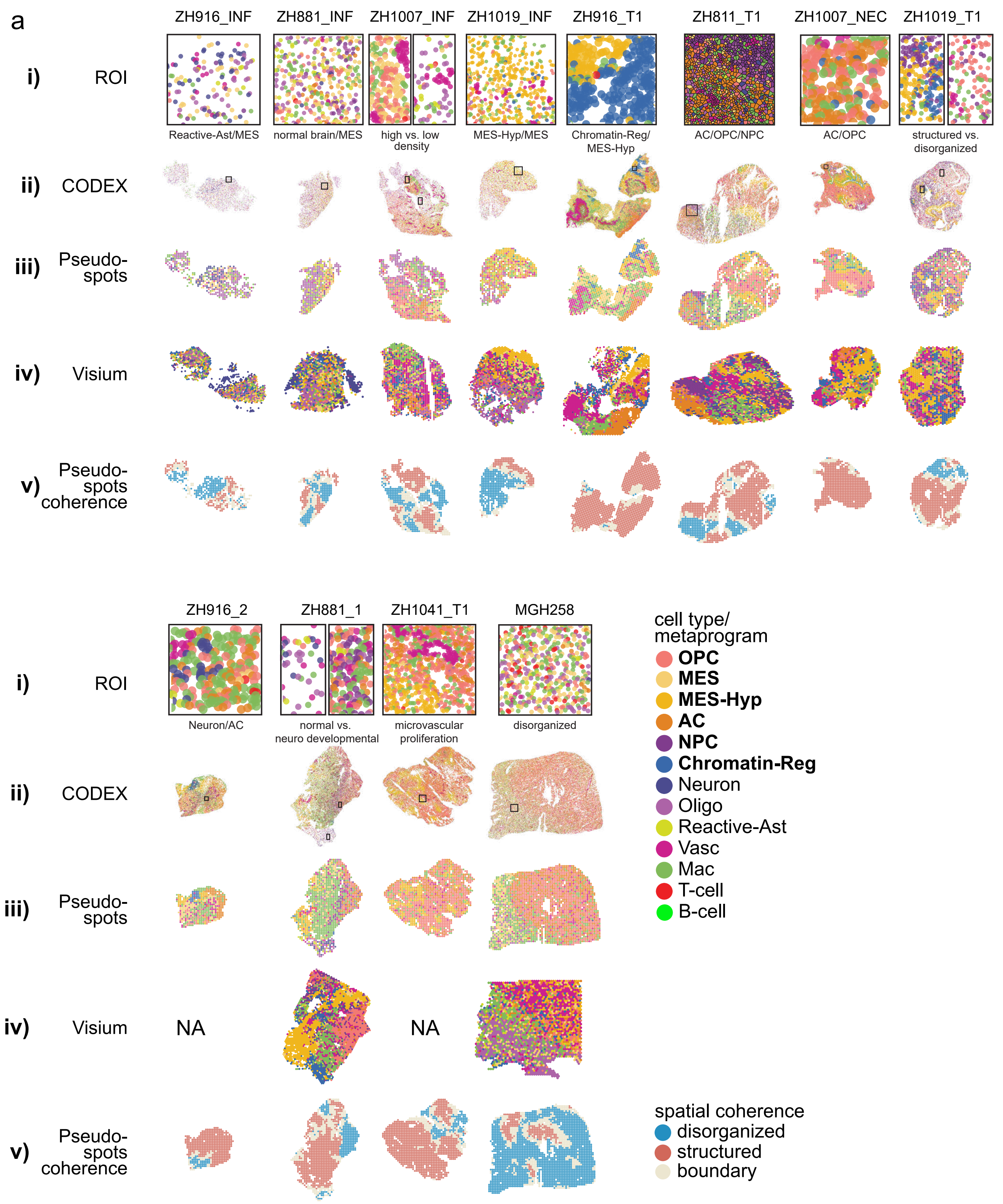
a
ZH916_INF
ZH881_INF
ZH1007_INF
ZH1019_INF
ZH916_T1
ZH811_T1
ZH1007_NEC
ZH1019_T1
i) ROI
Reactive-Ast/MES
normal brain/MES
high vs. low density
MES-Hyp/MES
Chromatin-Reg/ MES-Hyp
AC/OPC/NPC
AC/OPC
structured vs. disorganized
ii) CODEX
iii) Pseudo-spots
iv) Visium
v) Pseudo-spots coherence
ZH916_2
ZH881_1
ZH1041_T1
MGH258
Neuron/AC
normal vs. neuro developmental
microvascular proliferation
disorganized
NA
NA
cell type/ metaprogram
OPC
MES
MES-Hyp
AC
NPC
Chromatin-Reg
Neuron
Oligo
Reactive-Ast
Vasc
Mac
T-cell
B-cell
spatial coherence
disorganized
structured
boundary

Fig. S3

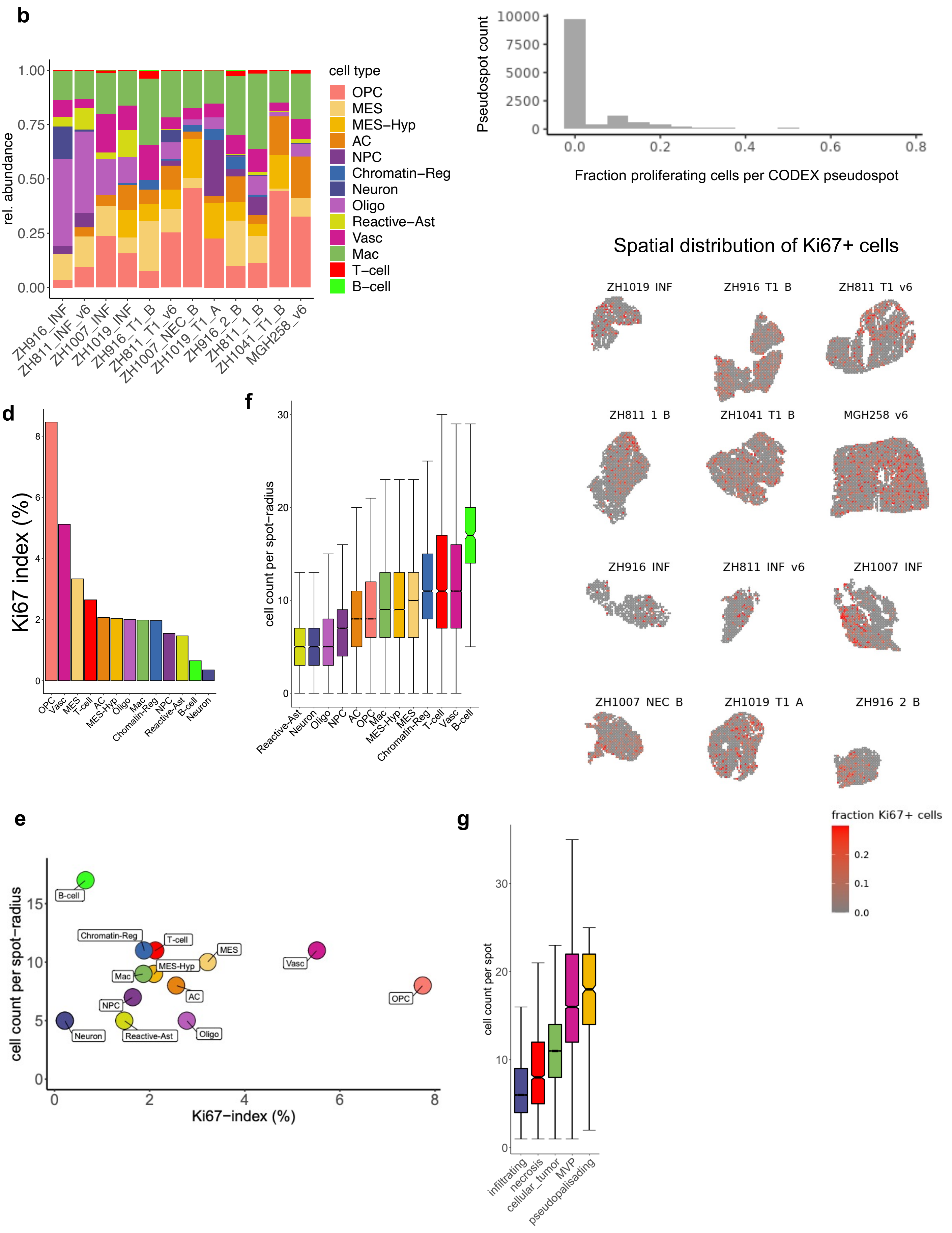

b
rel. abundance
1.00
0.75
0.50
0.25
0.00
ZH916_INF
ZH811_INF_v6
ZH1007_INF
ZH1019_INF
ZH916_T1_B
ZH811_T1_v6
ZH1007_NEC_B
ZH1019_T1_A
ZH916_2_B
ZH811_1_B
ZH1041_T1_B
MGH258_v6
cell type
OPC
MES
MES−Hyp
AC
NPC
Chromatin−Reg
Neuron
Oligo
Reactive−Ast
Vasc
Mac
T−cell
B−cell
c
Pseudospot count
10000
7500
5000
2500
0
0.0
0.2
0.4
0.6
0.8
Fraction proliferating cells per CODEX pseudospot
Spatial distribution of Ki67+ cells
ZH1019 INF
ZH916 T1 B
ZH811 T1 v6
ZH811 1 B
ZH1041 T1 B
MGH258 v6
ZH916 INF
ZH811 INF v6
ZH1007 INF
ZH1007 NEC B
ZH1019 T1 A
ZH916 2 B
fraction Ki67+ cells
0.2
0.1
0.0
d
Ki67 index (%)
8
6
4
2
0
OPC
Vasc
MES
T-cell
AC
MES-Hyp
Oligo
Mac
Chromatin-Reg
NPC
Reactive-Ast
B-cell
Neuron
f
cell count per spot-radius
30
20
10
0
Reactive-Ast
Neuron
Oligo
NPC
AC
OPC
Mac
MES-Hyp
MES
Chromatin-Reg
T-cell
Vasc
B-cell
e
cell count per spot−radius
15
10
5
0
B-cell
Chromatin-Reg
T-cell
MES
Vasc
Mac
MES-Hyp
AC
OPC
NPC
Neuron
Reactive-Ast
Oligo
0
2
4
6
8
Ki67−index (%)
g
cell count per spot
30
20
10
0
infiltrating
necrosis
cellular_tumor
MVP
pseudopalisading

**Figure S3. GBM profiling by CODEX**

**(a)** Spatial maps of CODEX samples: (i) Region of interest (ROI) derived from the CODEX single cell spatial maps (ii) highlighting the indicated cell relationships. (iii) Corresponding CODEX pseudo-spots annotated by the dominating cell type per pseudo-spot (iv) Corresponding near-adjacent sections: Visium spots annotated by the dominant MP derived from gene expression. Unique Visium MPs were excluded from this analysis (Prolif-Metab, Inflammatory-Mac, MES-Ast). (v) Results of spatial coherence classification on CODEX pseudo-spots. **(b)** Relative cell type abundance per sample. **(c)** Cell cycle analysis. Top: Histogram showing the spot count in the function of fraction proliferating cells per pseudo-spot. Note: Only 163 of 13.058 spots (1.2%) contain more than 40% cycling cells. Bottom: Spatial map of CODEX pseudo-spots with color representing the fraction of cycling cells per spot. Note: Most samples show local hotspots of proliferation. **(d)** Ki-67 proliferation index per cell type across all samples. **(e)** Scatter plot showing Ki67 proliferation index and cell count per spot-radius measurements for all cell types. **(f)** Boxplot of cell density (cell count per 55 µm diameter). **(g)** Boxplot of cell density by pathology annotation from H&E.

Fig. S4

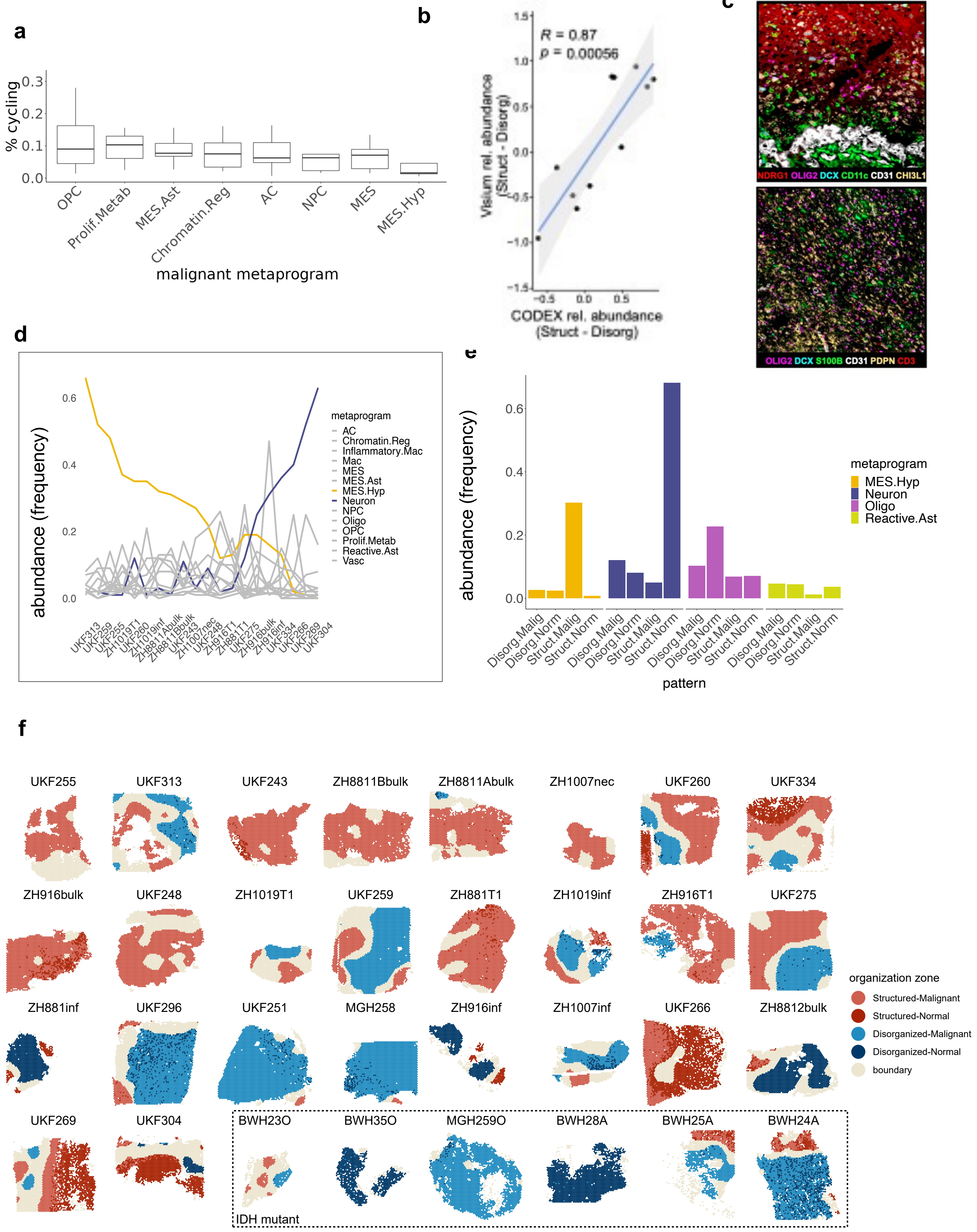
a
% cycling
0.0
0.1
0.2
0.3
OPC
Prolif.Metab
MES.Ast
Chromatin.Reg
AC
NPC
MES
MES.Hyp
malignant metaprogram
b
R = 0.87
p = 0.00056
Visium rel. abundance (Struct - Disorg)
CODEX rel. abundance (Struct - Disorg)
c
CODEX image
NDRG1 OLIG2 DCX CD11c CD31 CHI3L1
OLIG2 DCX S100B CD31 PDPN CD3
d
abundance (frequency)
0.0
0.2
0.4
0.6
metaprogram
AC
Chromatin.Reg
Inflammatory.Mac
Mac
MES
MES.Ast
MES.Hyp
Neuron
NPC
Oligo
OPC
Prolif.Metab
Reactive.Ast
Vasc
UKF313
UKF259
UKF255
ZH1019T1
UKF260
ZH1019inf
ZH8811Abulk
ZH8811Bbulk
UKF243
ZH1007nec
UKF248
ZH916T1
ZH881T1
UKF275
ZH916bulk
ZH916inf
UKF334
UKF266
UKF269
UKF304
e
abundance (frequency)
0.0
0.2
0.4
0.6
metaprogram
MES.Hyp
Neuron
Oligo
Reactive.Ast
Disorg.Malig
Disorg.Norm
Struct.Malig
Struct.Norm
pattern
f
UKF255
UKF313
UKF243
ZH8811Bbulk
ZH8811Abulk
ZH1007nec
UKF260
UKF334
ZH916bulk
UKF248
ZH1019T1
UKF259
ZH881T1
ZH1019inf
ZH916T1
UKF275
ZH881inf
UKF296
UKF251
MGH258
ZH916inf
ZH1007inf
UKF266
ZH8812bulk
UKF269
UKF304
BWH23O
BWH35O
MGH259O
BWH28A
BWH25A
BWH24A
IDH mutant
organization zone
Structured-Malignant
Structured-Normal
Disorganized-Malignant
Disorganized-Normal
boundary

Fig. S4

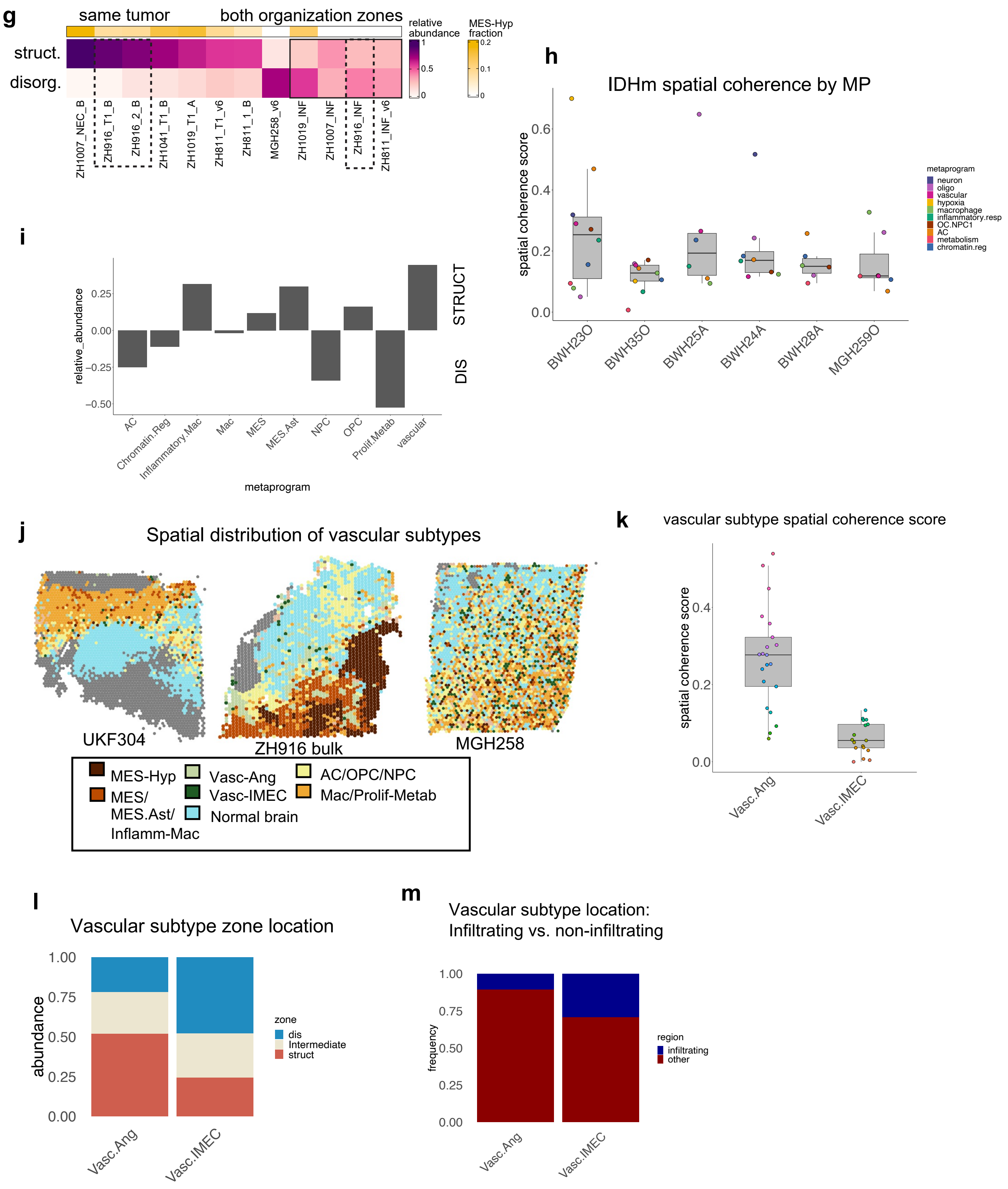
g
same tumor
both organization zones
relative abundance
1
0.5
0
MES-Hyp fraction
0.2
0.1
0
struct.
disorg.
ZH1007_NEC_B
ZH916_T1_B
ZH916_2_B
ZH1041_T1_B
ZH1019_T1_A
ZH811_T1_v6
ZH811_1_B
MGH258_v6
ZH1019_INF
ZH1007_INF
ZH916_INF
ZH811_INF_v6
h
IDHm spatial coherence by MP
spatial coherence score
0.0
0.2
0.4
0.6
BWH23O
BWH35O
BWH25A
BWH24A
BWH28A
MGH259O
metaprogram
neuron
oligo
vascular
hypoxia
macrophage
inflammatory.resp
OC.NPC1
AC
metabolism
chromatin.reg
i
relative_abundance
0.25
0.00
−0.25
−0.50
STRUCT
DIS
AC
Chromatin.Reg
Inflammatory.Mac
Mac
MES
MES.Ast
NPC
OPC
Prolif.Metab
vascular
metaprogram
j
Spatial distribution of vascular subtypes
UKF304
ZH916 bulk
MGH258
MES-Hyp
MES/ MES.Ast/ Inflamm-Mac
Vasc-Ang
Vasc-IMEC
Normal brain
AC/OPC/NPC
Mac/Prolif-Metab
k
vascular subtype spatial coherence score
spatial coherence score
0.0
0.2
0.4
Vasc.Ang
Vasc.IMEC
l
Vascular subtype zone location
abundance
1.00
0.75
0.50
0.25
0.00
Vasc.Ang
Vasc.IMEC
zone
dis
Intermediate
struct
m
Vascular subtype location: Infiltrating vs. non-infiltrating
frequency
1.00
0.75
0.50
0.25
0.00
Vasc.Ang
Vasc.IMEC
region
infiltrating
other

**Figure S4. Spatial coherence and organizational zones (a)** Boxplot showing the percentage of spots with a strong cell cycle signal (see Methods), across spots annotated for a given malignant state. **(b)** Correlation between relative abundance of structured spots vs. disorganized spots in near adjacent samples in Visium and CODEX ($R$=0.87, $p$=0.00056). **(c)** Example of a structured and disorganized region by immunostaining (corresponding to image in Fig. 4e). **(d)** Abundance of MP across structured samples. MES-Hyp and Neuron MPs are colored. **(e)** Mean frequency of MES-Hyp, Neuron, Oligo, and Reactive-Ast spots per spatial organization zone across all GBM samples. **(f)** Spatial maps of all Visium samples annotated by organizational zones (Struct-Malig, Struct-Norm, Disorg-Malig, Disorg-Norm) (Methods). **(g)** Abundance of structured vs. disorganized calculated from spatial coherence in CODEX samples. Annotation bar corresponds to MES-Hyp abundance. **(h**) MP spatial coherence per sample in IDH mutant samples. Each dot represents mean spatial coherence of all spots annotated with a given MP per sample. **(i)** Compositional differences in structured vs. disorganized regions (across all samples). Vasc (enriched in structured, $p$=0.03) and Prolif-Metab (enriched in disorganized, $p$=0.003) were significant. **(j)** Spatial maps highlighting the different spatial patterns and distribution of the two vascular subcluster, Vasc-Ang (light green) and Vasc-IMEC (dark green). **(k)** Spatial coherence score per vascular subcluster. Each dot presents the mean spatial coherence score for spots of a given vascular sub cluster per sample (p-value < 0.001). **(l)** Organization zone composition of each vascular subcluster (across all GBM samples). **(m)** Distribution of vascular subtypes in infiltrating vs. non-infiltrating areas (Methods).

Fig. S5

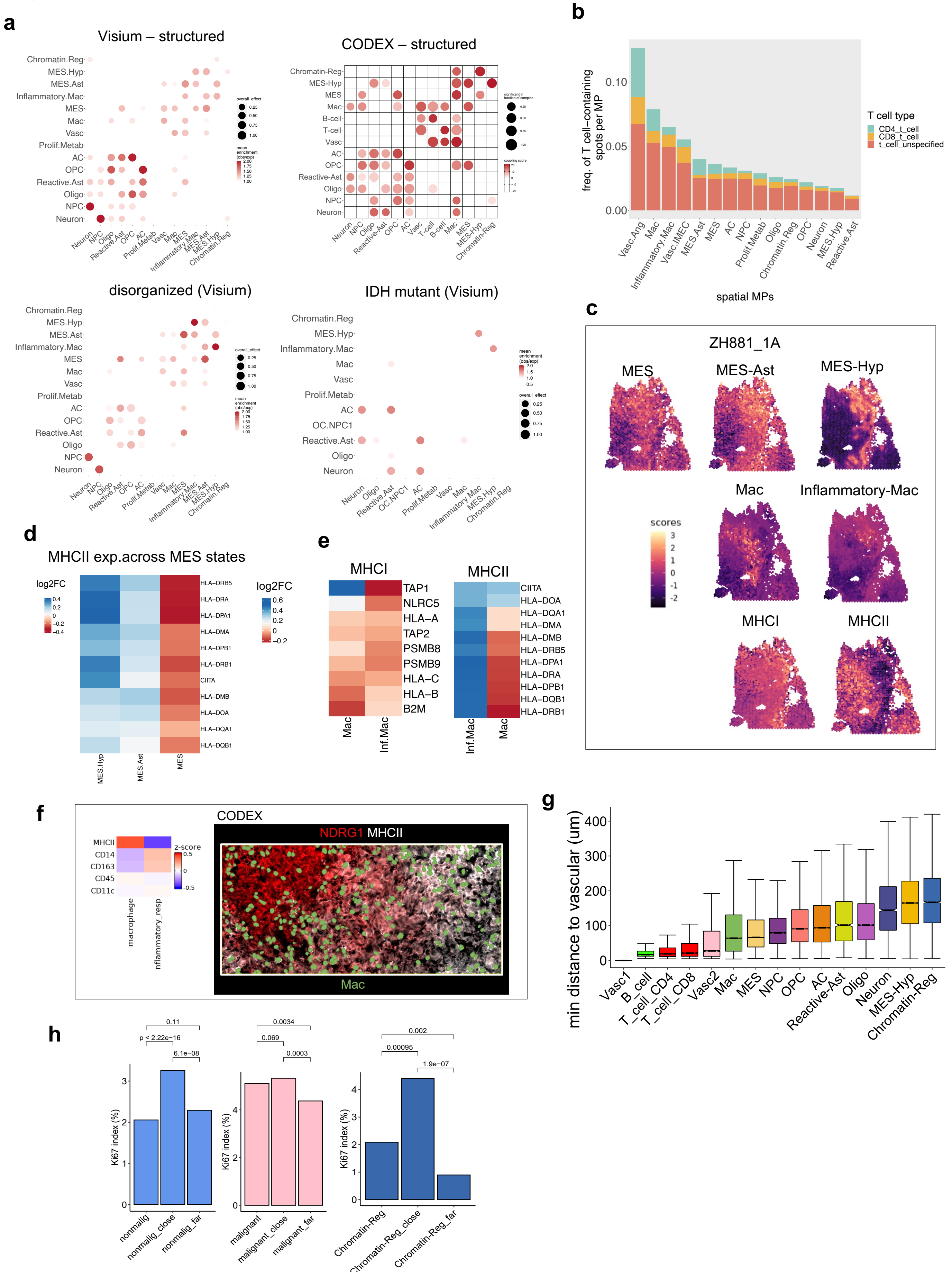
a
Visium – structured
CODEX – structured
disorganized (Visium)
IDH mutant (Visium)
Chromatin.Reg
MES.Hyp
MES.Ast
Inflammatory.Mac
MES
Mac
Vasc
Prolif.Metab
AC
OPC
Reactive.Ast
Oligo
NPC
Neuron
OC.NPC1
B-cell
T-cell
overall_effect
mean enrichment (obs/exp)
significant in fraction of samples
coupling score
b
freq. of T cell–containing spots per MP
spatial MPs
T cell type
CD4_t_cell
CD8_t_cell
t_cell_unspecified
Vasc.Ang
Vasc.IMEC
c
ZH881_1A
MES
MES-Ast
MES-Hyp
Mac
Inflammatory-Mac
MHCI
MHCII
scores
d
MHCII exp.across MES states
log2FC
HLA–DRB5
HLA–DRA
HLA–DPA1
HLA–DMA
HLA–DPB1
HLA–DRB1
CIITA
HLA–DMB
HLA–DOA
HLA–DQA1
HLA–DQB1
MES.Hyp
MES.Ast
MES
e
MHCI
MHCII
log2FC
TAP1
NLRC5
HLA–A
TAP2
PSMB8
PSMB9
HLA–C
HLA–B
B2M
CIITA
HLA–DOA
HLA–DQA1
HLA–DMA
HLA–DMB
HLA–DRB5
HLA–DPA1
HLA–DRA
HLA–DPB1
HLA–DQB1
HLA–DRB1
Mac
Inf.Mac
f
CODEX
MHCII
CD14
CD163
CD45
CD11c
z-score
macrophage
nflammatory_resp
NDRG1 MHCII
Mac
g
min distance to vascular (um)
Vasc1
B_cell
T_cell_CD4
T_cell_CD8
Vasc2
Mac
MES
NPC
OPC
AC
Reactive-Ast
Oligo
Neuron
MES-Hyp
Chromatin-Reg
h
Ki67 index (%)
p < 2.22e−16
0.11
6.1e−08
0.0034
0.069
0.0003
0.002
0.00095
1.9e−07
nonmalig
nonmalig_close
nonmalig_far
malignant
malignant_close
malignant_far
Chromatin-Reg
Chromatin-Reg_close
Chromatin-Reg_far

**Figure S5. Spatial relationship analysis**

**(a)** Colocalization of MP pairs within a spot for structured and disorganized regions in Visium, structured regions in CODEX, and IDH mutant samples. Mean enrichment values represent observed/expected spot colocalization (Methods) and dot size represents the proportion of samples for which a metaprogram pair was enriched (Methods). **(b)** Overall percentage of putative T cell containing spots per MP (across all GBM samples) and further categorization of the T cell containing spots (CD4, CD8, unspecified). T cell containing spots in Visium data were identified by detection of counts for canonical T cell markers (Methods). **(c)** Spatial maps of ZH881_1A bulk scored for mesenchymal MPs, myeloid MPs, and MHC class I and MHC class II genes. **(d)** Relative expression of MHC class II genes across the mesenchymal MPs. **(e)** Relative expression of MHC class I and MHC class II genes across myeloid MPs. Expression values represent average expression values across all GBM samples. **(f)** Left (CODEX): relative differences in protein expression between macrophages within hypoxic (Inflammatory-Mac) and outside hypoxic niches (Mac) for the indicated markers. Right (CODEX): Immunostaining showing macrophages with green segmentation masks. Mac in hypoxic niche ($NDRG1^{high}$) show downregulation of MHCII compared to non-hypoxic area ($NDRG1^{low}$) **(g)** The distance to closest vasc1 cell per cell type. Note: Median distance to MES-Hyp (= 161 µm). **(h)** Ki67-proliferation index for non-malignant (left), malignant cells (center) and Chromatin-Reg (right) in the function of the distance to the next vascular cell (close: < 50 µm, far: > 200 µm). **(i)** Spatial maps of Visium GBM samples depicting spots containing T cells as described in (b) and Methods.

### Fig. S6

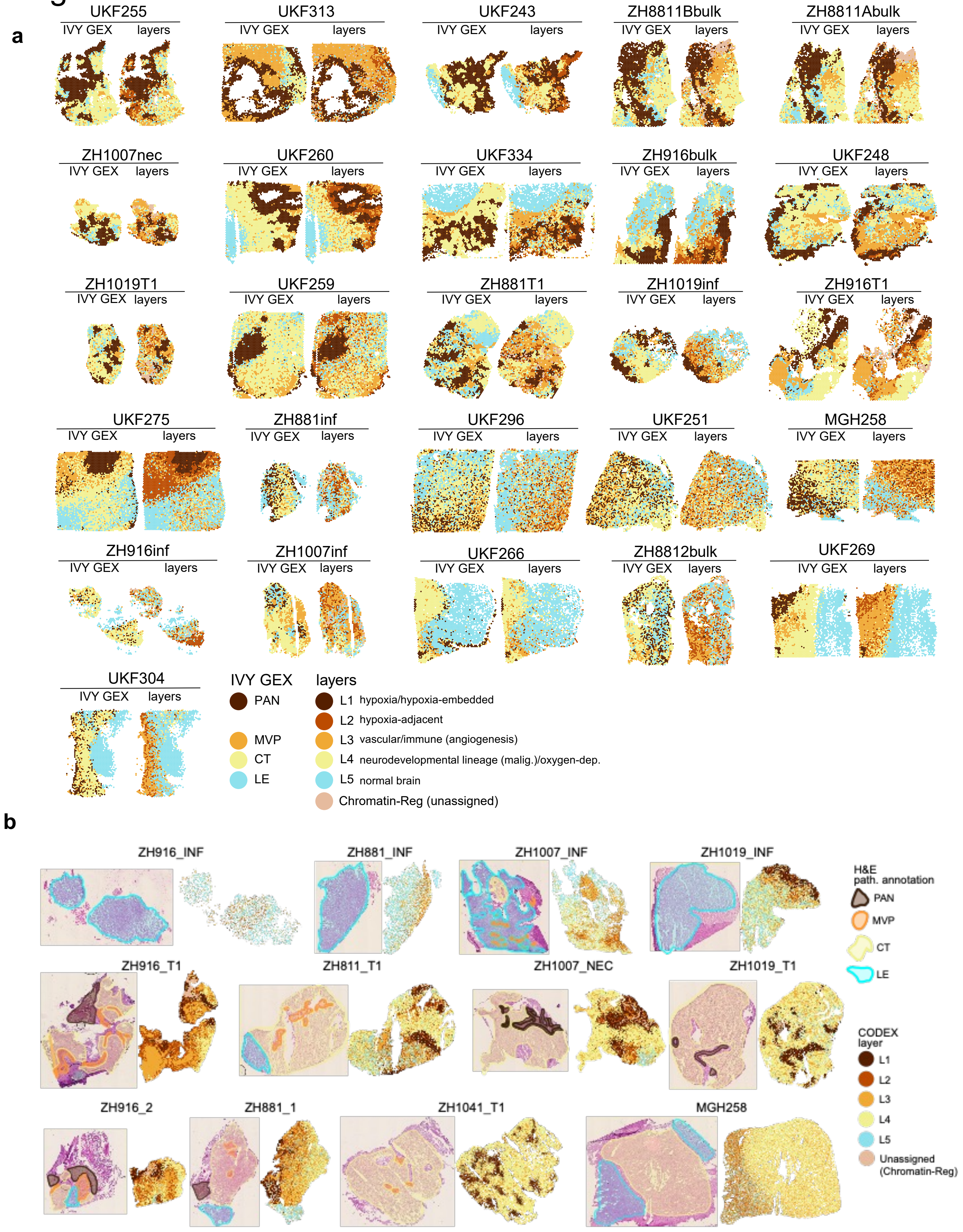
a
UKF255
IVY GEX layers
UKF313
UKF243
ZH8811Bbulk
ZH8811Abulk
ZH1007nec
UKF260
UKF334
ZH916bulk
UKF248
ZH1019T1
UKF259
ZH881T1
ZH1019inf
ZH916T1
UKF275
ZH881inf
UKF296
UKF251
MGH258
ZH916inf
ZH1007inf
UKF266
ZH8812bulk
UKF269
UKF304
IVY GEX
PAN
MVP
CT
LE
layers
L1 hypoxia/hypoxia-embedded
L2 hypoxia-adjacent
L3 vascular/immune (angiogenesis)
L4 neurodevelopmental lineage (malig.)/oxygen-dep.
L5 normal brain
Chromatin-Reg (unassigned)
b
ZH916_INF
ZH881_INF
ZH1007_INF
ZH1019_INF
ZH916_T1
ZH811_T1
ZH1007_NEC
ZH1019_T1
ZH916_2
ZH881_1
ZH1041_T1
MGH258
H&E path. annotation
PAN
MVP
CT
LE
CODEX layer
L1
L2
L3
L4
L5
Unassigned (Chromatin-Reg)

Fig. S6

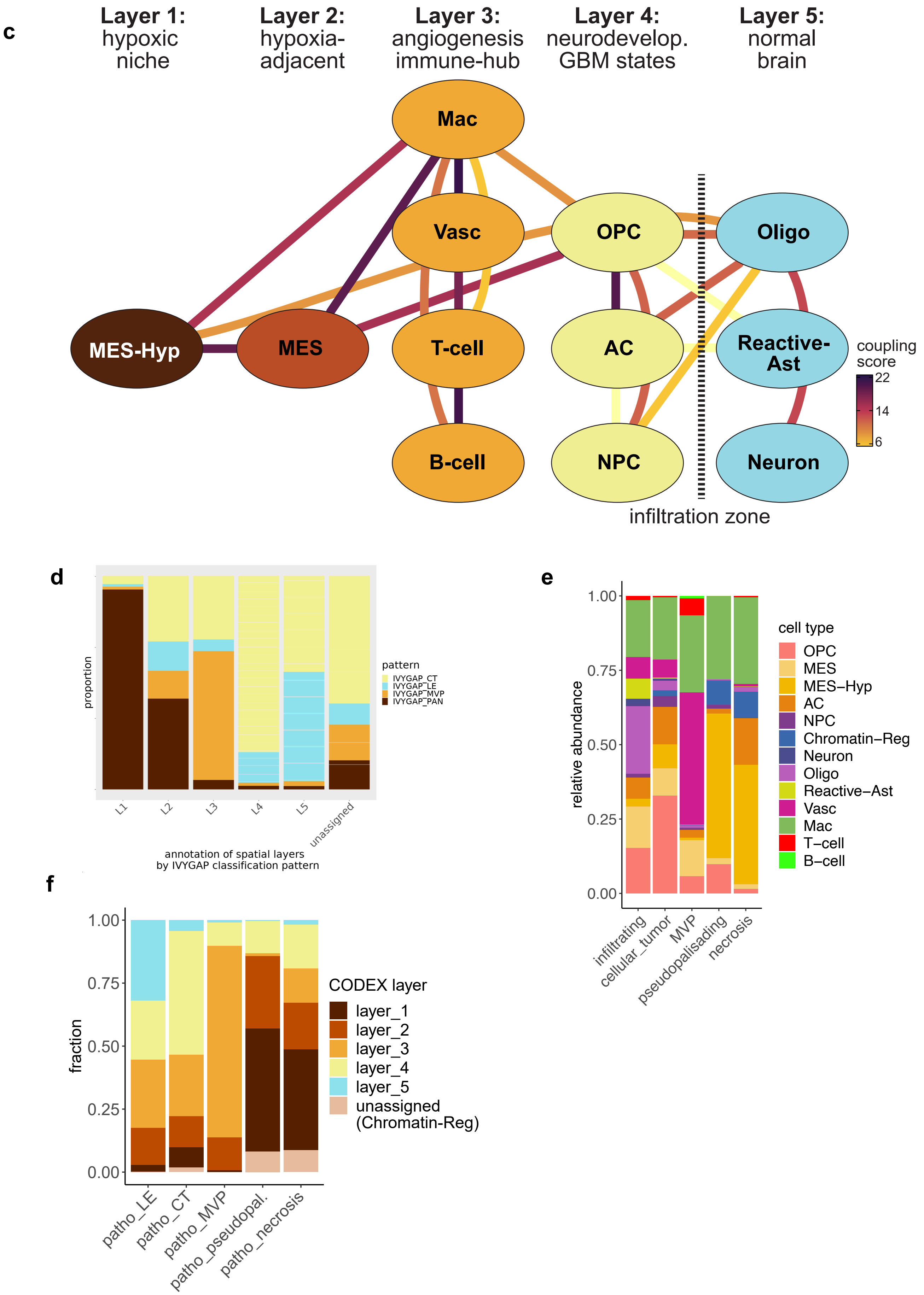
c
Layer 1:
hypoxic
niche
Layer 2:
hypoxia-
adjacent
Layer 3:
angiogenesis
immune-hub
Layer 4:
neurodevelop.
GBM states
Layer 5:
normal
brain
Mac
Vasc
OPC
Oligo
MES-Hyp
MES
T-cell
AC
Reactive-
Ast
B-cell
NPC
Neuron
coupling
score
22
14
6
infiltration zone
d
proportion
pattern
IVYGAP_CT
IVYGAP_LE
IVYGAP_MVP
IVYGAP_PAN
L1
L2
L3
L4
L5
unassigned
annotation of spatial layers
by IVYGAP classification pattern
e
relative abundance
1.00
0.75
0.50
0.25
0.00
infiltrating
cellular_tumor
MVP
pseudopalisading
necrosis
cell type
OPC
MES
MES–Hyp
AC
NPC
Chromatin–Reg
Neuron
Oligo
Reactive–Ast
Vasc
Mac
T–cell
B–cell
f
fraction
1.00
0.75
0.50
0.25
0.00
patho_LE
patho_CT
patho_MVP
patho_pseudopal.
patho_necrosis
CODEX layer
layer_1
layer_2
layer_3
layer_4
layer_5
unassigned
(Chromatin-Reg)

**Figure S6. GBM state layers**

**(a)** Spatial maps of Visium data annotated by state layer. **(b)** H&E stainings with neuro-pathologist histology annotation (left picture of sample pairs) next to CODEX spatial maps color-coded by the layers (right plot of sample pair) as depicted in a). **(c)** Network-graph with nodes representing cell types/states and edges representing mean z-scores of spatial attraction as derived by the CODEX co-localization analysis. Only edges representing recurrent interactions (> 25% of samples) were retained. **(d)** Composition of state layers by their Ivy-GAP transcriptional program annotation (Visium). **(e)** Composition of H&E histological annotations by MP identity (CODEX). **(f)** Compositions of H&E histological annotations by state layer (CODEX).

**Supplementary Note 1**

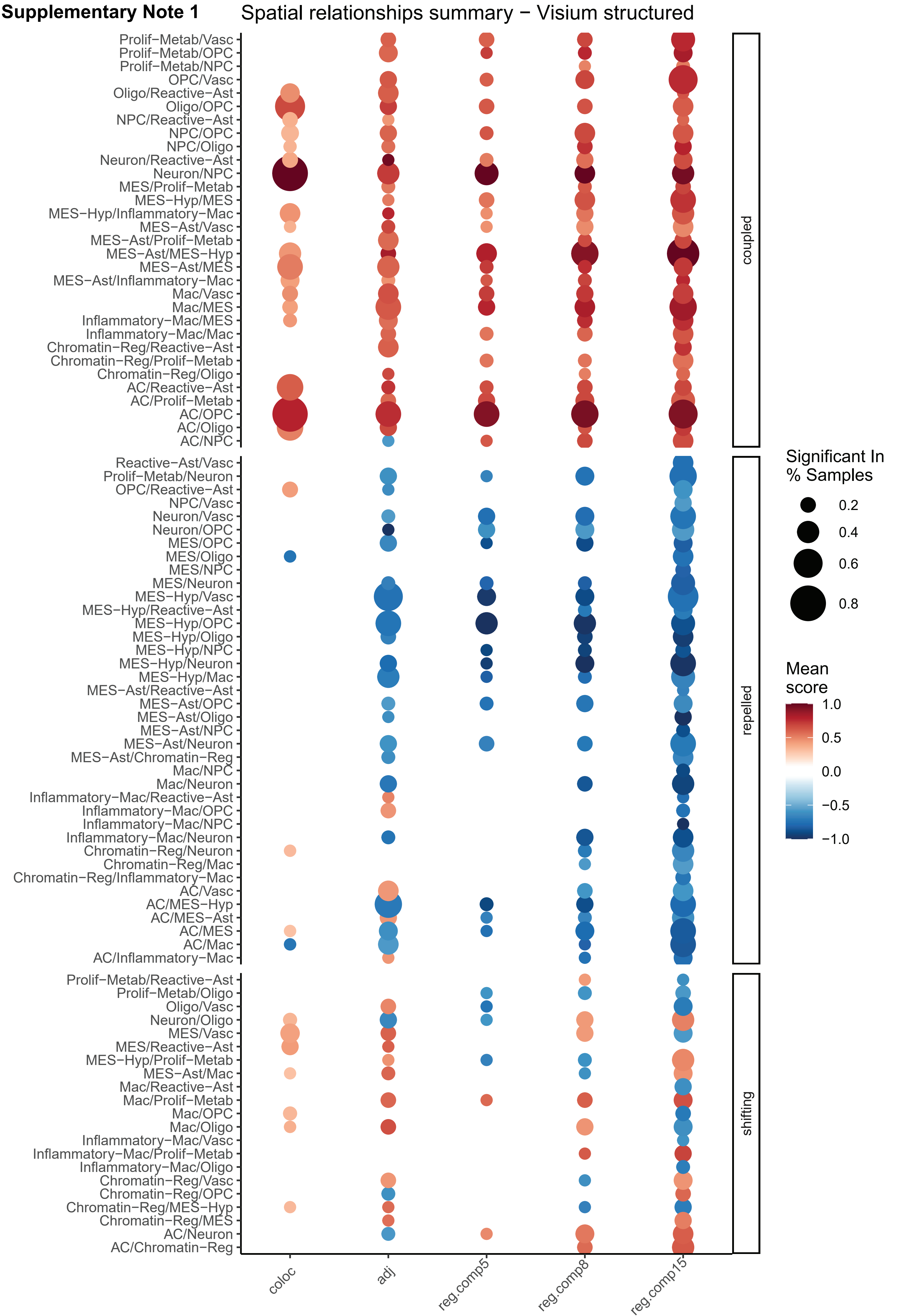

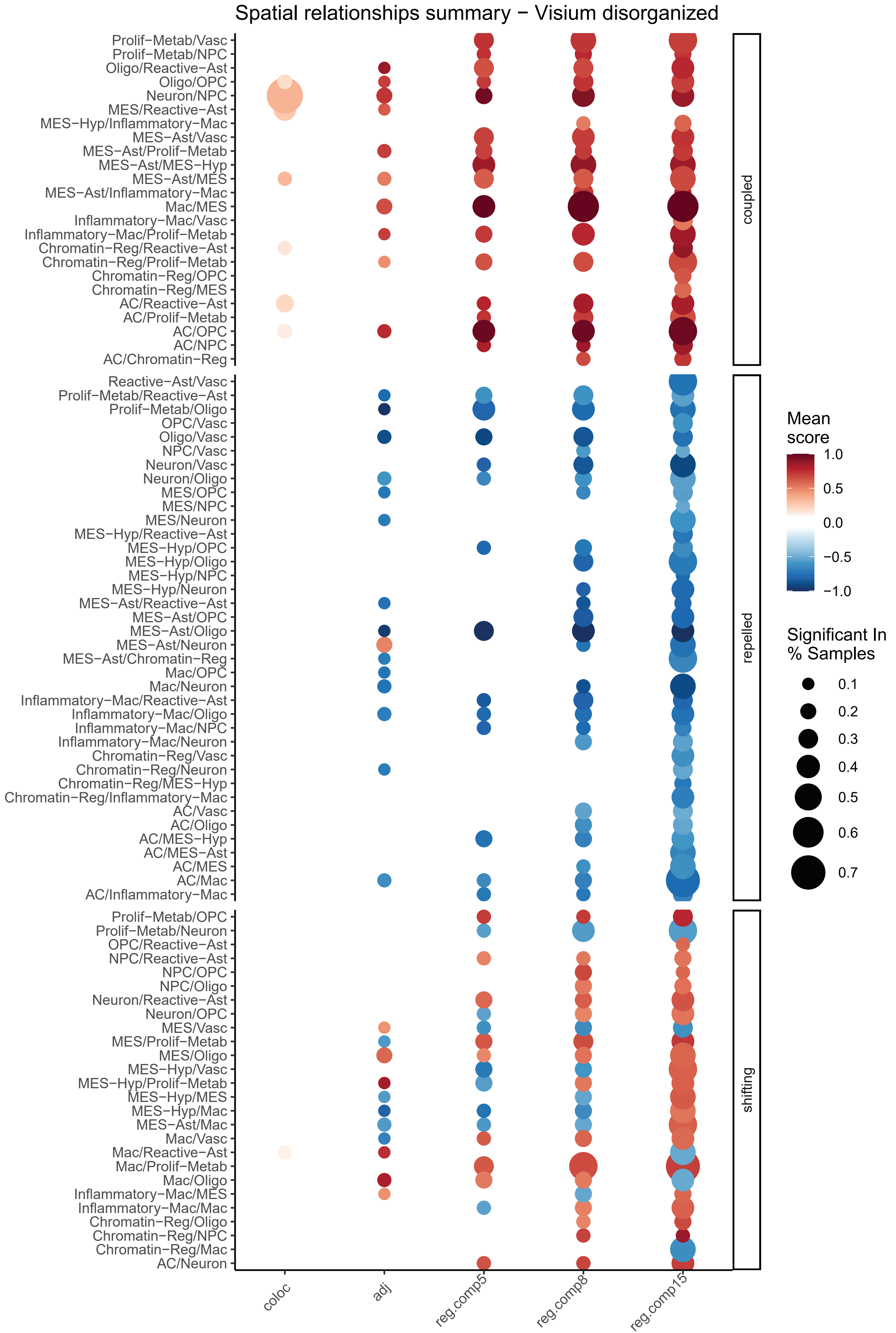

Spatial relationships summary − Visium disorganized
Prolif−Metab/Vasc
Prolif−Metab/NPC
Oligo/Reactive−Ast
Oligo/OPC
Neuron/NPC
MES/Reactive−Ast
MES−Hyp/Inflammatory−Mac
MES−Ast/Vasc
MES−Ast/Prolif−Metab
MES−Ast/MES−Hyp
MES−Ast/MES
MES−Ast/Inflammatory−Mac
Mac/MES
Inflammatory−Mac/Vasc
Inflammatory−Mac/Prolif−Metab
Chromatin−Reg/Reactive−Ast
Chromatin−Reg/Prolif−Metab
Chromatin−Reg/OPC
Chromatin−Reg/MES
AC/Reactive−Ast
AC/Prolif−Metab
AC/OPC
AC/NPC
AC/Chromatin−Reg
Reactive−Ast/Vasc
Prolif−Metab/Reactive−Ast
Prolif−Metab/Oligo
OPC/Vasc
Oligo/Vasc
NPC/Vasc
Neuron/Vasc
Neuron/Oligo
MES/OPC
MES/NPC
MES/Neuron
MES−Hyp/Reactive−Ast
MES−Hyp/OPC
MES−Hyp/Oligo
MES−Hyp/NPC
MES−Hyp/Neuron
MES−Ast/Reactive−Ast
MES−Ast/OPC
MES−Ast/Oligo
MES−Ast/Neuron
MES−Ast/Chromatin−Reg
Mac/OPC
Mac/Neuron
Inflammatory−Mac/Reactive−Ast
Inflammatory−Mac/Oligo
Inflammatory−Mac/NPC
Inflammatory−Mac/Neuron
Chromatin−Reg/Vasc
Chromatin−Reg/Neuron
Chromatin−Reg/MES−Hyp
Chromatin−Reg/Inflammatory−Mac
AC/Vasc
AC/Oligo
AC/MES−Hyp
AC/MES−Ast
AC/MES
AC/Mac
AC/Inflammatory−Mac
Prolif−Metab/OPC
Prolif−Metab/Neuron
OPC/Reactive−Ast
NPC/Reactive−Ast
NPC/OPC
NPC/Oligo
Neuron/Reactive−Ast
Neuron/OPC
MES/Vasc
MES/Prolif−Metab
MES/Oligo
MES−Hyp/Vasc
MES−Hyp/Prolif−Metab
MES−Hyp/MES
MES−Hyp/Mac
MES−Ast/Mac
Mac/Vasc
Mac/Reactive−Ast
Mac/Prolif−Metab
Mac/Oligo
Inflammatory−Mac/MES
Inflammatory−Mac/Mac
Chromatin−Reg/Oligo
Chromatin−Reg/NPC
Chromatin−Reg/Mac
AC/Neuron
coupled
repelled
shifting
coloc
adj
reg.comp5
reg.comp8
reg.comp15
Mean score
1.0
0.5
0.0
−0.5
−1.0
Significant In % Samples
0.1
0.2
0.3
0.4
0.5
0.6
0.7

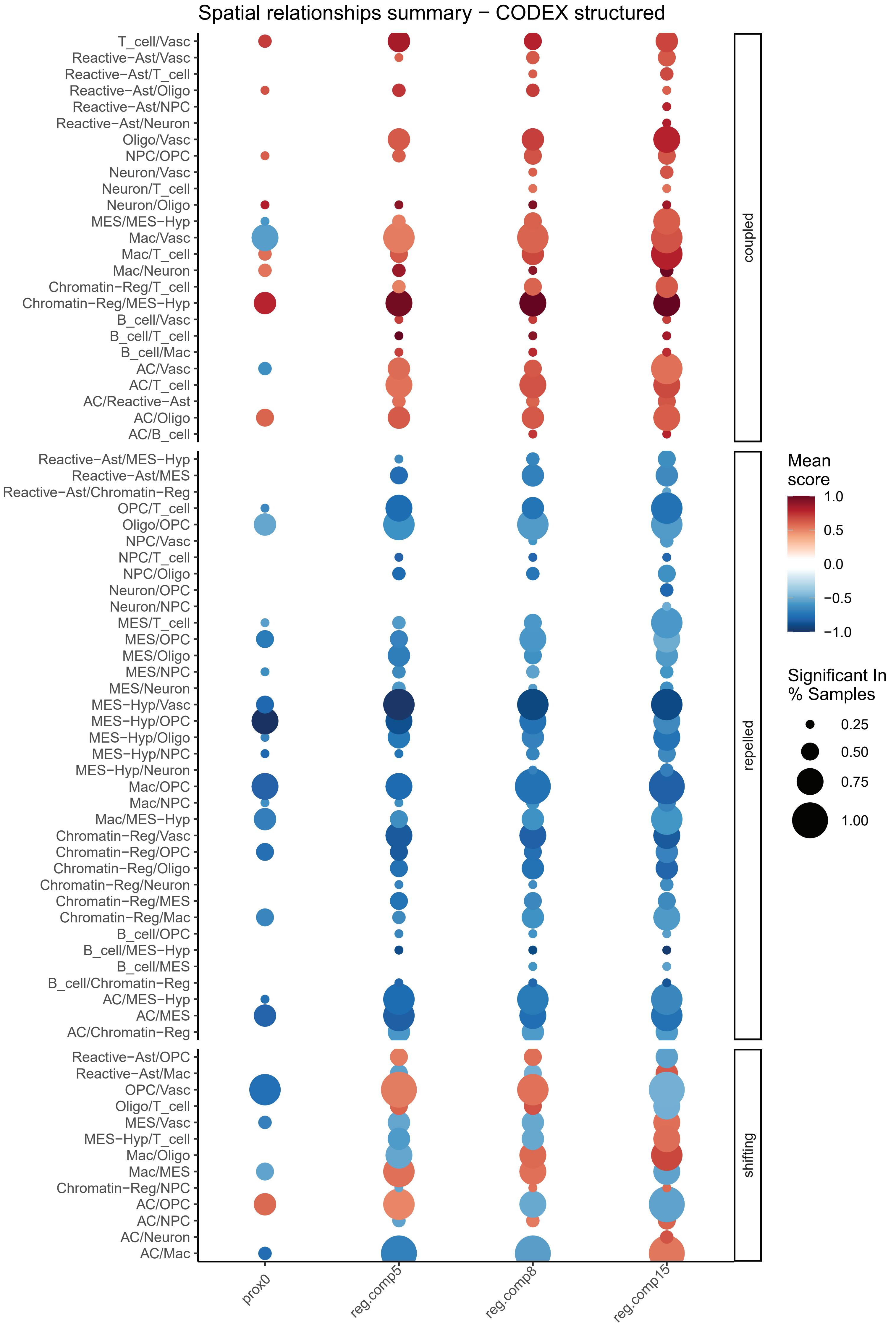
Spatial relationships summary − CODEX structured
T_cell/Vasc
Reactive−Ast/Vasc
Reactive−Ast/T_cell
Reactive−Ast/Oligo
Reactive−Ast/NPC
Reactive−Ast/Neuron
Oligo/Vasc
NPC/OPC
Neuron/Vasc
Neuron/T_cell
Neuron/Oligo
MES/MES−Hyp
Mac/Vasc
Mac/T_cell
Mac/Neuron
Chromatin−Reg/T_cell
Chromatin−Reg/MES−Hyp
B_cell/Vasc
B_cell/T_cell
B_cell/Mac
AC/Vasc
AC/T_cell
AC/Reactive−Ast
AC/Oligo
AC/B_cell
coupled
Reactive−Ast/MES−Hyp
Reactive−Ast/MES
Reactive−Ast/Chromatin−Reg
OPC/T_cell
Oligo/OPC
NPC/Vasc
NPC/T_cell
NPC/Oligo
Neuron/OPC
Neuron/NPC
MES/T_cell
MES/OPC
MES/Oligo
MES/NPC
MES/Neuron
MES−Hyp/Vasc
MES−Hyp/OPC
MES−Hyp/Oligo
MES−Hyp/NPC
MES−Hyp/Neuron
Mac/OPC
Mac/NPC
Mac/MES−Hyp
Chromatin−Reg/Vasc
Chromatin−Reg/OPC
Chromatin−Reg/Oligo
Chromatin−Reg/Neuron
Chromatin−Reg/MES
Chromatin−Reg/Mac
B_cell/OPC
B_cell/MES−Hyp
B_cell/MES
B_cell/Chromatin−Reg
AC/MES−Hyp
AC/MES
AC/Chromatin−Reg
repelled
Reactive−Ast/OPC
Reactive−Ast/Mac
OPC/Vasc
Oligo/T_cell
MES/Vasc
MES−Hyp/T_cell
Mac/Oligo
Mac/MES
Chromatin−Reg/NPC
AC/OPC
AC/NPC
AC/Neuron
AC/Mac
shifting
prox0
reg.comp5
reg.comp8
reg.comp15
Mean score
1.0
0.5
0.0
−0.5
−1.0
Significant In % Samples
0.25
0.50
0.75
1.00

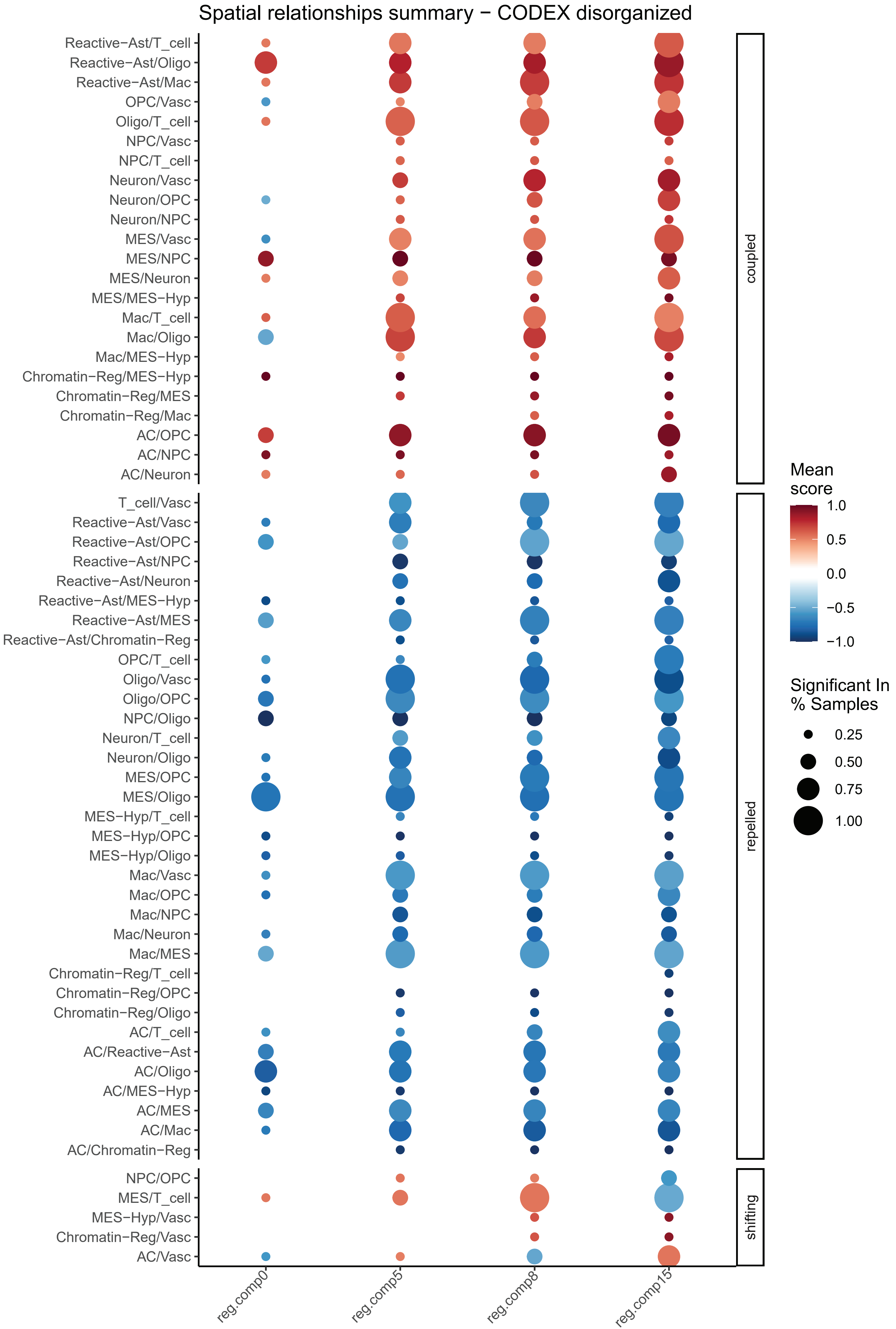

Spatial relationships summary − CODEX disorganized
Reactive−Ast/T_cell
Reactive−Ast/Oligo
Reactive−Ast/Mac
OPC/Vasc
Oligo/T_cell
NPC/Vasc
NPC/T_cell
Neuron/Vasc
Neuron/OPC
Neuron/NPC
MES/Vasc
MES/NPC
MES/Neuron
MES/MES−Hyp
Mac/T_cell
Mac/Oligo
Mac/MES−Hyp
Chromatin−Reg/MES−Hyp
Chromatin−Reg/MES
Chromatin−Reg/Mac
AC/OPC
AC/NPC
AC/Neuron
T_cell/Vasc
Reactive−Ast/Vasc
Reactive−Ast/OPC
Reactive−Ast/NPC
Reactive−Ast/Neuron
Reactive−Ast/MES−Hyp
Reactive−Ast/MES
Reactive−Ast/Chromatin−Reg
OPC/T_cell
Oligo/Vasc
Oligo/OPC
NPC/Oligo
Neuron/T_cell
Neuron/Oligo
MES/OPC
MES/Oligo
MES−Hyp/T_cell
MES−Hyp/OPC
MES−Hyp/Oligo
Mac/Vasc
Mac/OPC
Mac/NPC
Mac/Neuron
Mac/MES
Chromatin−Reg/T_cell
Chromatin−Reg/OPC
Chromatin−Reg/Oligo
AC/T_cell
AC/Reactive−Ast
AC/Oligo
AC/MES−Hyp
AC/MES
AC/Mac
AC/Chromatin−Reg
NPC/OPC
MES/T_cell
MES−Hyp/Vasc
Chromatin−Reg/Vasc
AC/Vasc
coupled
repelled
shifting
reg.comp0
reg.comp5
reg.comp8
reg.comp15
Mean score
1.0
0.5
0.0
−0.5
−1.0
Significant In % Samples
0.25
0.50
0.75
1.00

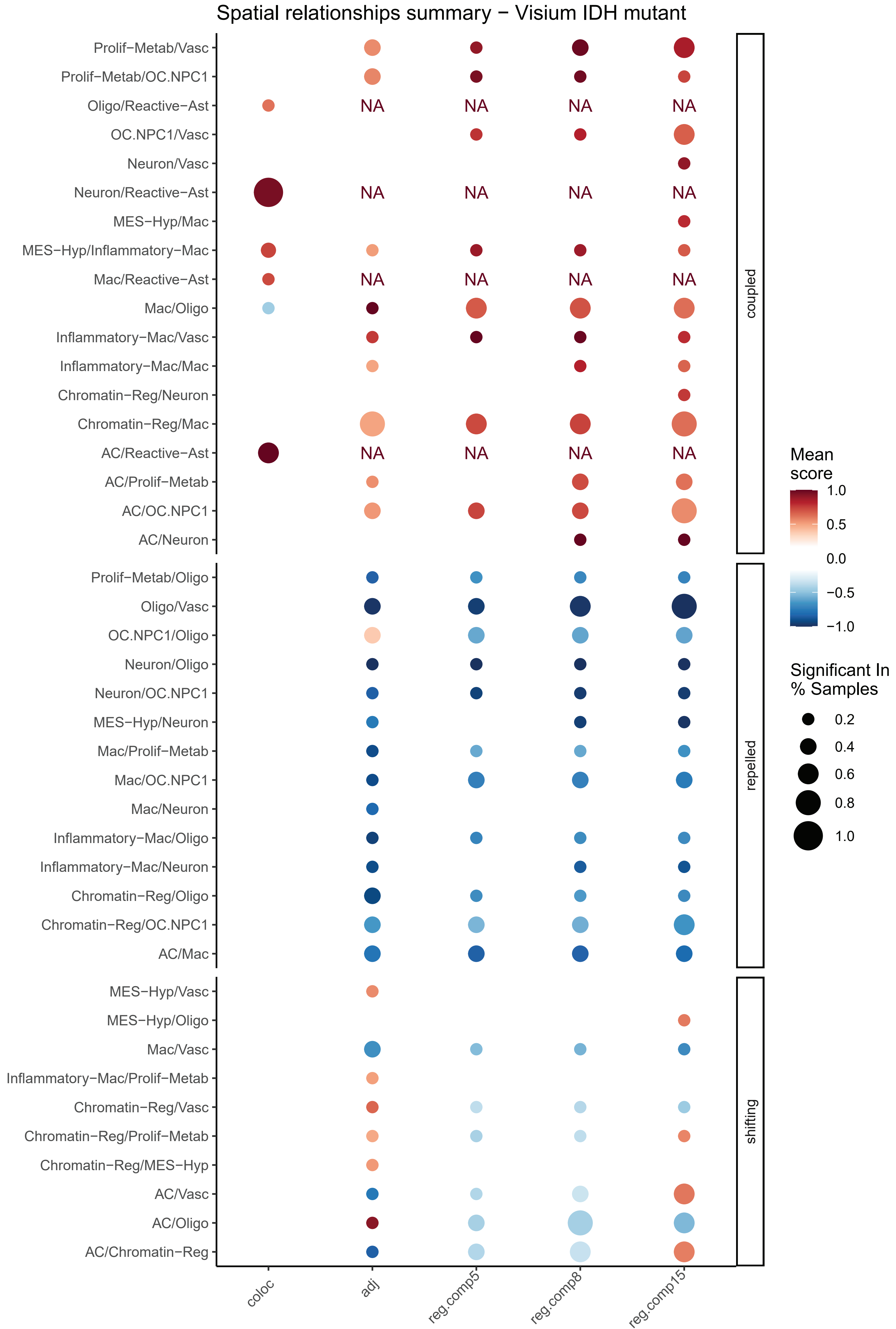
Spatial relationships summary − Visium IDH mutant
Prolif−Metab/Vasc
Prolif−Metab/OC.NPC1
Oligo/Reactive−Ast
OC.NPC1/Vasc
Neuron/Vasc
Neuron/Reactive−Ast
MES−Hyp/Mac
MES−Hyp/Inflammatory−Mac
Mac/Reactive−Ast
Mac/Oligo
Inflammatory−Mac/Vasc
Inflammatory−Mac/Mac
Chromatin−Reg/Neuron
Chromatin−Reg/Mac
AC/Reactive−Ast
AC/Prolif−Metab
AC/OC.NPC1
AC/Neuron
Prolif−Metab/Oligo
Oligo/Vasc
OC.NPC1/Oligo
Neuron/Oligo
Neuron/OC.NPC1
MES−Hyp/Neuron
Mac/Prolif−Metab
Mac/OC.NPC1
Mac/Neuron
Inflammatory−Mac/Oligo
Inflammatory−Mac/Neuron
Chromatin−Reg/Oligo
Chromatin−Reg/OC.NPC1
AC/Mac
MES−Hyp/Vasc
MES−Hyp/Oligo
Mac/Vasc
Inflammatory−Mac/Prolif−Metab
Chromatin−Reg/Vasc
Chromatin−Reg/Prolif−Metab
Chromatin−Reg/MES−Hyp
AC/Vasc
AC/Oligo
AC/Chromatin−Reg
NA
coupled
repelled
shifting
coloc
adj
reg.comp5
reg.comp8
reg.comp15
Mean score
1.0
0.5
0.0
−0.5
−1.0
Significant In % Samples
0.2
0.4
0.6
0.8
1.0

**Spatial relationships summary:** Summary heatmap of state-state associations across scales. Each column represents the summary of a different spatial relationship measure across GBM samples. Dots are colored by mean scaled relationship strength (Methods) and dot size corresponds to the fraction of samples in which the relationship is significant. State pairs are labeled as coupled, repelled or shifting by their mean score across all analyses. Panels title describe the platform (Visium or CODEX), the sample type (structured or disorganized) and glioma type (IDH-mutant or GBM if not specified) for which the data is shown.

#### Spatial relationship - adjacency - structured

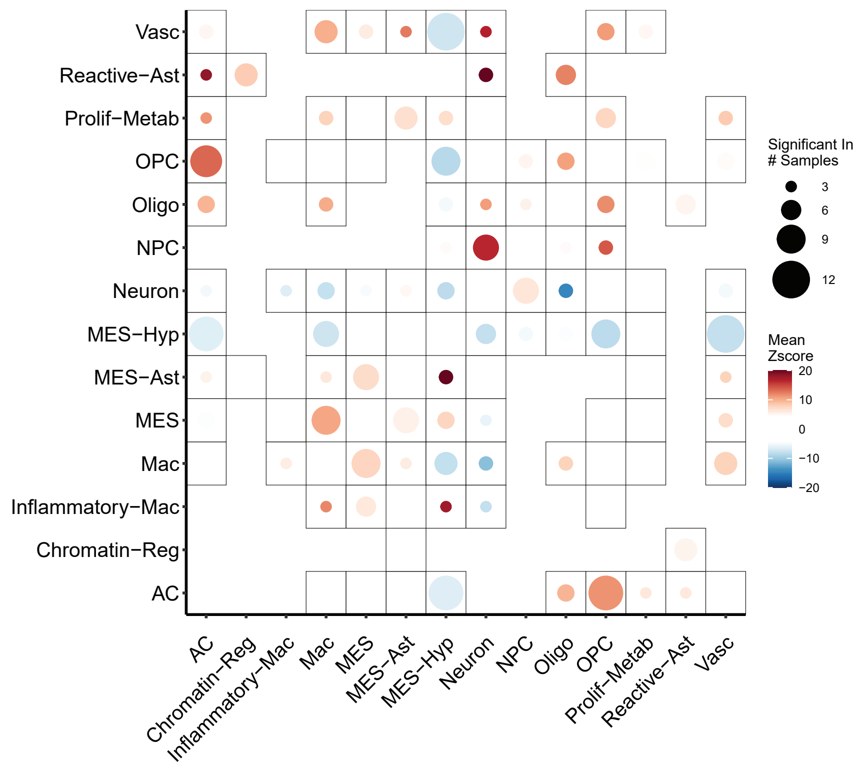

#### Spatial relationship - adjacency - disorganized

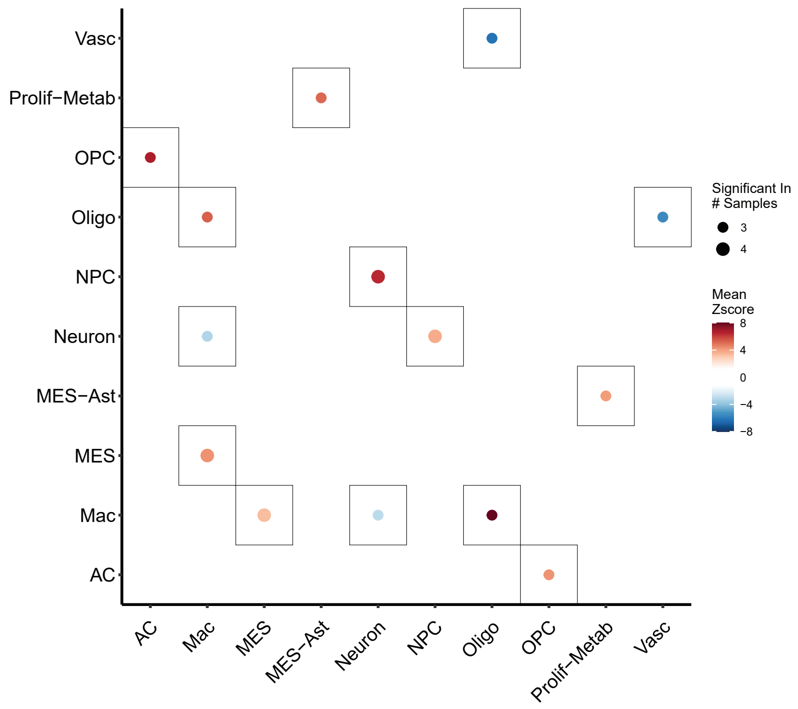

**Spatial relationship – adjacency:** Summary heatmap of state- state associations for adjacency analysis for Visium structured (upper panel) and disorganized (lower panel) samples. Row and columns corresponds to states. Dots are colored by mean relationship strength (Methods) and dot size corresponds to the fraction of samples in which the relationship is significant.

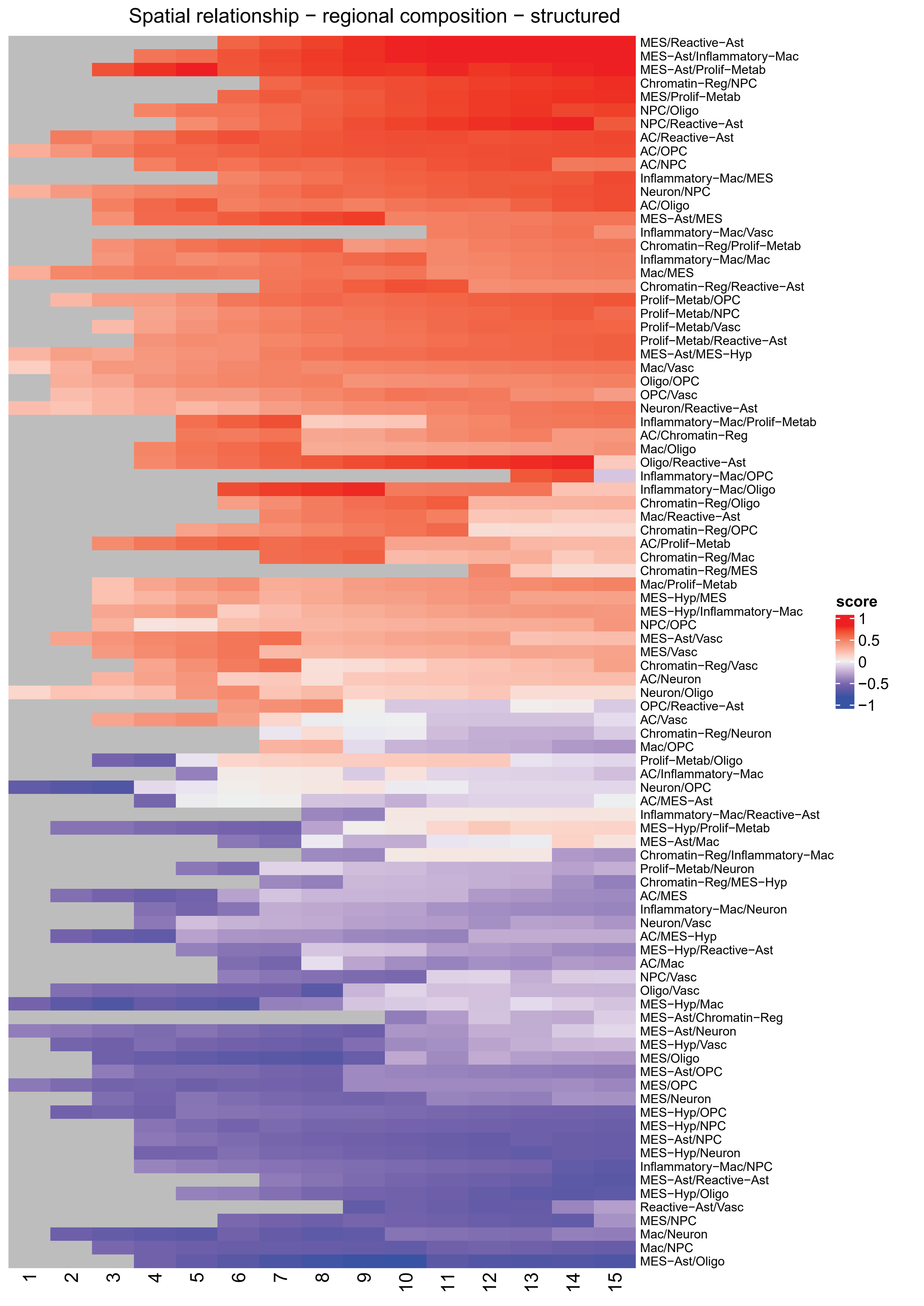
Spatial relationship − regional composition − structured
MES/Reactive−Ast
MES−Ast/Inflammatory−Mac
MES−Ast/Prolif−Metab
Chromatin−Reg/NPC
MES/Prolif−Metab
NPC/Oligo
NPC/Reactive−Ast
AC/Reactive−Ast
AC/OPC
AC/NPC
Inflammatory−Mac/MES
Neuron/NPC
AC/Oligo
MES−Ast/MES
Inflammatory−Mac/Vasc
Chromatin−Reg/Prolif−Metab
Inflammatory−Mac/Mac
Mac/MES
Chromatin−Reg/Reactive−Ast
Prolif−Metab/OPC
Prolif−Metab/NPC
Prolif−Metab/Vasc
Prolif−Metab/Reactive−Ast
MES−Ast/MES−Hyp
Mac/Vasc
Oligo/OPC
OPC/Vasc
Neuron/Reactive−Ast
Inflammatory−Mac/Prolif−Metab
AC/Chromatin−Reg
Mac/Oligo
Oligo/Reactive−Ast
Inflammatory−Mac/OPC
Inflammatory−Mac/Oligo
Chromatin−Reg/Oligo
Mac/Reactive−Ast
Chromatin−Reg/OPC
AC/Prolif−Metab
Chromatin−Reg/Mac
Chromatin−Reg/MES
Mac/Prolif−Metab
MES−Hyp/MES
MES−Hyp/Inflammatory−Mac
NPC/OPC
MES−Ast/Vasc
MES/Vasc
Chromatin−Reg/Vasc
AC/Neuron
Neuron/Oligo
OPC/Reactive−Ast
AC/Vasc
Chromatin−Reg/Neuron
Mac/OPC
Prolif−Metab/Oligo
AC/Inflammatory−Mac
Neuron/OPC
AC/MES−Ast
Inflammatory−Mac/Reactive−Ast
MES−Hyp/Prolif−Metab
MES−Ast/Mac
Chromatin−Reg/Inflammatory−Mac
Prolif−Metab/Neuron
Chromatin−Reg/MES−Hyp
AC/MES
Inflammatory−Mac/Neuron
Neuron/Vasc
AC/MES−Hyp
MES−Hyp/Reactive−Ast
AC/Mac
NPC/Vasc
Oligo/Vasc
MES−Hyp/Mac
MES−Ast/Chromatin−Reg
MES−Ast/Neuron
MES−Hyp/Vasc
MES/Oligo
MES−Ast/OPC
MES/OPC
MES/Neuron
MES−Hyp/OPC
MES−Hyp/NPC
MES−Ast/NPC
MES−Hyp/Neuron
Inflammatory−Mac/NPC
MES−Ast/Reactive−Ast
MES−Hyp/Oligo
Reactive−Ast/Vasc
MES/NPC
Mac/Neuron
Mac/NPC
MES−Ast/Oligo
score
1
0.5
0
−0.5
−1

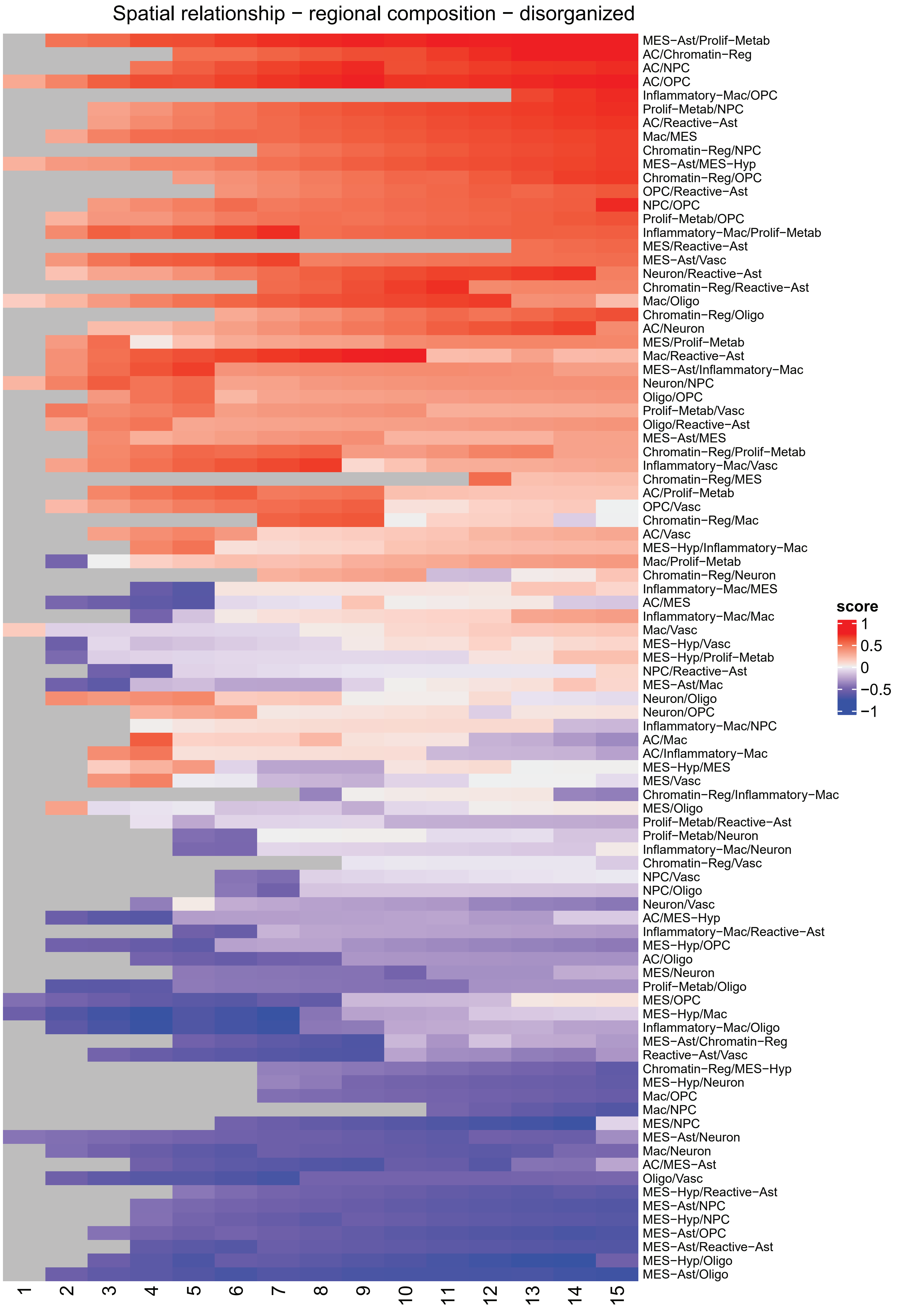
Spatial relationship − regional composition − disorganized
MES−Ast/Prolif−Metab
AC/Chromatin−Reg
AC/NPC
AC/OPC
Inflammatory−Mac/OPC
Prolif−Metab/NPC
AC/Reactive−Ast
Mac/MES
Chromatin−Reg/NPC
MES−Ast/MES−Hyp
Chromatin−Reg/OPC
OPC/Reactive−Ast
NPC/OPC
Prolif−Metab/OPC
Inflammatory−Mac/Prolif−Metab
MES/Reactive−Ast
MES−Ast/Vasc
Neuron/Reactive−Ast
Chromatin−Reg/Reactive−Ast
Mac/Oligo
Chromatin−Reg/Oligo
AC/Neuron
MES/Prolif−Metab
Mac/Reactive−Ast
MES−Ast/Inflammatory−Mac
Neuron/NPC
Oligo/OPC
Prolif−Metab/Vasc
Oligo/Reactive−Ast
MES−Ast/MES
Chromatin−Reg/Prolif−Metab
Inflammatory−Mac/Vasc
Chromatin−Reg/MES
AC/Prolif−Metab
OPC/Vasc
Chromatin−Reg/Mac
AC/Vasc
MES−Hyp/Inflammatory−Mac
Mac/Prolif−Metab
Chromatin−Reg/Neuron
Inflammatory−Mac/MES
AC/MES
Inflammatory−Mac/Mac
Mac/Vasc
MES−Hyp/Vasc
MES−Hyp/Prolif−Metab
NPC/Reactive−Ast
MES−Ast/Mac
Neuron/Oligo
Neuron/OPC
Inflammatory−Mac/NPC
AC/Mac
AC/Inflammatory−Mac
MES−Hyp/MES
MES/Vasc
Chromatin−Reg/Inflammatory−Mac
MES/Oligo
Prolif−Metab/Reactive−Ast
Prolif−Metab/Neuron
Inflammatory−Mac/Neuron
Chromatin−Reg/Vasc
NPC/Vasc
NPC/Oligo
Neuron/Vasc
AC/MES−Hyp
Inflammatory−Mac/Reactive−Ast
MES−Hyp/OPC
AC/Oligo
MES/Neuron
Prolif−Metab/Oligo
MES/OPC
MES−Hyp/Mac
Inflammatory−Mac/Oligo
MES−Ast/Chromatin−Reg
Reactive−Ast/Vasc
Chromatin−Reg/MES−Hyp
MES−Hyp/Neuron
Mac/OPC
Mac/NPC
MES/NPC
MES−Ast/Neuron
Mac/Neuron
AC/MES−Ast
Oligo/Vasc
MES−Hyp/Reactive−Ast
MES−Ast/NPC
MES−Hyp/NPC
MES−Ast/OPC
MES−Ast/Reactive−Ast
MES−Hyp/Oligo
MES−Ast/Oligo
score
1
0.5
0
−0.5
−1

**Spatial relationship – regional composition:** Summary heatmap of state- state associations for regional composition analysis for Visium structured (upper panel) and disorganized (lower panel) samples. Rows correspond to different pairs of states while columns correspond to different radii ranging from 1 to 15. Cells color reflect the regional composition correlation score of each pair for a given window. Grey cells correspond to pairs' score that were not significant across samples (Methods).

#### Supplementary Note 2

| Marker | IF | CODEX |
|---|---|---|
| CD45 | NA | 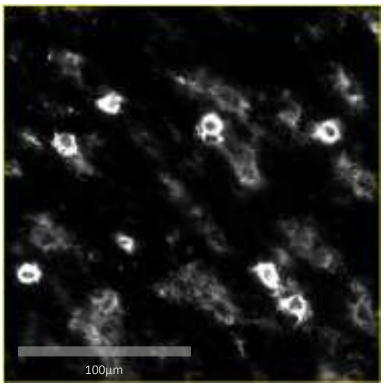  |
| CD3 | NA | 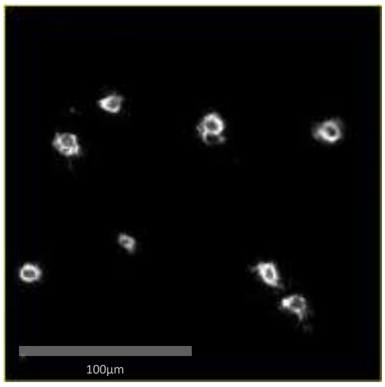  |
| CD69 | NA | 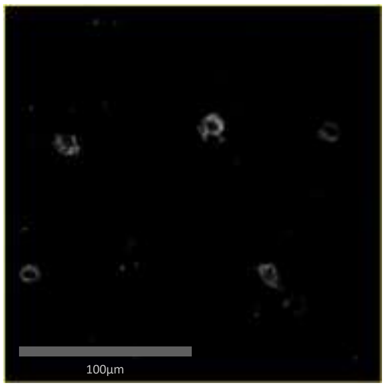  |
| CD90 | NA | 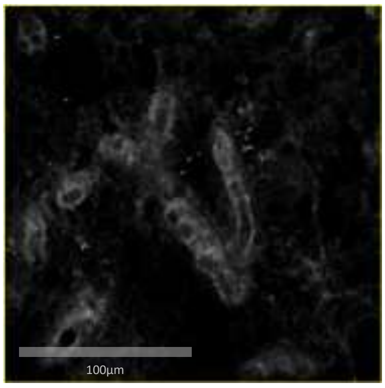  |
| CD11c | NA | 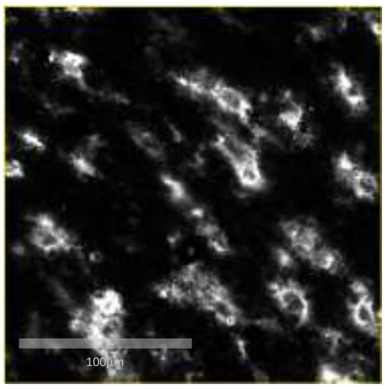  |

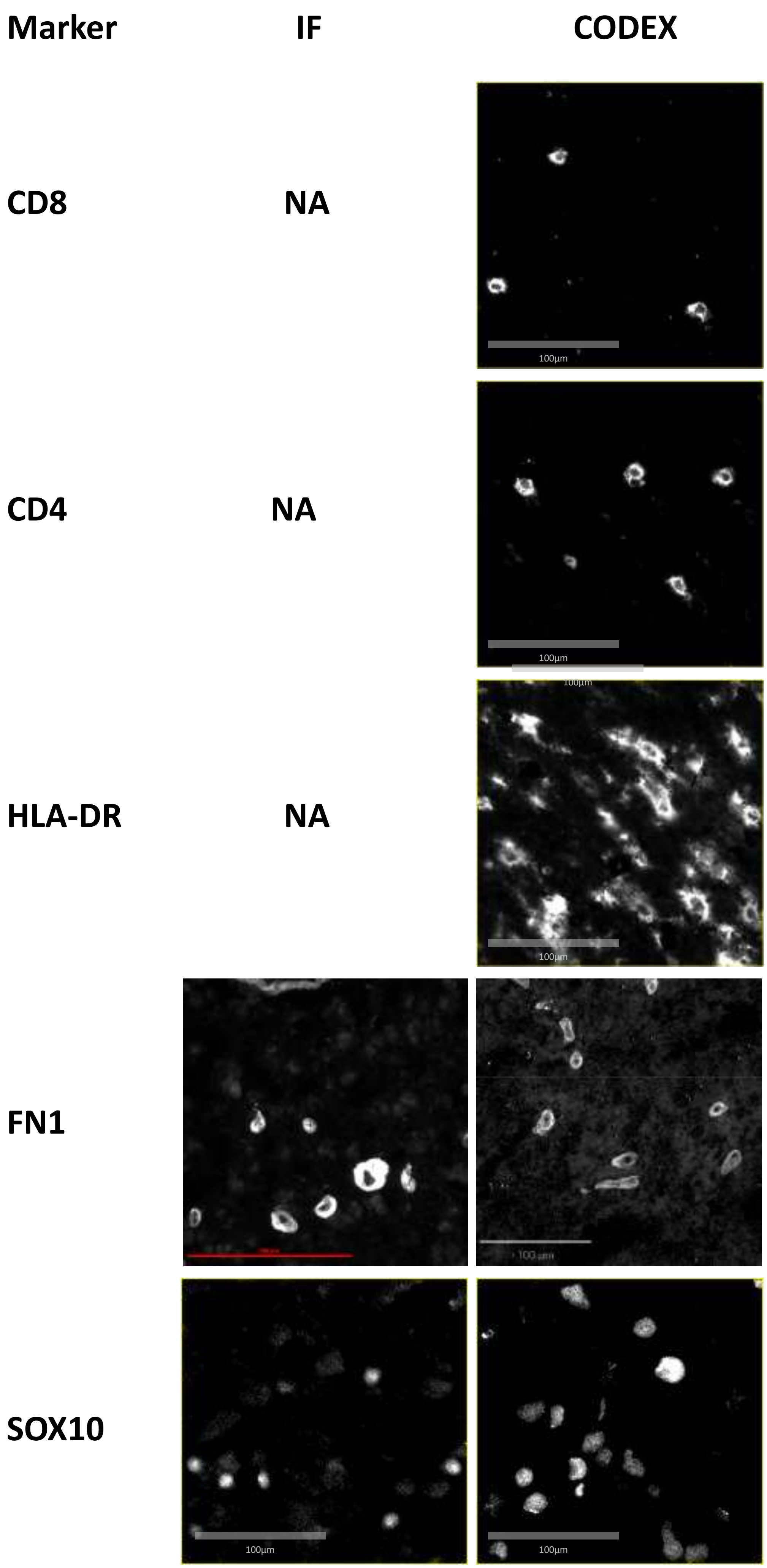
Marker
IF
CODEX
CD8
NA
CD4
NA
HLA-DR
NA
FN1
SOX10
100µm

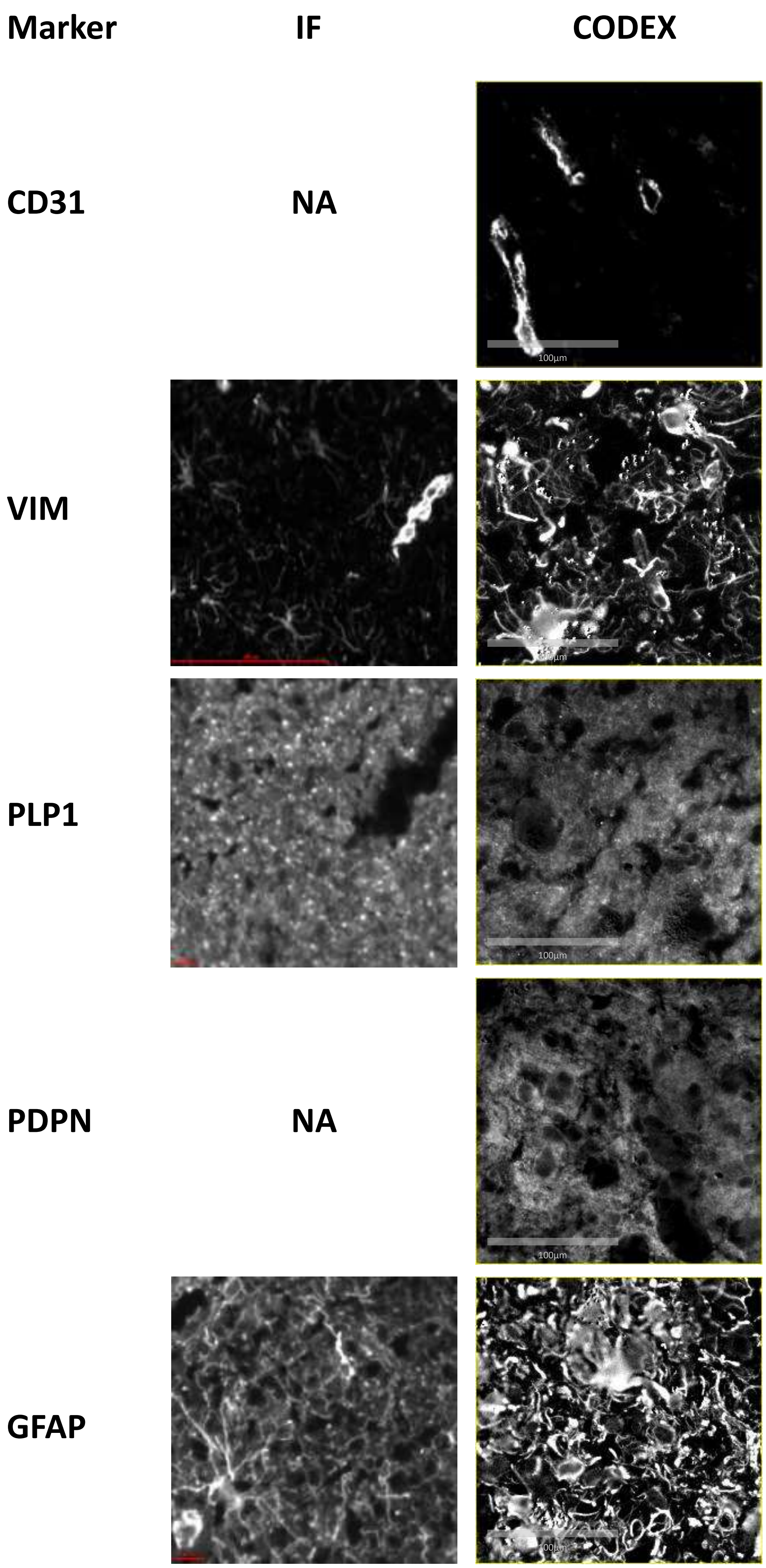
Marker
IF
CODEX
CD31
NA
VIM
PLP1
PDPN
NA
GFAP

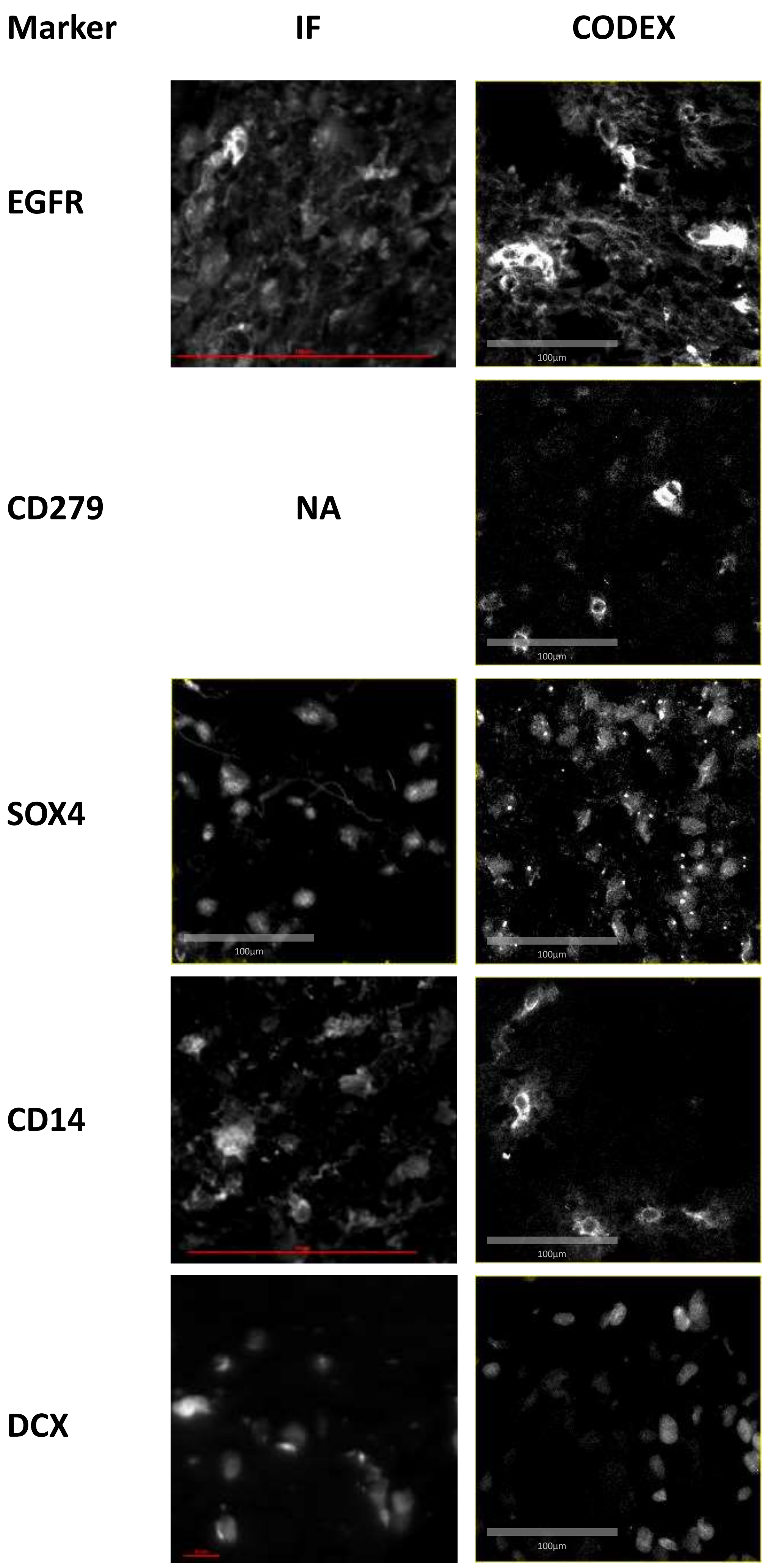
Marker
IF
CODEX
EGFR
100µm
CD279
NA
100µm
SOX4
100µm
100µm
CD14
100µm
DCX
100µm

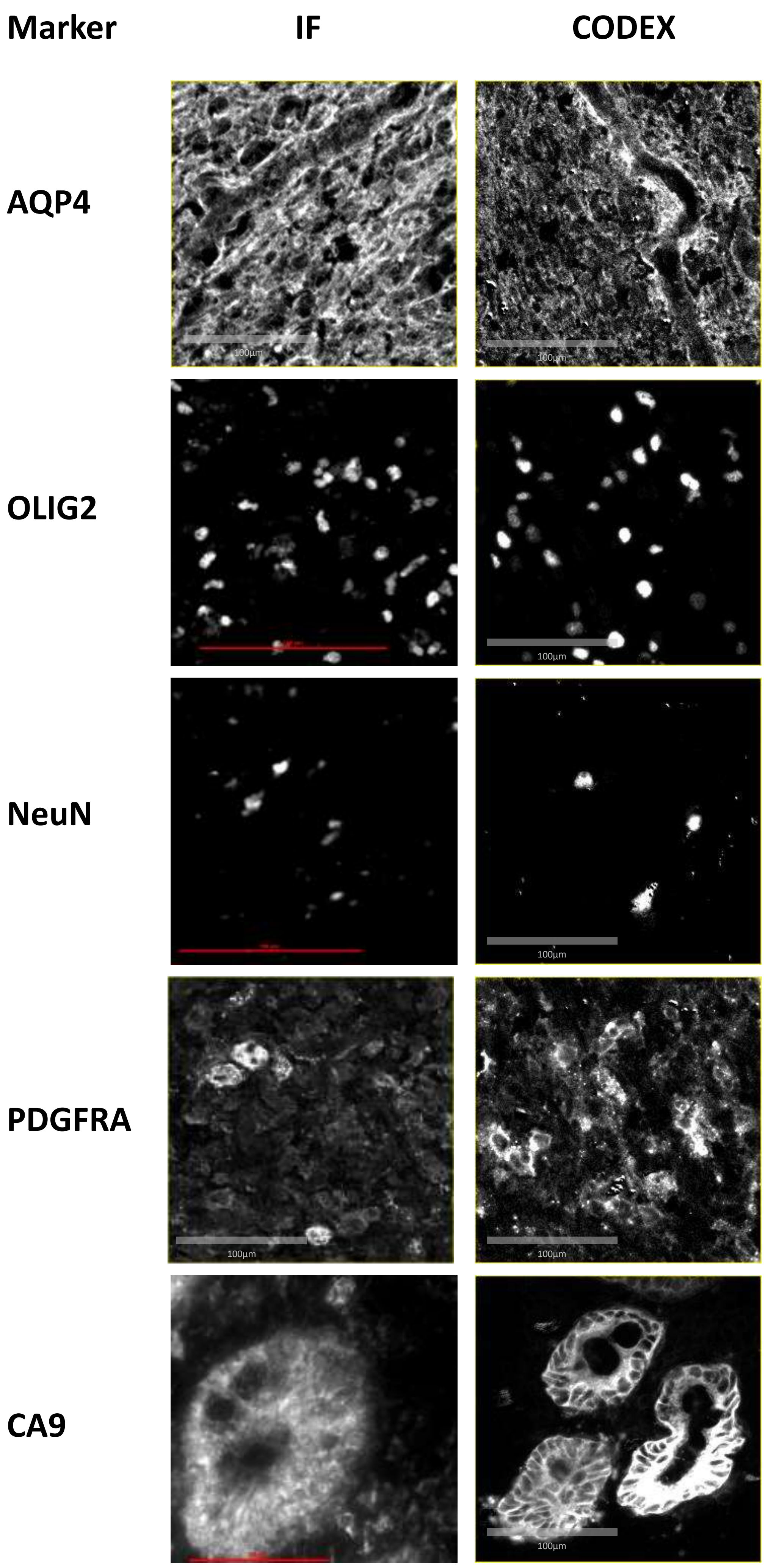
Marker
IF
CODEX
AQP4
OLIG2
NeuN
PDGFRA
CA9
100µm

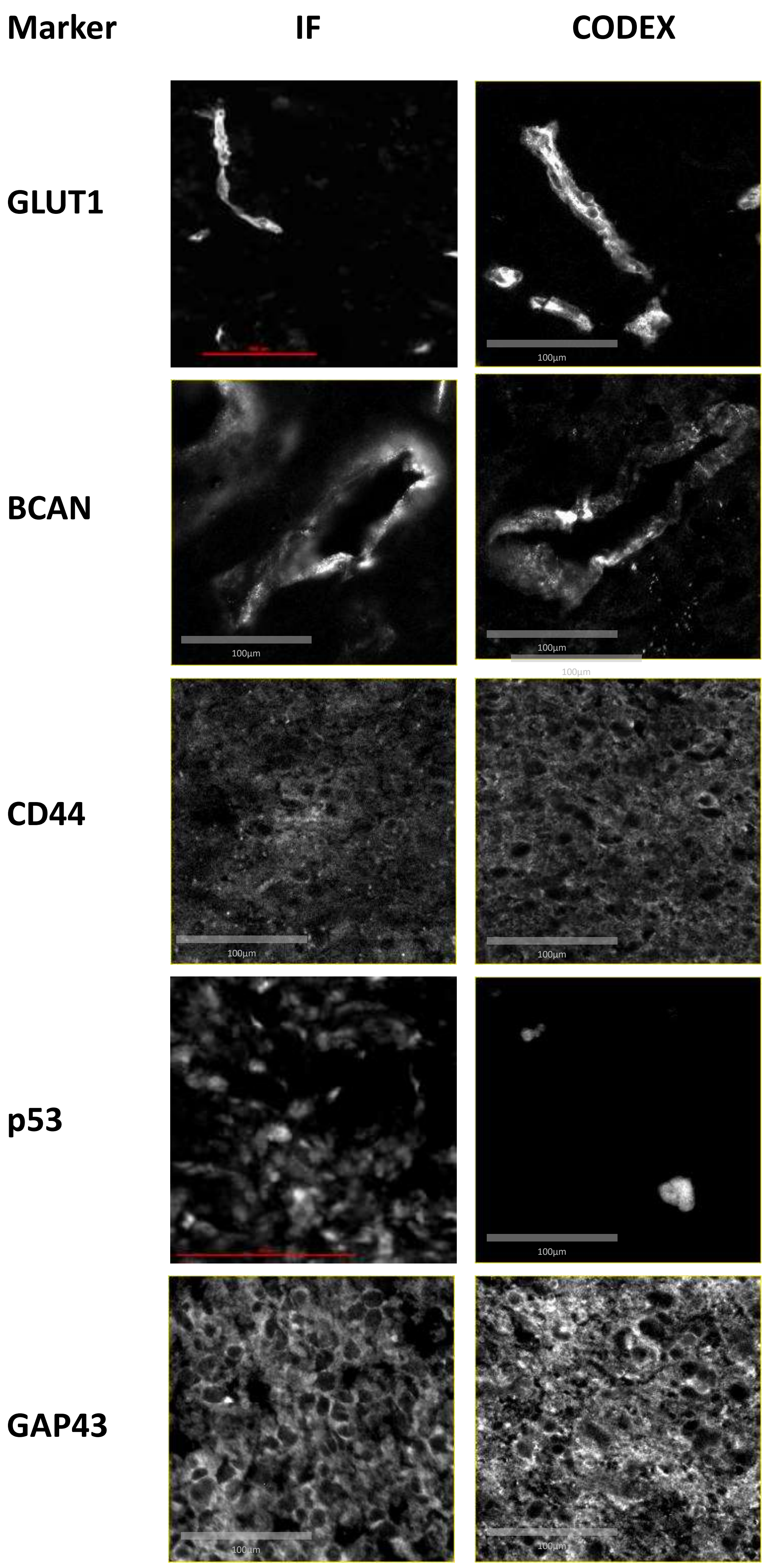
Marker
IF
CODEX
GLUT1
BCAN
CD44
p53
GAP43
100µm

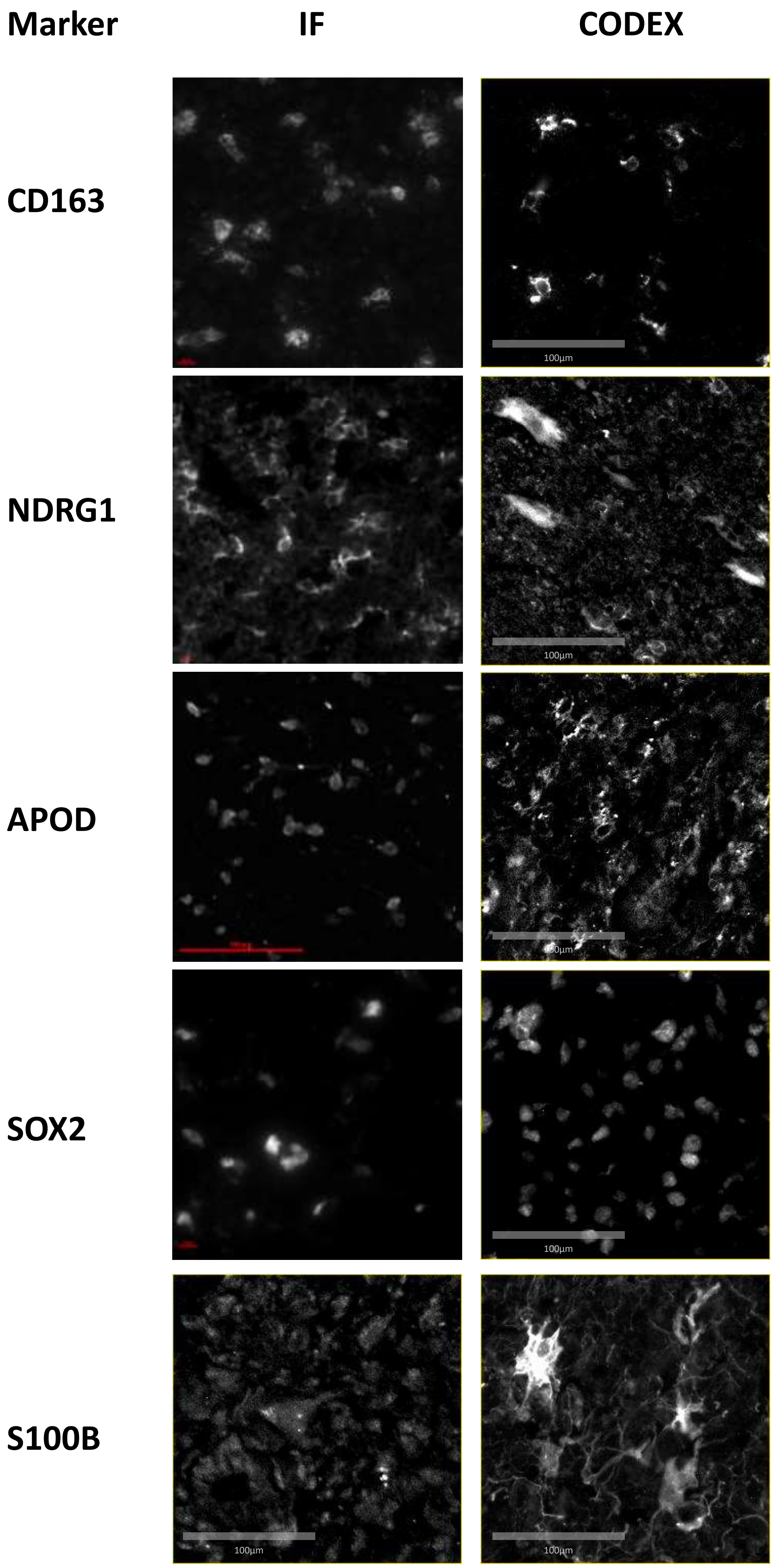
Marker
IF
CODEX
CD163
NDRG1
APOD
SOX2
S100B
100µm

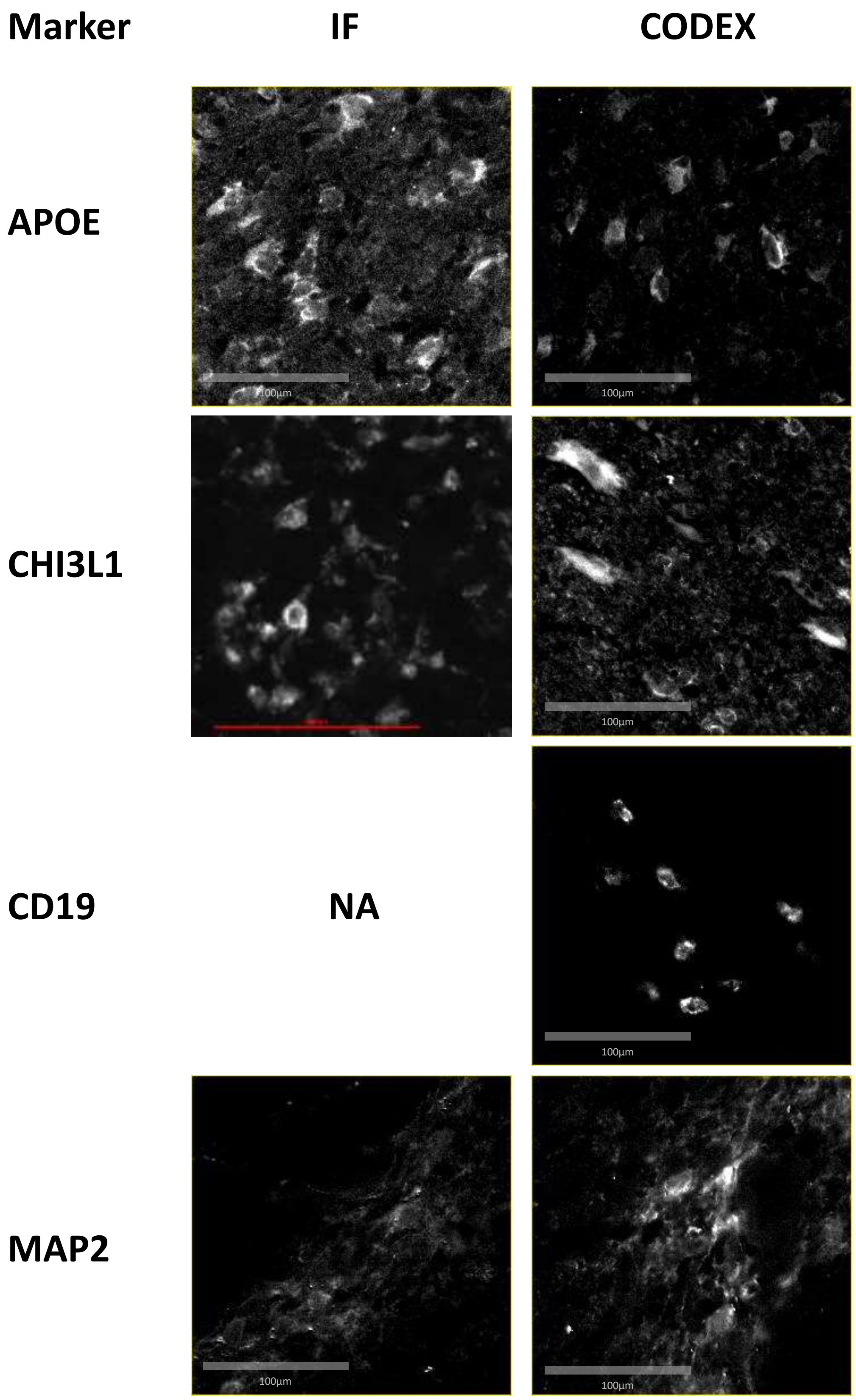

**Supplementary Note 2: Antibody validations.**
Images of antibody validation experiments using conventional immunofluorescence (IF) or CODEX for the indicated markers. Magnifications as indicated by scale bars. Antibodies pre-conjugated by Akoya were not validate with conventional IF (NA).
